# Computing the Human Interactome

**DOI:** 10.1101/2024.10.01.615885

**Authors:** Jing Zhang, Ian R. Humphreys, Jimin Pei, Jinuk Kim, Chulwon Choi, Rongqing Yuan, Jesse Durham, Siqi Liu, Hee-Jung Choi, Minkyung Baek, David Baker, Qian Cong

## Abstract

Protein-protein interactions (PPI) are essential for biological function. Recent advances in coevolutionary analysis and Deep Learning (DL) based protein structure prediction have enabled comprehensive PPI identification in bacterial and yeast proteomes, but these approaches have limited success to date for the more complex human proteome. Here, we overcome this challenge by 1) enhancing the coevolutionary signals with 7-fold deeper multiple sequence alignments harvested from 30 petabytes of unassembled genomic data, and 2) developing a new DL network trained on augmented datasets of domain-domain interactions from 200 million predicted protein structures. These advancements allow us to systematically screen through 200 million human protein pairs and predict 18,316 PPIs with an expected precision of 90%, among which 5,578 are novel predictions. 3D models of these predicted PPIs nearly triple the number of human PPIs with accurate structural information, providing numerous insights into protein function and mechanisms of human diseases.

## Main text

Detecting the interacting partners of proteins and determining the 3D structures of protein complexes are essential to understanding protein function (*1*–*3*). Large-scale experimental methods such as yeast two-hybrid (Y2H) and affinity-purification mass spectrometry (APMS) have been used to identify protein-protein interactions (PPI) across the human proteome (*4*–*7*). While powerful, these methods detect PPIs in non-physiological conditions and are associated with considerable false-positive and false-negative rates (*8, 9*). This is illustrated by the fact that although the human interactome is estimated to contain 74,000 to 200,000 PPIs (*10*), the experimental data in various PPI databases, UniProt (*11*), BioGrid (*12*), and STRING (*13*), suggest over 1 million (M) human PPIs may exist. Additionally, high-confidence PPIs from these databases show low consistency (**Fig. 1A**), with only 3988 PPIs regarded as confident by all three.

**Figure 1.**
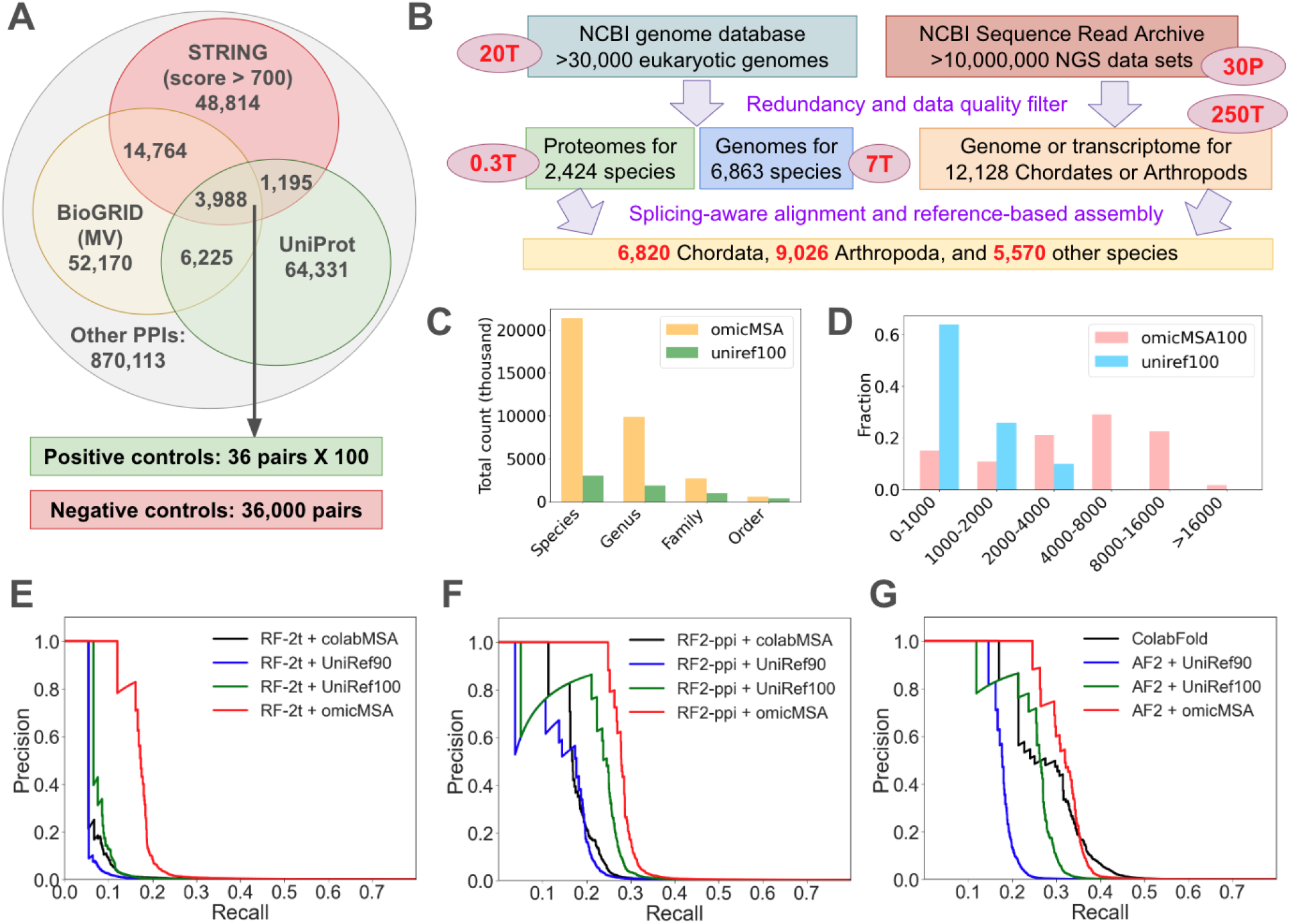
OmicMSA increases PPI identification accuracy. **(A)** Intersections of confident PPIs from different databases and our strategy to select the benchmark sets. **(B)** Strategies to assemble protein sequences from genomic data at NCBI. The boxes show statistics of different datasets, and the ovals by the boxes indicate their volume (T: terabytes, P: petabytes). Texts in purple by the arrows describe the data processing procedures. **(C)** Taxonomic diversity of our MSAs derived from genomic data and eukaryotic protein sequences in UniRef100. **(D)** Distribution of pMSA depth (filtered at 100% identity) for protein pairs in the benchmark sets. **(E-G)** Precision and recall curves (averaged over the 100 positive control sets) of different methods in distinguishing the positive from the negative controls (signal:noise = **1:1000**). OmicMSA consistently improves the performance of different methods, such as **(E)** RF-2t (*23*), **(F)** RF2-ppi, and **(G)** AF2.

Computational approaches have been developed to complement experimental studies in resolving the human interactome. They predict PPIs based on homology to known interacting partners, quality of the predicted interfaces, and functional associations (*14*–*21*). Coevolutionary analysis (residues at PPI interfaces must coevolve to maintain their interactions) has been combined with 3D structure prediction methods, such as AlphaFold (AF) and RoseTTAFold (RF), to identify PPIs on a proteome-wide scale in bacteria and yeast (*22, 23*). The probability for any protein pair to interact is estimated by 1) creating multiple sequence alignments (MSAs) of orthologs of the two proteins in many species, 2) concatenating sequences of the same species to build paired MSAs (pMSAs), and 3) determining the probabilities for residues in the first protein to interact with residues in the second based on coevolution and complex structure modeling (*24*). While similar approaches have been used to detect interactions between selected human proteins (*18, 25*–*27*), a *de novo* proteome-wide PPI screen in humans is challenging due to the daunting computational scale, and the small number of high eukaryotic genomes that limits the statistical power of coevolution between human proteins.

We set out to overcome these challenges and systematically identify PPIs in humans. To enhance the statistical power of coevolutionary analysis, we reasoned that it should be possible to harness the petabytes of untapped genomic sequence data in draft eukaryotic genomes and genomic reads. To analyze hundreds of millions human protein pairs efficiently, we sought to develop a fast but accurate Deep Learning (DL) network for PPI prediction by 1) optimizing the architecture of state-of-the-art DL networks and 2) augmenting the training set of experimentally determined PPIs in the PDB with a much larger distillation set of domain-domain interactions (DDIs) harvested from accurately predicted monomer structures in the AlphaFold protein structure Database (AFDB) (*28*), based on the assumption that DDI interfaces should resemble PPI interfaces in coevolutionary and physicochemical properties.

### Harnessing untapped genomic sequence data

Biologists typically rely on databases with protein sequences annotated from genomes (such as Uniref) or metagenomic assemblies (such as BFD (*29*) and MGnify (*30*)). However, the majority of available eukaryotic genomes, especially for higher Eukaryotes, have not been annotated with protein sequences. Identifying protein-coding regions in eukaryotic genomes is non-trivial due to the complicated and frequent lineage-specific rules of translation initiation and mRNA splicing (*31*). Hence, it is not a routine practice to annotate and deposit the protein-coding sequences for draft eukaryotic genomes: among the 36,840 eukaryotic genomes available at NCBI in June, 2024, only 7,355 (20%) were associated with annotated proteins. Moreover, assembling diploid, heterozygous, and repeat-rich eukaryotic genomes is challenging without long-read sequencing (*32*). The NCBI Sequence Read Archive (SRA) database (*33*) has over 30 petabytes of shotgun sequencing reads, which have not been assembled into contigs or analyzed to predict the proteins they encode.

To enrich the coevolutionary information in our MSAs, we mined the NCBI genome and SRA databases to extract whole genome and whole transcriptome datasets for 21,414 diverse Eukaryotes, focusing on Chordates and Arthropods, two large phylums of higher Eukaryotes. We selected one representative genomic dataset per species, including 1) 2,424 species whose proteome sequences were previously annotated from genomes, 2) 6,863 species with draft genomes but no protein annotations, and 3) 12,128 species with whole genome or whole transcriptome shotgun reads. We developed a bioinformatic pipeline (see **supplemental Methods M4**) to assemble protein-coding sequences from each type of dataset, utilizing splicing-site-aware sequence aligners (*34, 35*) and reference protein sets from model organisms (**Fig. 1B**). All predicted protein sequences were aligned to their human orthologs, if available, and we used reciprocal best-hit criteria (*36*) to distinguish orthologs from paralogs. We name the resulting alignments omicMSAs, because they are directly derived from genomic data. Our omicMSAs include sequences from 21,414 species spanning 9,905 genera, 2,727 families, and 626 orders, significantly expanding the taxonomic diversity of available protein sequences in UniRef100, which only contains 3,082 species (**Fig. 1C**).

We reasoned that the rich evolutionary information in omicMSAs should improve the ability of DL networks to distinguish true PPIs from false ones. We tested this hypothesis using benchmarks derived from PPI databases. There are 75,739, 77,147, and 68,761 confident human PPIs from UniRef, BioGrid (multi-validated), and STRING (physical interactions with combined score > 700), respectively. However, only 3,988 (5.3%–5.8%) PPIs are shared by all (**Fig. 1A**), from which we selected 100 non-overlapping positive control sets, each containing 36 PPIs. The negative control set contains 36,000 random pairs of human proteins showing no evidence for their interactions, and its much larger size than the positive control sets allows performance evaluation at a low signal-to-noise ratio (1:1000) that approximates the situation of proteome-wide PPI screen (74k–200k true PPIs out of 200M pairs). We compared omicMSAs against MSAs built by other commonly used strategies. The first is HHblits (*37*) against UniRef, a widely used approach to prepare MSA inputs for RF2 and AF2. HHblits by default filters the output MSAs at 90% sequence identity, and we named this strategy UniRef90. Filtering MSAs before building pMSAs significantly reduces the number of taxa that can be paired; to increase pMSA depth, we disabled this filter and termed the alternative strategy UniRef100.

Metagenomic sequences are frequently used to obtain deeper MSAs and improve structure prediction (*38*). We used the ColabFold MSA pipeline (*39*) to combine UniRef and metagenomic sequences.

We found that omicMSA led to the best performance for all the tested methods (**supplemental fig S24**), including RF-2t used in yeast PPI screen (*23*), the newly developed RF2-ppi in this study, and AF2, in distinguishing true PPIs from random pairs (red curves in **Fig. 1E-G**). The ColabFold MSA (black curves in **Fig. 1E-G**) improves over UniRef100 when used with AF2, but it did not perform better with other tools. Metagenomic data, mostly sequenced from environmental samples, do not contain sequences of higher Eukaryotes and are not annotated with the assumption of mRNA splicing (*40*); thus, they do not necessarily improve human PPI prediction. In contrast, we focused on assembling sequences from genomic data of higher Eukaryotes with pipelines aware of mRNA splicing. All the tools we tested were trained using MSAs built from UniRef and metagenomic sequences, and they are not adapted to work with omicMSA. Thus, the superior performance of omicMSA demonstrates the power of stronger evolutionary signals gleaned from a much wider range of taxa. The available eukaryotic genomic data is constantly growing (∼20% more species per year), providing even greater potential in the future.

### RoseTTAFold2-PPI: a DL network for rapid PPI identification

Because the physicochemical properties of residue-residue contacts at PPI interfaces are expected to differ from contacts within tightly packed protein monomers (*2, 3*), AlphaFold-multimer (AFmm) (*41*) was trained with PPIs in the PDB to improve its ability to model 3D structures of protein complexes. However, such training may not improve an DL network’s ability on the distinct task of distinguishing true PPIs from false ones — instead, this might bias the network to predict PPIs between random pairs without strong coevolutionary signals. Indeed, AFmm does not perform well in distinguishing true PPIs from random pairs at a signal-to-noise ratio of 1:1000 (**Fig. 2D**). Trained on PPIs from the PDB and optimized for speed, we developed RF2-lite (*22*) to distinguish true PPIs from random pairs. RF2-lite is 20 times faster than AF2, but shows considerably lower accuracy than AF2 (**Fig. 2D**). We reason that one factor limiting the accuracy of current methods is the amount of training data. At 30% sequence identity, there are 24,358 clusters of PPIs (hetero-oligomers only) in the PDB (December, 2023), among which many pairs are human proteins or their homologs (MMseqs e-value < 0.00001); after removing these complexes to avoid information leakage for human PPI prediction, only 13,231 clusters remain (**Fig. 2B left**).

**Figure 2.**
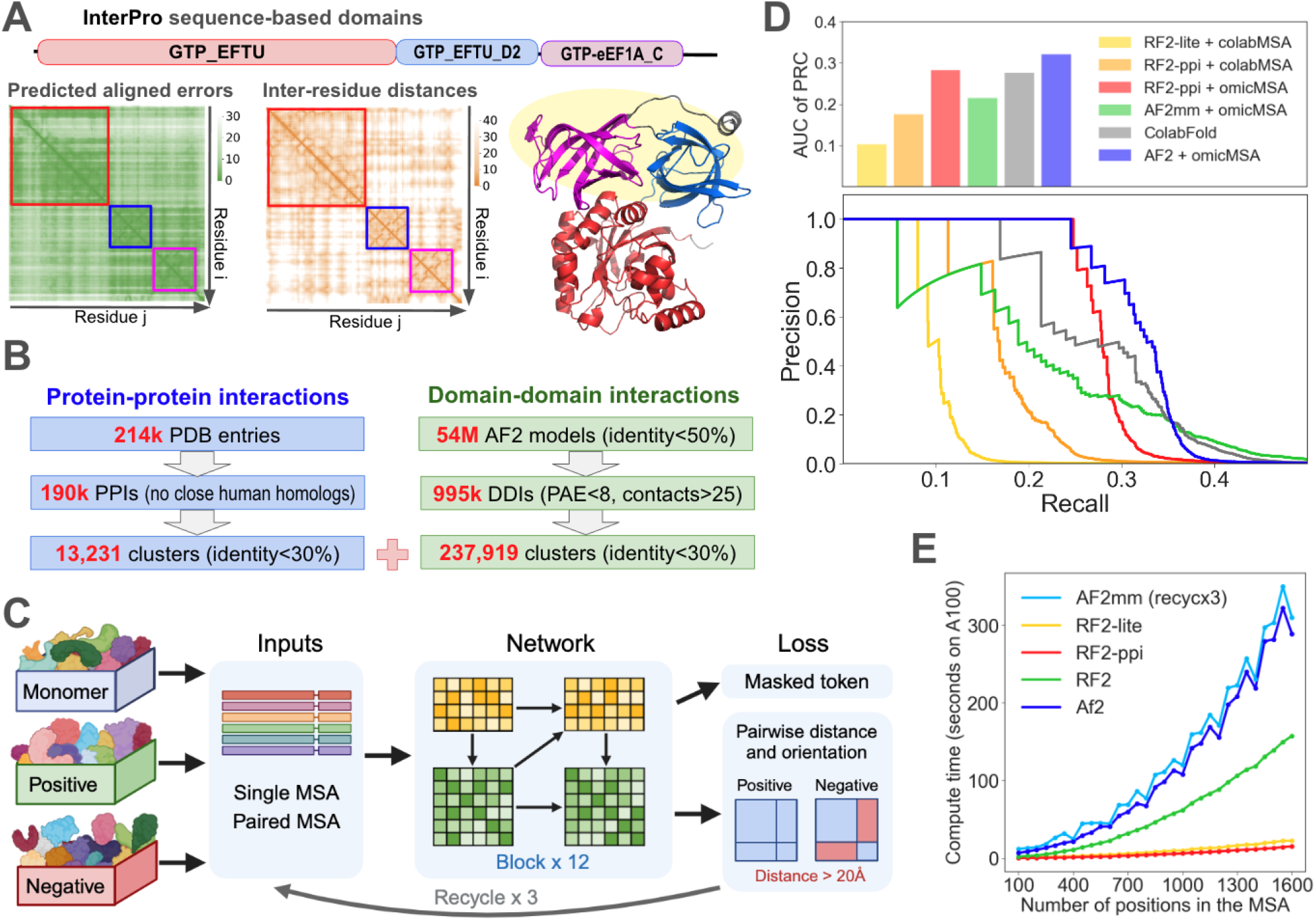
A DL network for PPI prediction, trained using datasets augmented with predicted DDIs. **(A)** Illustration of our domain segmentation protocol. Top: domains annotated by InterPro; left: PAE matrix; middle: inter-residue distance matrix; right: 3D structure. In both matrices, the parsed domains are marked by boxes colored the same as in the 3D structure. The two domains in yellow shade belong to the DDI training set; their interaction with the 3rd domain did not pass our inter-domain PAE cutoff. **(B)** Statistics of the PPI and DDI training datasets. **(C)** Architecture and training routine for RF2-ppi. **(D)** Precision and recall curves (below) and the area under the curves (above) for different methods in their ability to distinguish true PPIs from false ones (signal: noise = 1:1000). **(E)** Compute time as a function of input protein size for different methods.

We hypothesized that extracted DDIs from the 200M AF2 models in the AFDB (*42*) could significantly enlarge the training dataset for PPI predictors and improve their performance. DDIs within the same protein should resemble PPIs (*43, 44*). Domains are structural and evolutionary units that are recombined to create new proteins throughout evolution (*45*), and the interfaces between domains closely resemble those between distinct protein partners (*43, 44*). We previously developed a method to segment AF2 models into domains based on structural features (inter-residue distances and predicted aligned errors (PAEs)) and homology to previously classified domains from PDB entries (*46, 47*). We repurposed this method to integrate structural features and domain annotations of UniProt entries from InterPro (**Fig. 2A** and **supplemental Methods M2.2**). We found 12.4M multi-domain proteins among the 53.7M AFDB models filtered at 50% sequence identity (*48*). These models contain 22.6M high-quality (mean PAE within domains < 8) domain pairs from the same proteins. We focused on pairs with at least 25 inter-residue contacts (distance < 6Å) and high confidence in their interaction (mean inter-domain PAE < 8), resulting in 1M DDIs. Clustering these DDIs at 30% sequence identity resulted in 237,919 clusters (**Fig. 2B right**), 10-fold larger than the PPI training set.

We used both the PDB PPI and AFDB DDI sets to develop RF2-ppi (**Fig. 2C**) based on the RF2 architecture (*49*), which integrates the MSA, inter-residue, and 3D structure features through attention-based blocks. Additionally, we included monomeric 3D structures to ensure that RF2-ppi learns principles of protein folding; we added negative controls – random protein or domain pairs from the same organism with no evidence of being functionally related – to ensure that RF2-ppi learns to distinguish interacting partners from random pairs. The monomers, positive DDIs, negative DDIs, positive PPIs, and negative PPIs were combined at a 2:1:1:1:1 ratio to generate the training dataset. We explored a number of strategies to optimize the performance of RF2-ppi (see **supplemental Methods M3.2**); we found that adding the DDI training set and removing the 3D structure features and losses significantly improved its performance (**supplemental figs. S10 and S11**). The lightweight RF2-ppi performed better than RF2 for PPI identification (**supplemental fig. S15**), which again suggests that optimizing predicted 3D structures of protein complexes does not necessarily help to distinguish true PPIs from random pairs. This may be because weak human PPIs are frequently mediated by simple motifs (single helices or intrinsically disordered regions (IDRs)), while our training sets are dominated by interactions with larger interfaces. Direct integration of 3D features may drive the DL network to emphasize on the geometric and chemical complementarity at the PPI interfaces instead of the coevolutionary signals essential for predicting weak PPIs. The 3D structure module of AF2 is not directly integrated into the Evoformer, possibly allowing AF2 to properly balance coevolutionary signals and physicochemical properties.

We compared the performance of RF2-ppi with the ColabFold pipeline (skyblue in **Fig. 2D**) widely used by the scientific community (*39*) and other methods. Using area under the precision and recall curve (AUCPR) as a metric, RF2-ppi (orange in **Fig. 2D**) showed a 1.7-fold improvement over our previous method, RF2-lite (gold in **Fig. 2D**). When combined with omicMSAs, RF2-ppi (red in **Fig. 2D**) slightly outperforms the ColabFold pipeline in its ability to identify true PPIs and is 20 times faster than AF2/ColabFold (**Fig. 2E**). AFmm (green in **Fig. 2D**), despite its superior performance in modeling 3D structures of protein complexes, shows significantly worse performance than RF2-ppi in distinguishing true PPIs from random pairs. The inferior performance of AFmm is related to the low signal-to-noise ratio (1:1000) associated with *de novo* PPI screens; at a signal-to-noise ratio of 1:10, AFmm will outperform the other methods. The best performance is achieved when AF2 is used in combination with omicMSAs (deepblue in **Fig. 2D**). We used both RF2-ppi and AF2 to identify and model human PPIs: the former allows us to carry out our screen on a proteome-wide scale, while the latter is needed to obtain high-quality 3D structure models of confident PPIs.

### Proteome-wide identification of human PPIs

Equipped with omicMSA and RF2-ppi, we set out to comprehensively predict the human interactome. We took AF2 models for the human proteome (20,504 proteins) from the AFDB and identified domains based on structural compactness (*46*) and evolutionary conservation (*50*). We included these domains and relatively conserved residues in the inter-domain linkers (conservation > 25% quantile of residues in domains) in our screen; excluding regions that are poorly conserved and lack rigid 3D structures improves the performance of our PPI screen pipeline (**supplemental fig. S23**). To fit into limited GPU memory, we split large proteins into segments with few inter-residue contacts and flexible relative orientation (see **supplemental Methods M5.1**) between them. In total, we screened 191M pairs comprised of 19,528 proteins, excluding a small fraction (4.8%) of proteins due to protein size limits or the lack of compact structures and conserved motifs.

We began with an unbiased systematic search for PPIs across the 191M protein pairs. To make the search more tractable, we focused on pairs in the same cellular compartment (CC) based on keywords annotated by UniProt. We analyzed 53.8M pairs sharing CC annotations and 57.4M pairs involving proteins without CC annotations sequentially through Direct Coupling Analysis (DCA) (*50, 51*), RF2-ppi, and AF2 (**Fig. 3A**). Although DCA, a statistical method to detect coevolution, shows significantly worse performance than the state-of-the-art DL tools, it is 5-fold faster than RF2-ppi and we used it to select 43.6M (40%) pairs. Based on our benchmark, most (over 80%) pairs showing high interaction probability by AF2 will pass this DCA pre-filter (**supplemental table S4**). This *de novo* pipeline (see **Fig. 3B** for performance evaluation and cutoff at each step) predicted 6,966 PPIs with an expected precision of 90%. We complemented these results by carrying out a second screen incorporating the extensive information about human PPIs derived primarily from high throughput experiments. Evaluating such noisy experimental data with our *in silico* pipeline allows us to identify a smaller set of PPIs at high confidence. Prior evidence of interactions allowed us to use lower RF2-ppi and AF2 cutoffs while maintaining the same level of expected precision. We predicted 10,627 PPIs among the 5.23M genetically interacting pairs from STRING (**Fig. 3A**) and 12,832 PPIs from the 1.15M pairs with evidence for physical interactions from UniProt, BioGRID, and STRING. The *de novo* (unbiased) and evidence-guided pipelines allowed us to predict 18,307 PPIs at 90% precision and 30.8% recall (see **supplemental Methods M5.6**).

**Figure 3.**
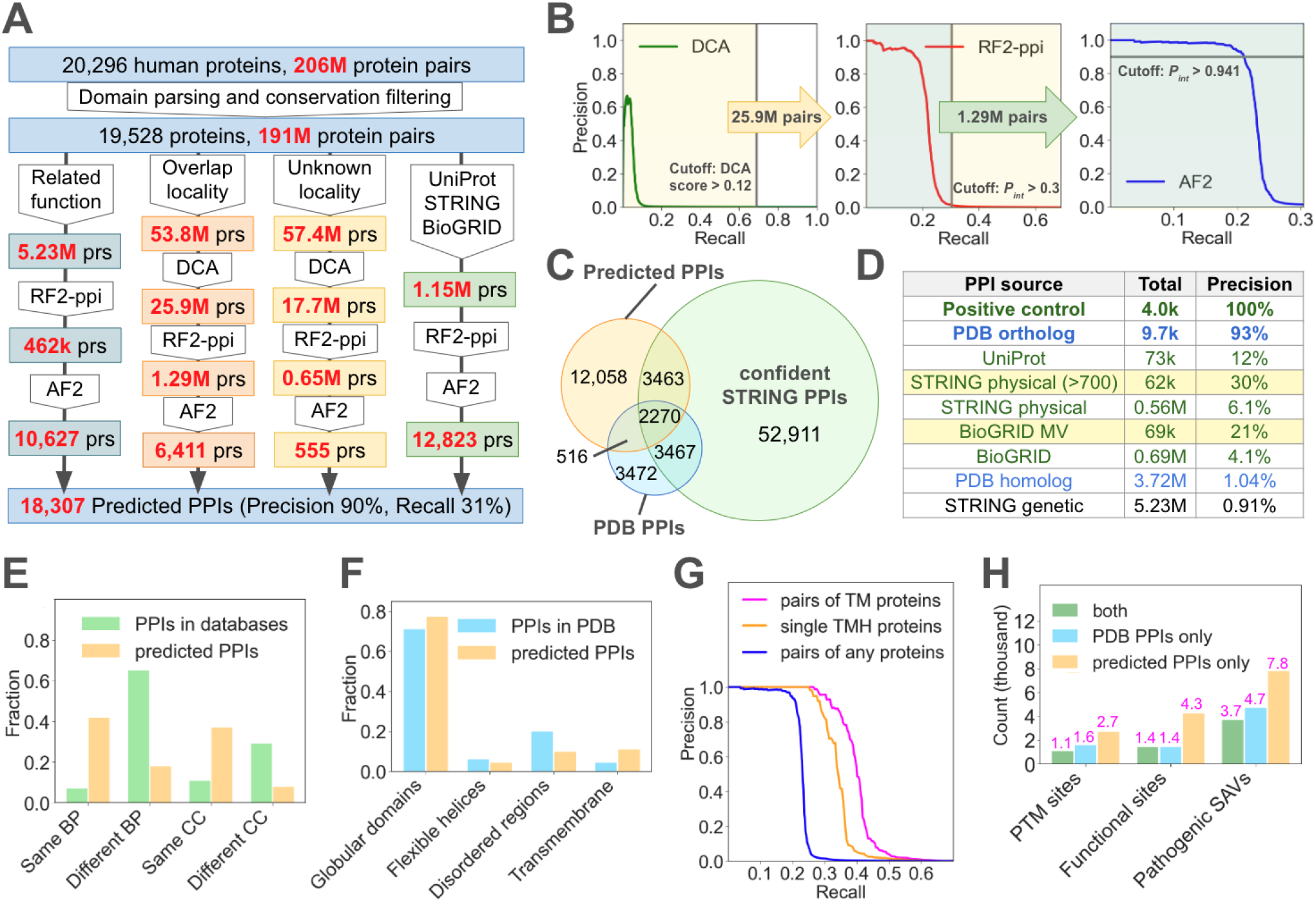
Human Proteome-wide PPI identification. **(A)** PPI screen strategies. Key methods are shown in white connectors and numbers of selected pairs after key steps are shown in colored squares. **(B)** Performance of our *de novo* pipeline applied to colocalized protein pairs. The grey line indicates the recall (vertical) or precision (horizontal) cutoff (corresponding score labeled by the line) at each stage; number of pairs selected at each stage is shown in the arrow. **(C)** A Venn diagram showing the overlap between our predictions and PPIs from other sources; such overlap are used to estimate **(D)** the precision of PPIs in other databases: PDB ortholog, with orthologous PDB templates; PDB homolog, with homologous PDB templates; yellow highlights: the confident subsets from STRING and BioGRID. **(E)** Predicted PPIs show a much higher chance to participate in the same biological processes (BP) and localize in the same cellular components (CC) than human PPIs in databases. Same: complete overlap of UniProt keywords; Different: no overlap; partial overlap not shown. **(F)** Fraction of different interface types in predicted and experimentally determined PPIs. Globular domains: >50% interface residues for both; flexible helices: >50% interface residues for one; disordered regions: >50% interface residues for one; transmembrane: >25% interface residues for one. **(G)** Performance of our *de novo* pipeline for TMPs. **(H)** Numbers (labeled on top) of functional sites at PPI interfaces.

We compared our predictions against PPIs from other databases (**Fig. 3C**). We reasoned that the fraction of PPIs from another source that can be identified by our *in silico* screen (*f*_*AI/DB*_) reflects the precision of PPIs in that database, which can be computed as *pre*_*DB*_ = *f*_*AI/DB*_ */ rec*_*AI*_, *rec*_*AI*_ = 0. 308. PPIs with orthologous PDB templates are mostly (93%) true. However, not all contacting chains in PDB entries interact in physiological conditions; some instead result from experimental conditions and crystal packing (*52, 53*). The estimated precision of other PPI databases is low (4% – 12%), but the confident subset selected by each database has higher precision (20% – 30%, **Fig. 3D**). Compared to PPIs in databases, predicted PPIs display a much stronger tendency to participate in the same biological processes and locate to the same subcellular components (**Fig. 3E**). Thus, our predictions can be used to infer the functions and subcellular localities of poorly characterized proteins by finding their well-characterized partners.

We computed 3D structures of predicted PPIs using AF2 and AFmm and selected the best models (see **supplemental Methods M6.2**). We compared the properties of predicted PPIs against experimentally determined PPIs, i.e. 9,725 human PPIs with orthologous templates in the PDB. Similar to PDB complexes, most (76%) predicted PPIs are primarily mediated by globular domains. However, interfaces mediated by flexible helices or IDRs are much rarer among predicted PPIs than in PDB complexes (**Fig. 3F**), consistent with the fact that PPIs with smaller interfaces are harder to predict (**supplemental fig. S28**). A higher fraction of predicted PPI interfaces involves transmembrane helices (TMHs) than experimentally determined. We wondered if the hydrophobic nature of TMHs make them prone to false positives because DL networks might have learned to pack hydrophobic TMHs against each other even if they do not coevolve or interact *in vivo*. We tested the performance of our pipeline on pairs of transmembrane proteins (TMPs) and found that it predicts a larger fraction (magenta curve versus blue curve in **Fig. 3G**) of their interactions at a high precision, including those mediated by single-TMH proteins. These observations suggest that our pipeline is suitable for identifying interactions between TMPs, which are hard to detect experimentally (*54*–*56*). The 3D models of predicted PPIs in this study nearly triple the number of human PPIs with high-quality 3D structures. Thousands of post-translational modification (PTM) sites, functional (active and substrate/cofactor binding) sites, and disease-associated single amino acid variations (SAVs) (*57*–*59*) map to the predicted PPI interfaces (**Fig. 3H**), and thus in-depth analysis of our models will provide numerous mechanistic insights of protein functions.

### Biological insights from novel PPIs

Among the 18,307 predictions, 2,786 (15%) PPIs or their orthologs have experimental 3D structures; another 9,953 (54.3%) are reported in PPI databases (UniProt, BioGRID, and STRING physical); the remaining 5,568 pairs (30.4%) are novel compared to these established sources of PPIs (**Fig. 4A**). Many of the novel predictions are indirectly supported by other evidence such as genetic interactions in STRING (3,443 pairs) and homologous (not orthologous) templates in the PDB (2,673 pairs); some have been experimentally tested based on literature search but are not yet propagated into PPI databases. Based on a recent catalog of poorly characterized proteins (*60*), 1,582 predicted PPIs involve proteins of unknown functions: linking them to well-characterized proteins provides shortcuts to investigating their functions. For our in-depth analysis, we prioritized 2,895 PPIs that are absent from PPI databases and lack homologous PDB templates. We classified predicted PPIs into functional categories according to UniProt keywords, and found categories enriched (P-value < 0.01) in such novel PPIs (**Fig. 4B**). Focusing on several such categories, below we illustrate how our results can be used to study protein function. Additionally, many novel predictions involve proteins associated with genetic disorders or cancers (**Fig. 4C-4H**) and potentially offer insights into human disease (see **supplemental Results R2.1 and R2.6)**.

**Figure 4.**
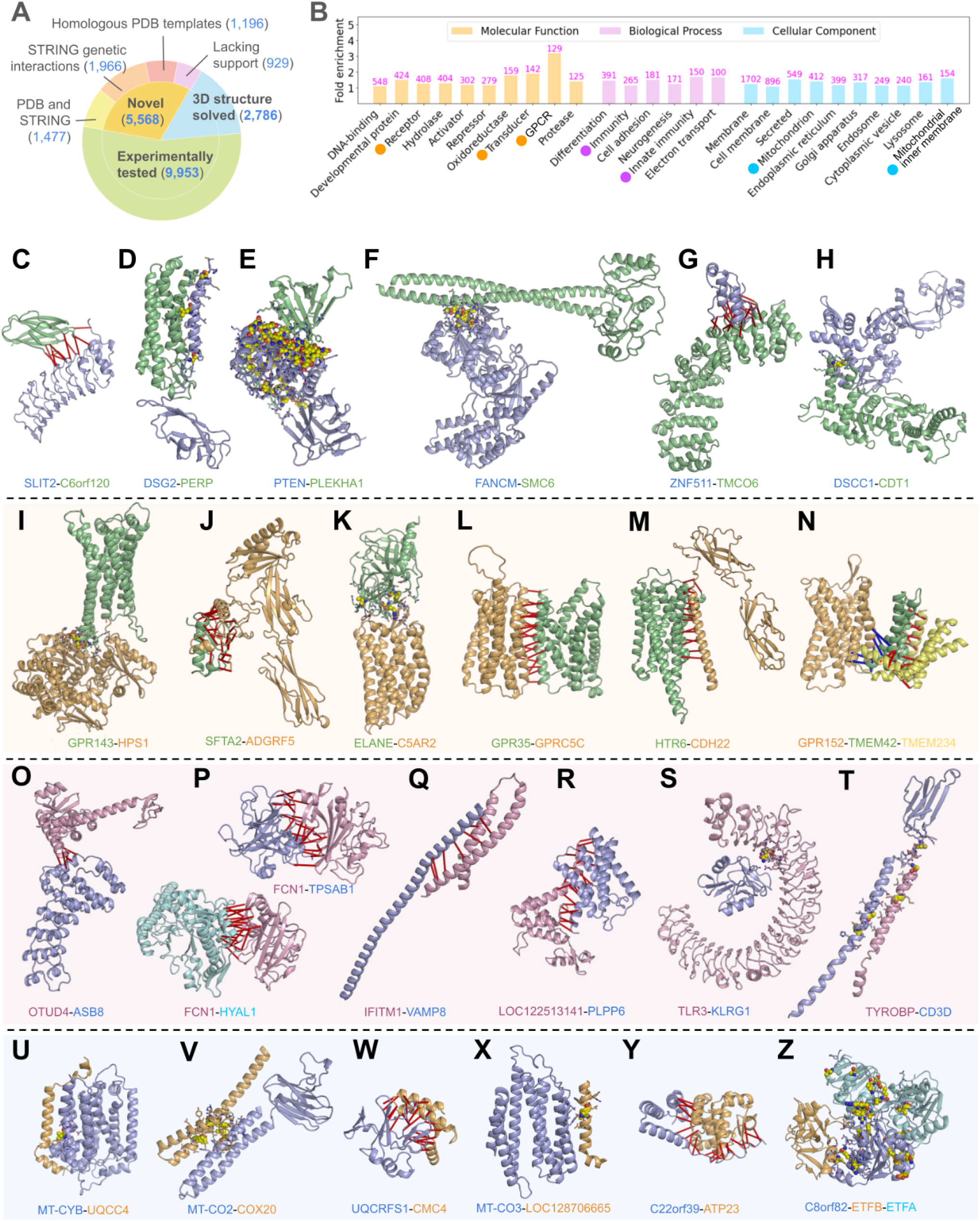
Examples of novel functional insights. **(A)** A pie chart showing the fraction of predicted PPIs with experimental evidence. 3D structure solved: with orthologous PDB template; experimentally tested: present in UniProt, STRING (physical), or BioGRID databases, but 3D structures not resolved; the remaining novel PPIs are further partitioned by whether they show evidence of genetic interactions, have homologous templates, or both. **(B)** Functional categories (UniProt keywords) that are enriched (P-value < 0.01) with novel PPIs (numbers labeled above the bars). Dots by the labels mark categories discussed in the main text. **(C-H)** PPIs involving cancer-related proteins. **(I-N)** PPIs involving GPCRs. **(O-T)** PPIs related to immunity. **(U-Z)** Mitochondrial PPIs. **(C-Z)** Yellow spheres indicate interface residues associated with disease-causing SAVs; otherwise, red and blue bars connect residues with their most confidently predicted inter-protein contacts.

### G protein-coupled receptors (GPCRs)

GPCRs are a large family of TMPs that detect extracellular signals and mediate cellular responses (*61*). They play vital roles in human physiology and are important drug targets (*62*). We detect novel interactions between GPCRs and their putative downstream signaling molecules and ligands or modulators. For example, we predict an interaction between GPR143 and HPS1 (**Fig. 4I**). GPR143 is an atypical GPCR localized in endolysosomes and melanosomes, while HPS1 is a component of a guanine nucleotide exchange factor (GEF) complex that activates Ras-related GTPase RAB32 and RAB38 (*63*). The GPR143-HPS1 interaction is intriguing because both proteins are expressed in melanosomes, involved in melanosome biogenesis, and associated with albinism (*64*–*67*). In addition, we detect a potential ligand or modulator for ADGRF5 (**Fig. 4J**), an adhesion GPCR critical for lung surfactant homeostasis (*68*); its predicted interaction with SFTA2 via its extracellular domain suggests that this secreted small protein, predominantly expressed in lung (*69*), can be the ligand or modulator for ADGRF5. Further, we predict an interaction between C5AR2 (a GPCR) and ELANE, a serine protease (**Fig. 4K**). The former is known to be a receptor for C5a anaphylatoxin peptide and plays a role in chemotaxis and inflammation (*70*), while the latter inhibits C5a-dependent chemotaxis and neutrophil function (*71*). This predicted C5AR2-ELANE interaction and their opposite roles in chemotaxis suggests that ELANE is a negative modulator of C5AR2.

We also observe examples of predicted interactions between pairs of GPCRs as well as between GPCRs and other TMPs that may function together to deliver complex cellular responses. For example, we predict an interaction between GPR35 and GPRC5C (**Fig. 4L**). GPR35 is involved in chemotaxis of macrophage and inflammation responses (*72*–*74*), while GPRC5C promotes the dormancy of hematopoietic stem cells (*75*). This predicted heterodimer, with a non-conventional dimer interface (between TMH1&2 of GPR35 and TMH6&7 of GPRC5C), may unravel novel signaling pathways in blood cell differentiation. In addition, HTR6, a GPCR mediating neurotransmission (*76*), is predicted to interact with CDH22, a single-TMH protein in the cadherin family involved in cell adhesion in the brain (*77, 78*) (**Fig. 4M**). Such an interaction provides a potential link between GPCR signaling and cell adhesion. Finally, we predict an interaction between an orphan GPCR, GPR152 (*79*), and TMEM42 (**Fig. 4N**), an uncharacterized TMP showing homology to a variety of transporters according to HHpred (*80*). The GPR152-TMEM42 interaction points to the possibility of transporter activity regulation by a GPCR.

### Immunity proteins

The evolutionary arms race between hosts and pathogens drive hosts to utilize a wide range of mechanisms to defend against infectious bacteria and viruses (*81*). We predict 265 novel PPIs involving immunity proteins, many of which mediate innate immunity signaling pathways. For instance, we predict an intriguing interaction between a deubiquitinase, OTUD4, and an E3 ubiquitin ligase, ASB8 (**Fig. 4O**). OTUD4 removes ubiquitin chains from MAVS, an important innate immune adapter to detect viral RNA, preventing its degradation (*82*); in contrast, ASB8 introduces polyubiquitins to MAVS’s downstream protein kinase TBK1, promoting its degradation (*83*). The interaction of these two proteins with opposing roles might help maintain a balance in the regulation of protein turnover during innate immune responses. We predict interactions between FCN1, an extracellular receptor for pathogen recognition (*84*), and several extracellular enzymes (**Fig. 4P**), including two tryptases (TPSAB1 and TPSAB2) and a hyaluronidase (HYAL1). These interactions suggest a potential role of FCN1 in innate immune responses in the extracellular matrix, opening up new directions to investigate its functions. Interestingly, we predict an interaction between an interferon-induced TMP (IFITM1) and a SNARE protein VAMP8 (**Fig. 4Q**). IFITM1 inhibits the entry of many viruses, including SARS-CoV, into the host cells (*85*), while VAMP8 is involved in the fusion of autophagosomes and lysosomes (*86, 87*). This suggests a potential mechanism for IFITM1’s antiviral activity by recruiting VAMP8 and promoting the delivery of virus cargo in the autophagosomes to lysosomes for degradation.

Other novel PPI predictions involve proteins with roles in the differentiation and activation of specialized immune cells. For instance, PLPP6 is an enzyme contributing to neutrophil activation via dephosphorylation of presqualene diphosphate, a potent inhibitor of this process (*88, 89*). We predict an interaction between PLPP6 and an unknown protein LOC122513141 (**Fig. 4R**). Both proteins are located in the Endoplasmic Reticulum (ER) membrane, and this predicted interaction suggests that LOC122513141 could serve as a regulator of PLPP6 function. In addition, KLRG1 is a TM receptor that inhibits the activity of natural killer cells and effector T cells (*90, 91*). The extracellular domain of KLRG1 is predicted to interact with TLR3, a key player in recognizing double-stranded RNA and inducing immune responses (*92*) (**Fig. 4S**). Such an interaction suggests a potential mechanism for KLRG1’s function as an immunity inhibitor. Finally, we predict an interaction between two remote homologs, TYROBP and CD3D (**Fig. 4T**). The former mediates the activation of a variety of immune cells (*93*–*95*), while the latter transmits signals from T-cell receptors and activates T-cells (*96*). The TYROBP-CD3D interaction suggests TYROBP’ role in T-cell activation, potentially an example of the complicated cooperation between cell surface receptors in activating various immune cells.

### Mitochondrial targeting proteins

More than half of PPIs not found in databases (1,697 out of the 2,895) involve TMPs. These TMPs are frequently located in cellular organelles, such as mitochondrion, ER, Golgi apparatus, endosome, and lysosome (**Fig. 4B**). Many PPIs we predict in the mitochondria are related to the assembly of mitochondrial complexes that constitute the respiratory electron transport chain (ETC). For example, we detect interactions between the complex III component MT-CYB (*97*) and the complex III assembly factor UQCC4 (*98*) (**Fig. 4U**), and between the complex IV subunit MT-CO2 (*99*) and the complex IV assembly factor COX20 (*100*) (**Fig. 4V**). Our predictions also reveal several proteins of unknown functions that may serve as additional assembly factors of the ETC complexes. For example, we link CMC4 (**Fig. 4W**), LOC128706665 (**Fig. 4X**), C22orf39 (**Fig. 4Y**), and C8orf82 (**Fig. 4Z**) to components or assembly factors of complex III, IV, V, and the electron transfer flavoprotein complex, respectively. Among them, LOC128706665 was annotated as “alternative protein MKKS” in UniProt because its coding region overlaps with the 5’ UTR of the molecular chaperone MKKS that primarily resides in the centrosome (*101*). However, our prediction suggests that LOC128706665 should be a novel mitochondrial protein not related to MKKS.

### New multi-protein complexes

By integrating our predictions and PPIs derived from orthologous PDB templates, we identify 379 predicted multi-protein complexes, wherein each component interacts with at least two other components (**supplemental Methods M6.3**). We classify PPIs in these complexes into three categories: (1) with experimental structures, (2) confident entries of PPI databases; (3) no or weak experimental support. We focus on putative complexes dominated by PPIs lacking confident experimental support (3rd category). These complexes (**Fig. 5**) are frequently scaffolded by proteins with long IDRs or flexible helices that interact with multiple components. They likely adopt flexible 3D structures and are hard for experimental characterization; predicting their existence and modeling their 3D structures open new directions for future studies.

**Figure 5.**
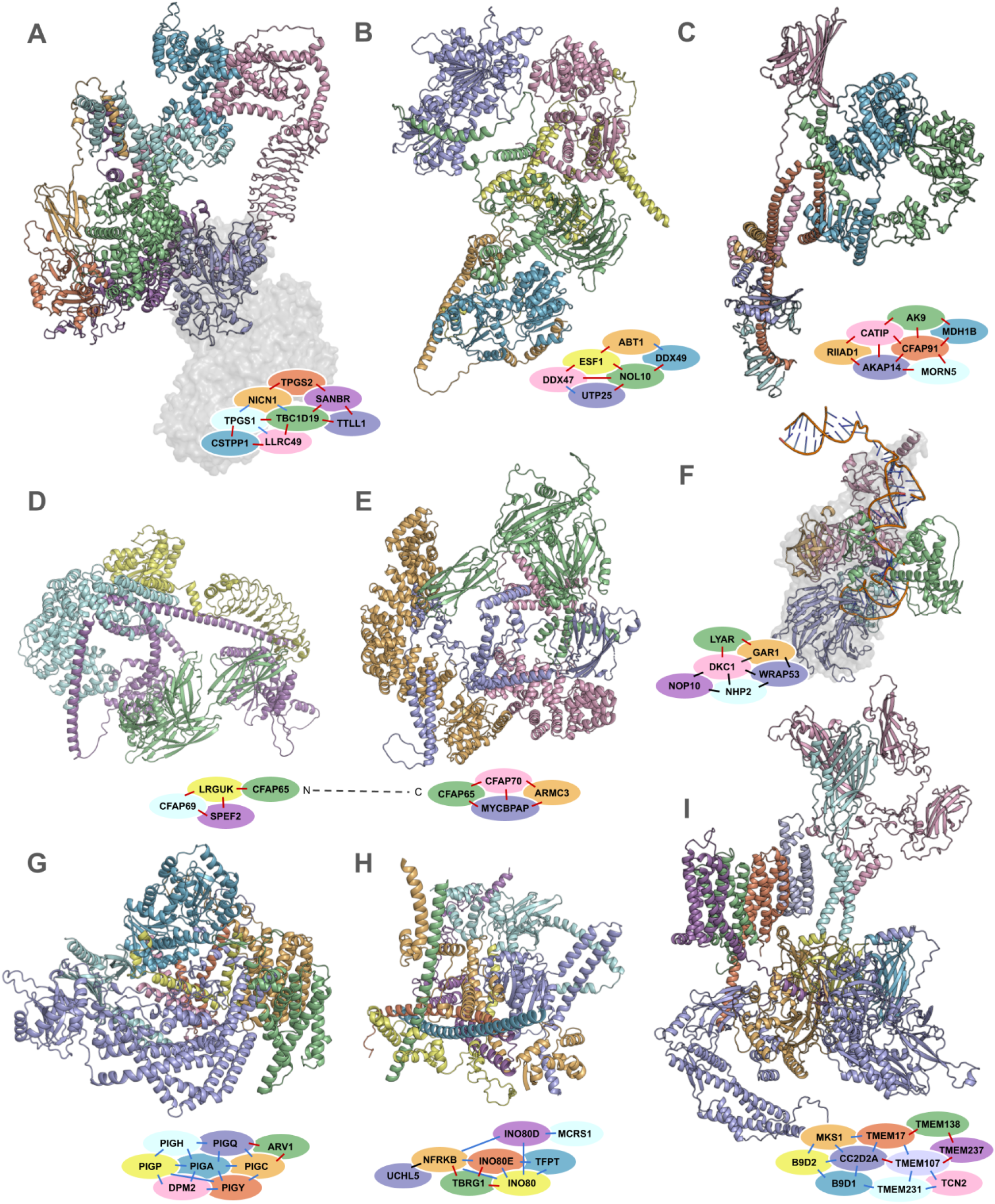
Examples of novel protein complexes (A-E) and predicted new components to known complexes (F-I). Each panel contains the 3D structure of a complex and a network graph. Nodes in the graph are labeled by gene names and colored the same as the corresponding subunits in the 3D structure. Edges in the graph represent PPIs and are colored as follows: black, PPIs with experimental structures; blue, confident PPIs from databases; red: novel PPIs or with weak experimental support. **(A)** The tubulin polyglutamylase complex. An experimental structure (PDB: 5jh7) of the catalytic subunit and its substrate (tubulins) is shown as surface representation and colored in gray. **(B)** A complex connecting basal transcription regulation with ribosome biosynthesis. **(C-E)** Complexes related to cilia and flagella biogenesis. **(F)** Adding LYAR to the telomere maintenance complex; an experimental structure of this complex (PDB: 8ouf) is shown as surface representation and colored in gray. **(G)** Adding ARV1 to a GPI anchor biosynthesis complex. **(H)** Adding TBRG1 to the INO80 chromatin remodeling complex. **(I)** Adding TMEM138 to the transition zone complex.

Tubulin polyglutamylase (TPG) is an enzyme complex responsible for adding polyglutamate chains to the glutamate residues in the C-terminal tail of tubulin, a modification important for the regulation of microtubule functions (*102, 103*). In addition to the catalytic subunit TTLL1, the TPG complex is known to contain four other subunits: TPGS1, TPGS2, LRRC49 and NICN1 (*104*). We predict 3 additional subunits of TPG: TBC1D19, CSTPP1, and SANBR, each interacting with multiple known TPG components (**Fig. 5A**). TBC1D19 is the central component of the complex, interacting with 5 other subunits (TTLL1, LRRC49, NICN1, TPGS1, and SANBR). TBC1D19 and SANBR directly interact with the catalytic subunit TTLL1 via interfaces apart from its ATP binding and catalytic sites (*105*), possibly affecting the activity of TTLL1 by allosteric regulation.

Another predicted complex consisting of NOL10, UTP25, DDX47, ESF1, ABT1, and DDX49, could link the regulation of basal transcription with ribosome biogenesis (**Fig. 5B**). The two processes should be coordinated to ensure that transcribed mRNA can be efficiently translated into proteins. The central component, NOL10, is a poorly characterized nucleolar protein made of an N-terminal beta-propeller domain and C-terminal flexible helices. ABT1 and its interacting partner ESF1 are suggested to regulate basal transcription (*106, 107*), while UTP25 and DDX47 are implicated in ribosome biogenesis (*108*). The connection between this complex and the ribosome is also supported by its predicted interactions with ribosomal proteins via ESF1, DDX47, and NOL10; the N-domain of NOL10 is even part of an experimental ribosome structure (PDB: 7MQ8) (*109*). Future studies of this putative complex may reveal new insights about the coordination between the two fundamental processes, transcription and translation.

We predict several complexes made of proteins associated with the biogenesis of cilia and flagella, two cellular organelles sharing the same molecular machineries during their formation (*110*). The flagellum enables a sperm to swim towards and fertilize the egg (*111*), linking these complexes to sperm motility and reproduction. The first predicted complex includes CATIP, RIIAD1, MDH1B, CFAP91, AKAP14, AK9, and MORN5, among which CFAP91, an intrinsically disordered protein, serves as a scaffold to assemble all but one (RIIAD1) other proteins (**Fig. 5C**). Involvement of this complex in flagella biogenesis and sperm motility is supported by several facts: 1) CFAP91 and CATIP have been shown to participate in cell skeleton organization during cilia/flagella biogenesis (*112*–*115*); 2) AK9 affects sperm mobility (*116*); and 3) defects in CATIP and AK9 are associated with asthenozoospermia (*115, 116*). We predict two additional complexes which involve proteins implicated in sperm biogenesis. One consists of CFAP69, LRGUK, and SPEF2 (**Fig. 5D**); the other is made of CFAP65, CFAP70, MYCBPAP, and ARMC3 (**Fig. 5E**). These complexes are likely parts of a larger machinery, connected via an interaction between CFAP65 and LRGUK.

### New components of known protein complexes

We identify 83 potential new components of known human protein complexes (**supplemental Methods M6.3**) in the complex portal database (*117*). Each new component is predicted to interact with at least two subunits of a known complex and primarily interact (>50% of predicted PPIs) with this complex. For example, LYAR, a nuclear protein involved in rRNA processing (*118*), is predicted to interact with two telomere maintenance complex (TMC) proteins, DKC1 and GAR1 (**Fig. 5F**). Additional components of TMC include NOP10, NHP2, and WRAP53, and they participate in the processing and trafficking of the RNA template (*119*–*123*) used to synthesize telomeric DNA (*124*). Multiple components of the TMC, including DKC1, GAR1, NOP10, and NHP2, have been shown to participate in rRNA processing and ribosome biosynthesis (*119, 125*–*127*). The binding of LYAR to these components might channel the TMC to process rRNA instead of telomeric RNA template.

We predict a new component to the Glycosylphosphatidylinositol-N-acetylglucosaminyltransferase (GPI-GnT) complex, which catalyzes the first step of GPI anchor biosynthesis (*128*). The GPI-GnT complex consists of seven subunits: PIGA, PIGC, PIGH, PIGP, PIGQ, PIGY, and DPM2 (*128*), and its 3D structure remains unresolved. ARV1, a TMP found to be related to GPI biosynthesis (*129*–*131*), is predicted to interact with PIGC and PIGQ. Our 3D model of the GPI-GnT complex with ARV1 (**Fig. 5G**) shows that a hydrophobic helix (residues 422-442) in the catalytic subunit, PIGA, is positioned in parallel to the membrane and docked onto a heterotrimer of the 2-pass TMPs: PIGP, PIGY, and DPM2. ARV1 (5 TMHs) is positioned in between PIGC (8 TMHs) and PIGQ, and several hydrophobic helices of PIGQ lie in parallel to the membrane. These subunits form a ring-like structure with a large cavity in the middle of the membrane that could facilitate substrate binding or transport of the product.

We predict multiple interactions between components of the INO80 chromatin remodeling complex (CRC) (*132*), which allowed us to model a subcomplex comprised of the regulatory subunits (INO80D, INO80E, NFRKB, TFPT, MCRS1, and UCHL5). This model reveals that the regulatory subcomplex interacts with the N-terminal IDR of the CRC core subunits, INO80, while the C-terminal domains of INO80 interact with other core CRC subunits to perform the chromatin remodeling function (*133*). The regulatory subunits likely help to coordinate the complicated roles of CRC, including transcription regulation, DNA replication, and DNA repair (*134, 135*). Recently, it was suggested that TBRG1 could be a new subunit of the CRC based on its interaction with INO80 (*136*). Consistent with this study, we predict that TBRG1 interacts with multiple CRC subunits through a long N-terminal helix, not only INO80, but also TFPT, NFRKB, and INO80E **(Fig. 5H)**. TBRG1, as well as TFPT, are related to the regulation of cell cycle and cell growth (*137*), potentially linking the CRC function to these processes.

Finally, we predict a new component of the MKS transition zone complex (TZC) located between the basal body and axoneme of cilia and flagella (*138, 139*). Predicted PPIs between TZC components enabled us to model a large complex **(Fig. 5I)** spanning the extracellular (TCN2 and TMEM231), membrane (TMEM17, TMEM107, TMEM138, and TMEM237), and cytoplasmic space (MKS1, B9D1, B9D2, and CC2D2A). TMEM138 has been suggested to localize to the transition zone based on immunofluorescence assays and is associated with the same ciliopathies as other transition zone proteins (*140*). We further predict TMEM138 as a subunit of TZC, sandwiched between TMEM17 and TMEM237 in our computed model. We included one copy of each TZC subunit in our model; however, it is likely that these proteins will homo-oligomerize to form a much larger complex which constitutes the transition zone.

## Conclusions

Despite significant advances in DL methods for modeling the 3D structures of protein complexes, distinguishing the 74k-200k true PPIs amongst the 200M pairs of human proteins has remained a grand challenge. Here, we tackle this challenge by leveraging the largest sequence and structure datasets available and focusing on two key innovations: 1) strengthening the coevolutionary signals between interacting proteins using 7-fold deeper MSAs directly assembled from genomic data and 2) developing a new and fast DL network, trained on 10 times larger datasets derived from predicted DDIs, to differentiate true PPIs from random protein pairs. These improvements enabled us to systematically identify human PPIs on a proteome-wide scale, predicting over 18k interactions with an expected precision of 90%. Notably, these predictions include more than 5.5k novel PPIs, particularly among TMPs that are challenging to characterize experimentally. These predictions will offer numerous valuable insights into human biology and diseases, as demonstrated by our biological vignettes. The power of our approach will continue to grow as more sequence and structure data become available and DL techniques advance. Integrating the rapidly improving computational approaches with experimental studies, we show that the nearly complete characterization of the human interactome is within reach.

## Availability

We present our predictions and intermediate results at http://prodata.swmed.edu/humanPPI, which allows users to easily navigate through our findings and accelerate future discoveries. We share our omicMSAs, training datasets, trained weights of RF2-ppi, predicted interaction probabilities, and 3D models of protein complexes at: https://conglab.swmed.edu/humanPPI/humanPPI_download.html. We deposited the RF2-ppi network at GitHub: https://github.com/CongLabCode/RoseTTAFold2-PPI. To seamlessly use our omicMSA to build protein complexes with AF2 or AFmm, please use our Colab notebook at: https://colab.research.google.com/drive/1suhoIB5q6xn0APFHJE8c1eMiCuv9gCk_

## Acknowledgment

The authors thank Nick V. Grishin, Jian Zhou, Rohith Krishna, Frank Dimaio, Edin Muratspahic, and Helen H. Hobbs for inspiring discussions and insightful suggestions. We also thank Luki Goldschmidt and Aaron Guillory for computing resource management, and Linda Stewart and Lance Stuart for logistical support. QC is a Southwestern Medical Foundation Endowed Scholar and DB is a Howard Hughes Medical Institute Investigator. This research is supported by I-2095-20220331 and V-I-0004-20230731 from the Welch Foundation, 1K99AI180984-01A1, 5-R01-HL-159946-03, and contract No. 75N93022C00036 from NIH, Department of Health and Human Services, Bill & Melinda Gates Foundation Investments INV-010680 and INV-043758, Novo Nordisk A/S contract RDC 2022-002902, NRF/MSIT (RS-2023-00210147, RS-2024-00440824, and RS-2024-00407331), IITP/MSIT (RS-2023-00220628), and Perlmutter grant NERSC award BER-ERCAP0022018 for access to the Perlmutter high-performance computing resources.

