## Supplemental Information for "Computing the Human Interactome"


**Supplemental Methods** **3**
M1. Gathering experimental data and preparing benchmark sets 3
M2. Preparing training datasets for PPI prediction 5
M2.1. Pairs of interacting proteins derived from protein complexes in PDB 6
M2.2. DPAM-IPR: a Domain Parser for AF2 Models utilizing InterPRo data 8
M2.3. Pairs of interacting domains derived from multi-domain AFDB models 13
M2.4. Preparing input data for the training datasets 15
M3. Developing Artificial Intelligence networks for PPI prediction 16
M3.1. Simplifying the RoseTTAFold2 network and performance evaluation 16
M3.2. Testing training dataset and model architectures 17
M3.3. Comparing the performance of RF2-ppi against other tools 21
M4. Generating multiple sequence alignments from genomic data 23
M4.1. Stage1: gathering genomic datasets for Eukaryotes 23
M4.2 Stage2: assembling orthologs from draft genomes associated with proteomes 26
M4.3 Stage3: assembling orthologs from draft genomes 32
M4.4 Stage4: assembling orthologs from genomic and transcriptomic reads 33
M4.5 Stage5: generating multiple sequence alignments for human proteins 37
M4.6 Evaluating the performance of our multiple sequence alignments 40
M5. Developing a hierarchical PPI prediction pipeline 42
M5.1 Preparing proteins for PPI screening 42
M5.2 Prioritizing pairs based on subcellular localization information 42
M5.3 Prioritizing pairs based on coevolution signals 43
M5.4 Using RF2-ppi to identify PPIs 46
M5.5 Using AF2, AFmm, and ColabFold for PPI modeling 48
M5.6 Selecting the final set of predicted PPIs 49
M6. Analyzing predicted PPIs 51
M6.1 Integrating predicted PPIs with knowledge in databases 51
M6.2 Generating and analyzing 3D models for predicted PPIs 52
M6.3 Identifying high-order protein complexes and new components to known complexes based on our predicted PPIs 55
**Supplemental Results** **57**
R1. Features of PPIs that cannot be predicted by current methods 57
R2. Biological insights revealed from predicted PPIs 60
R2.1 PPIs of proteins involved in cancer 60
R2.2 PPIs of G protein-coupled receptors and other membrane proteins 63
R2.3 PPIs of proteins involved in immunity 65
R2.4 PPIs of mitochondrial proteins 68
R2.5 PPIs of proteins involved in cilium function 70
R2.6 PPIs with disease mutations mapped to interaction interfaces 71
R2.7 Other interesting PPIs 74
R3. Multi-subunit complex modeling 76
**Supplemental References** **84**

### Supplemental Methods

#### M1. Gathering experimental data and preparing benchmark sets

We downloaded 3D structure models of the human proteome that were modeled by AlphaFold (*1*) and released through the AlphaFold protein structure DataBase (AFDB) (*2*). These models cover nearly all reviewed entries from the human proteome UP000005640 released in Uniprot (*3*) version 2021_02. We used AFDB models to parse each human protein into domains, which allowed us to split large proteins into smaller segments that fit into the limited memory of our GPUs (mostly below 48 GB). We excluded a small fraction (208, 1%) of long proteins that were modeled in multiple frames, and the remaining 20,296 human proteins which we call, the **human protein set**, were used in this study.

We utilized three well-established and constantly updated databases to gather experimental data about **physical** protein-protein interactions (PPIs) formed by proteins from **the human protein set** (**Figure S1**). First, we obtained the interacting partners for each protein from UniProt (accessed in Nov 2023) (*3*, *4*) by turning on the "interacts with" column and downloading the customized table. We obtained 75,739 candidate interactions, namely, the **UniProt PPIs**. Second, we downloaded the entire set of interactions (BIOGRID-ALL), the human-specific set (BIOGRID-ORGANISM-Homo_sapiens), and the set of confident PPIs validated in multiple experiments (BIOGRID-MV-Physical) from the BioGRID database (version 4.4.225) (*5*). Human PPIs from BioGRID are mostly physical interactions, with only 1.5% genetic interactions; we thus did not discard the genetic interactions from these datasets. From the BioGRID sets, we extracted PPIs between proteins in our **human protein set**, resulting in 771,432 candidate interacting pairs, i.e., **BioGRID PPIs**. Among these pairs, 77,147 are regarded as confident PPIs according to BioGRID (belonging to the multi-validated set), and we name them as **confident BioGRID PPIs**. Third, we downloaded physical interactions of human proteins (9606.protein.physical.links) from the STRING (version 12.0) database (*6*). In addition, we downloaded the human protein sequences (9606.protein.sequences) used in the STRING database, which allowed us to map the STRING human proteins to our **human protein set** by MMseqs (-s 7 -a 1) (*7*). We considered a STRING protein to be equivalent to one in our **human protein set** if their identity in sequence is above 95% and the aligned residues covered more than 90% of the query and the hit. Based on information from STRING, we identified 626,513 candidate PPIs, i.e., **STRING PPIs**; among them, 68,761 PPIs are considered confident (**confident STRING PPIs**) with STRING combined scores above 700 (a cutoff suggested by the STRING database).

We compared the sets of **UniProt PPIs**, **confident BioGRID PPIs**, and **confident STRING PPIs**, and found a surprisingly poor consistency between the three sets (**Figure S1**): only 3,988 PPIs were included in all three datasets, with 22,184 additional pairs supported by two out of the three databases. This poor consistency highlights the difficulty of identifying true interacting partners for a human protein. PPIs from these databases are mostly uncovered from large-scale experimental studies that are plagued by non-specific interactions in non-physiological conditions. Therefore, they are expected to contain a very high rate of false positives. We took the 3,988 PPIs shared by all three databases as the **positive control set**. This positive control set was to monitor the performance of our newly developed AI network, RoseTTAFold2-ppi (**RF2-ppi**) in the process of optimizing the training routine and network architecture. To ensure that the improved performance we observed is not a result of over-optimization on this positive control set, we derived an additional **positive validation set** made up of 22,184 PPIs supported by only two databases.

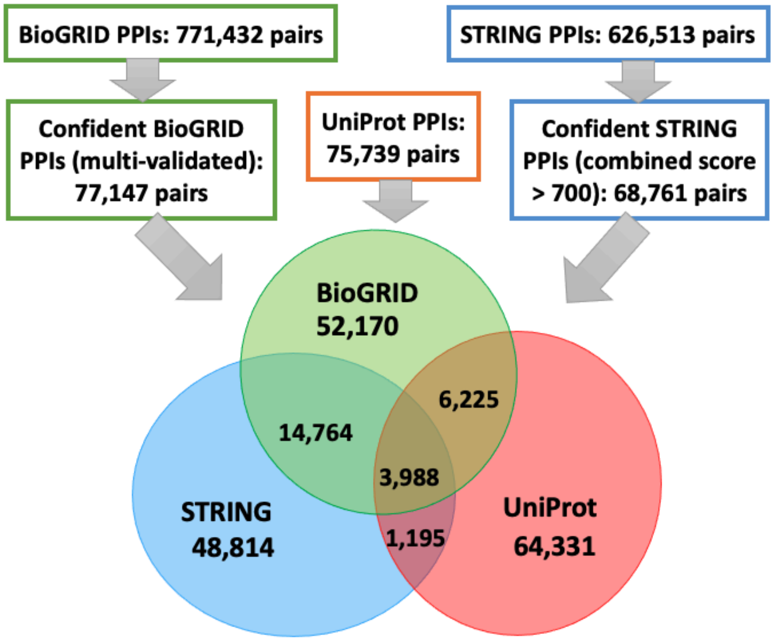


**Figure S1. The procedure to prepare positive controls (top) and a Venn diagram showing the consistency between confident PPIs obtained from different databases.**

In addition to the sets of physically interacting protein pairs mentioned above, we obtained functionally related human protein pairs from the STRING database (9606.protein.links). While functionally related proteins mostly do not directly interact with each other, they are useful to derive the negative controls for which we want to exclude any protein pairs that may function together. We enumerated all possible pairs of proteins in our **human protein set**, and excluded pairs from any of the aforementioned sets of physical or genetic interactions. In addition, we removed any pairs that are homologous to pairs in these datasets because an interacting partner of a protein is more likely to interact with its homologs than another random protein. As a result, we obtained a **negative control set** containing 140,156,818 random protein pairs without prior evidence of interacting. The entire lists of the **human protein set**, **positive control set**, **positive validation set**, and **negative control set** can be downloaded online from https://conglab.swmed.edu/humanPPI/humanPPI_download.html and https://prodata.swmed.edu/humanPPI/ bulk_download.

#### M2. Preparing training datasets for PPI prediction

The interacting chains from the PDB (**PPI set**) were used to train RoseTTAFold2 (RF2) (*8*) and AlphaFold-multimer (AFmm) (*9*) to improve their ability in modeling 3D structures of protein complexes. However, one drawback of the **PPI set** is its small size; after clustering at 30% sequence identity (MMseqs: --min-seq-id 0.3, -c 0.8), the **PPI set** used to train RF2 contains less than 20,000 clusters. It was shown that using larger distilled datasets, e.g., AF2 models for a diverse set of proteins, has significantly improved the performance of AF2 (*1*) and RF2. We therefore wondered whether we can improve the performance of RF2-ppi in PPI prediction by enlarging the training dataset with predicted 3D structures of protein complexes. However, confidently predicted protein complexes by these state-of-the-art methods are rare. Instead, AF2 models in AFDB contain a large number of interacting domains. These domains are the evolutionary and structure units (*10*, *11*), and they are recombined in evolution to constitute proteins to perform specific functions (*12*, *13*). We reasoned that the interactions between these domains should resemble the interaction between proteins (*14*, *15*), and thus, a dataset of domain-domain interactions (DDI) derived from AFDB models (**DDI set**) might be useful to develop AI networks for modeling protein complexes and predicting true PPIs among random protein pairs.

##### M2.1. Pairs of interacting proteins derived from protein complexes in PDB

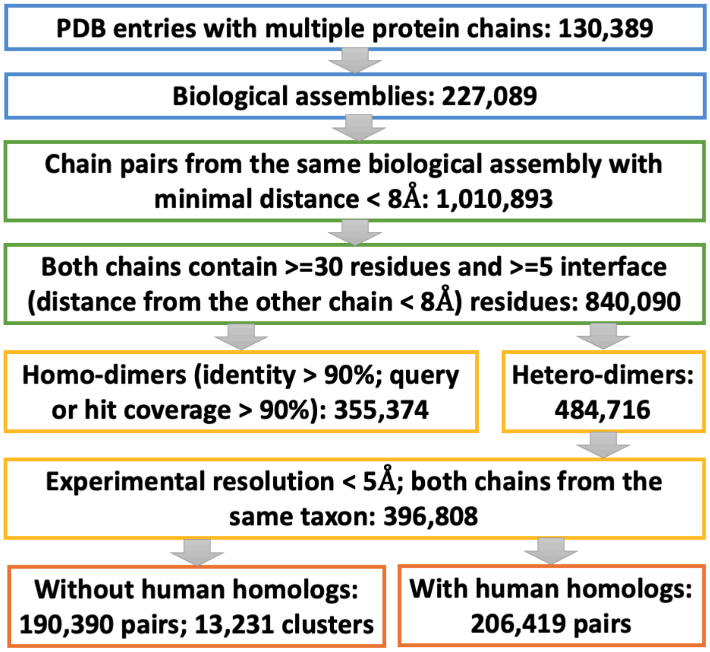


**Figure S2. The procedure to prepare positive PPI training data from protein complexes in PDB.**

The **PPI set** used to train the latest RF2 was derived from PDB entries released prior to Aug 2021. We updated this dataset to include PDB entries of complexes released before January, 2024 using a pipeline shown in **Figure S2**. We downloaded PDB entries in the mmcif format and focused on 130,389 entries with more than one polypeptide chain. We extracted biological assemblies from each PDB entry. Within each biological assembly, we detected interacting chain pairs satisfying the following criteria: (1) both chains have more than 30 residues, (2) more than 5 residues from both chains are interacting with (minimal distance < 8Å) residues from the other chains. As a result, we identified a total of 840,090 interacting chain pairs.

We compared the sequences of these interacting chain pairs and considered pairs with high sequence similarity (identity > 90%; query or hit coverage > 90% by MMseqs) as homo-dimers. We further filtered the remaining 484,716 hetero-dimers using two criteria: (1) both chains should be from the same organism (with the same TaxID); (2) they belong to PDB entries determined by X-ray crystallography or Cryo-EM with resolution below 5Å. As a result, we obtained 396,808 heterodimeric protein complexes, which we call the **positive PPI set**. We clustered single protein chains in this set at 30% sequence identity cutoff using MMseqs (--min-seq-id 0.3 -c 0.8 --cov-mode 0); based on these clusters of single chains, we further clustered the binary protein complexes by placing two complexes, AB and A'B', in the same cluster if A' is in the same single-chain cluster as A and B' is in the same single-chain cluster as B. The updated **positive PPI set** contains 24,583 clusters (compared to 18,568 clusters in the PPI set used to train RF2). During training, each cluster, instead of each case, is weighted equally to avoid overemphasizing large protein families. Finally, because we focused on predicting the human interactome in this work, to avoid any data leakage from experimentally determined human protein complexes in PDB, we excluded any pairs where both PDB chains showed homology to human proteins detectable by MMseqs (e-value < 0.00001), resulting in a total of 190,390 PPIs from 13,231 clusters. This dataset, named as the **filtered positive PPI set,** is the primary dataset used to train AI networks in this study.

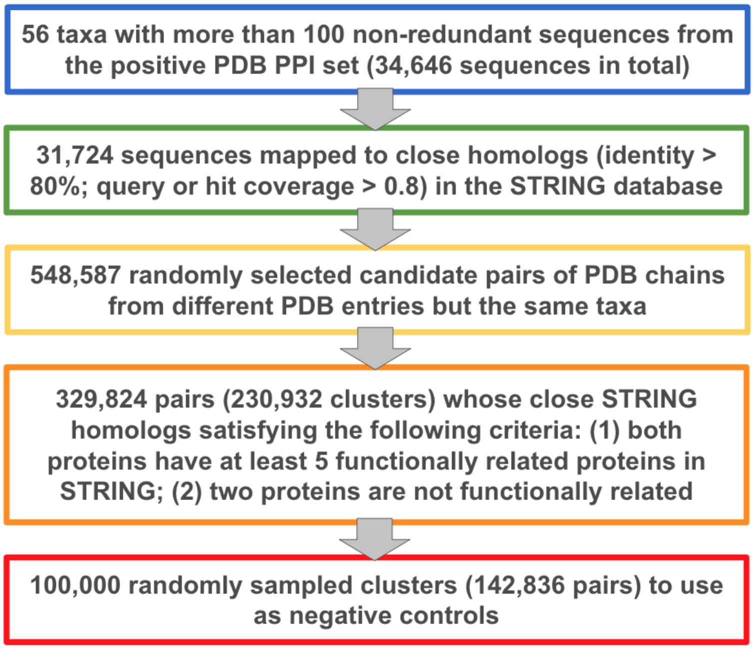


**Figure S3. The procedure to prepare negative PPI training dataset by randomly pairing PDB chains that are NOT functionally related according to STRING.**

In order to train an AI network to distinguish true interacting proteins from random pairs of proteins, we also assembled a negative PPI dataset (**Figure S3**) by randomly pairing PDB chains from the **positive PPI set**. These random pairs need to be from the same organism (according to TaxID) but show no evidence of interaction. We analyzed the taxonomy information for PDB chains in the **positive PPI set** and focused on 34,646 non-redundant PDB sequences from 56 taxa, where each taxon has more than 100 sequences. We mapped these sequences to entries in the STRING database using MMseqs, keeping only 31,724 sequences that can be mapped to closely related STRING sequences (query or hit coverage > 0.8, identity > 80%). The STRING database focuses on genetic interactions, and it provides information about functionally related protein pairs. Thus, mapping to STRING entries allowed us to evaluate whether a pair of PDB chains represent proteins that are functionally related.

Next, we randomly selected candidate negative control pairs from each taxon, requiring the number of such pairs in a taxon to be no more than twice of the number of pairs from this taxon in the **positive PPI set**. As a result, 548,587 pairs were selected as candidates for negative controls, each from the same source organism but different PDB entries. We considered a candidate pair to be functionally related if they map to any functionally related pairs in STRING. If not a single pair they map to is functionally related, we further checked if the proteins they map to both have at least 5 other functionally related proteins in STRING, indicating that their functional associations are well studied and the lack of a functional interaction is not due to experimental challenges or low scientific interest. We considered any pair fitting these criteria unlikely to interact and included it in our pool of negative controls. We found 329,824 PDB chain pairs that can be used as negative controls. Similar to what we did for the **positive PPI set**, we clustered these pairs by placing pairs AB and A'B' into the same cluster if (1) A belongs to the same MMseqs cluster (--min-seq-id 0.3 -c 0.8 --cov-mode 0) as A' and (2) B belongs to the same MMseqs cluster as B'. The 329,824 negative control pairs represented 230,932 clusters, nearly 10 times more than the number of clusters in the **positive PPI set**. Because we mixed positive controls and negative controls at equal ratio to train our AI networks, having too many negative control pairs is not useful. Thus, we downsampled the negative controls to obtain 100,000 clusters comprised of 142,836 pairs, resulting in a **negative PPI set**.

#### M2.2. DPAM-IPR: a Domain Parser for AF2 Models utilizing InterPRo data

To expand the training dataset, we utilized interacting domains from AF2 models in AFDB. We recently developed a Domain Parser for AF2 Models (DPAM) (*16*). DPAM uses inter-residue distance matrices, predicted aligned error (PAE) matrices, and homology to previously classified structural domains in the Evolutionary Classification of protein Domains (ECOD (*17*), **Figure S4**) database to segment AF2 models domains. Integrating structural (distance and PAE matrices) and evolutionary evidence (ECOD domains) allowed us to find domains representing the structural and evolutionary units of proteins. The drawback of this pipeline is that it is time-consuming to find similar ECOD domains via sequence (HHsuite (*18*)) and structure (Dali (*19*)) searches. To analyze the massive amount of AFDB models, we needed a much faster tool. We therefore relied on homology evidence in the InterPro database (*20*). This shortcut is possible because AFDB contains models of UniProt proteins, which are all annotated with homologous domains from various databases in InterPro.

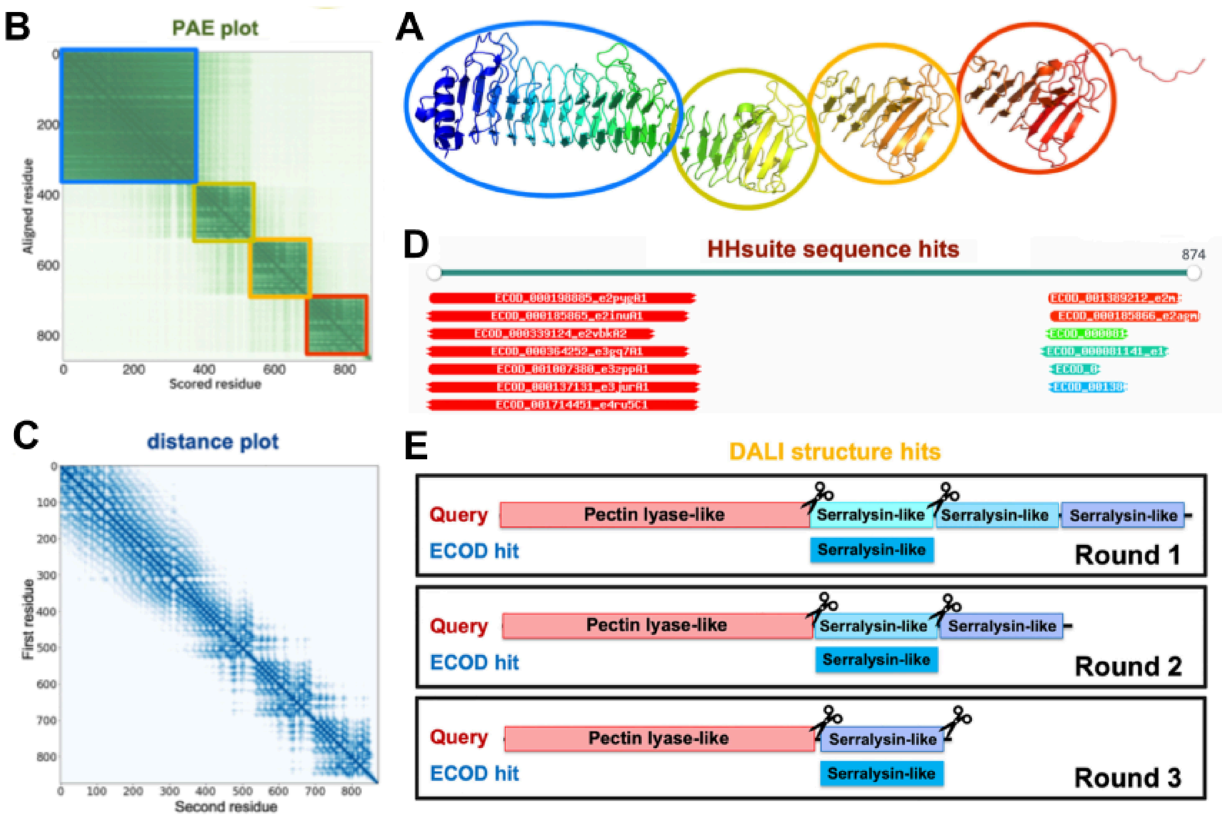


**Figure S4. Evidence to parse an AF2 model into globular domains. (A)** An example AF2 model (UniProt accession: Q9ZFH0). **(B)** A PAE plot. **(C)** A inter-residue distance plot. **(D)** Similar sequences detected by HHsuite. **(E)** Similar structures detected by Dali.

Compared to the homologous structural domains in ECOD detected by the sensitive sequence profile and structure comparisons, relying on domain annotations in the InterPro database has three potential caveats. First, these domains are annotated by aligning single sequences to the sequence profiles in various databases, an approach not as sensitive as HHsuite or Dali in detecting remote homology. As a result, more domains might be missed based on such homology evidence. However, based on our previous comparison of ECOD and Pfam (one major database used by InterPro), the fraction of missed domains should be below 10%. Given the massive scale of AFDB models, missing a small fraction of domains will not prevent us from identifying a large number of interacting domains to train AI networks. Second, domain classification hierarchies hosted by InterPro mostly consist of sequence-based domains. Without knowing the 3D structures, such domains frequently lack accurate boundaries and are particularly problematic when the evolutionary units consist of discontinuous segments in sequence. We reason that such problems with sequence-based domains can be alleviated by combining domains annotated by multiple resources in the InterPro database, as well as integrating the InterPro domains with structural features (distances and PAEs). Third, we previously found that sequence domains defined in the InterPro database sometimes contain multiple independent folding units (structural domains). Failing to split such sequence domains into smaller units will reduce the number of interacting domains we can find from AFDB models but will not jeopardize the quality of our dataset of DDIs: the interactions of sequence domains consisting of multiple structural units probably resemble PPIs even better.

Building on our experience developing DPAM, in this work, we developed a new method: DPAM-IPR (IPR stands for InterPro) to segment AFDB models into evolutionary and functional units with more precise domain boundaries. We created a benchmark set to evaluate the performance of DPAM-IPR as we optimize its parameters. We took the proteins we previously analyzed using DPAM (*16*, *21*). We reasoned that splitting domains defined by DPAM should be penalized, but merging two DPAM-defined domains should not be penalized because the conserved evolutionary units we want to find may contain multiple DPAM domains. However, to prevent DPAM-IPR from under-splitting domains, we calibrated to PDB entries. We reasoned that experimental constructs in PDB entries designed by experts should include the complete functional and evolutionary units. Thus, we penalize against merging regions included in the experimental constructs with those that were excluded. We mapped proteins previously analyzed by DPAM to PDB entries by BLAST (*22*) (e-value < 0.00001) and identified hits that can be nearly fully aligned to the query (unaligned residues at both termini < 10; total unaligned residues < 0.2 x hit length). We reasoned that the query region aligned to such a hit and the region not aligned should belong to different domains; if both aligned and un-aligned regions contain more than 100 residues, we used such information to optimize DPAM-IPR parameters. As a result, 100,261 AFDB entries that were 1) previously analyzed by DPAM and 2) partially mapped to PDB chains were used as the **DPAM-IPR benchmark set**.

From InterPro, we considered “Domains”, “Families”, and “Homologous superfamilies” as domains, and we used domains annotated by PFAM (*23*), CATH-Gene3D (*24*), SSF (*25*), SMART (*26*), PROFILE (*27*), and CDD (*28*) databases as homology-based evidence for domain parsing. From the 100,261 **DPAM-IPR benchmark** proteins, CATH-Gene3D, PFAM, SSF, PROFILE, SMART, and CDD annotated 197061, 180705, 151867, 104875, 97285, and 95886 domains, respectively. Because PROFILE, SMART, and CDD annotate far fewer domains than other databases, we used the following four sources of homology evidence: CATH-Gene3D domains, PFAM domains, SSF domains, and other domains (a merged set of PROFILE, SMART, and CDD domains).

Similarly to DPAM, DPAM-IPR parses domains based on the probability that a given residue pair belong to the same domain, which we denoted as $P_{COMB}$. This probability is calculated as the weighted geometric mean of probabilities for residue pairs to be in the same domain based on different types of evidence. We used the following formula $P_{COMB} = P_{DIST}^{0.25} \cdot P_{PAE}^{0.25} \cdot P_{HOMO}^{0.5}$, where $P_{DIST}$, $P_{PAE}$, and $P_{HOMO}$ are the same-domain probability according to inter-residue distance, PAE, and homology evidence. $P_{DIST}$ and $P_{PAE}$ are calculated based on regression analysis of benchmark ECOD domains as described in our DPAM paper (*16*). The weights (0.25, 0.25, and 0.5) in the above formula were not optimized but place equal weight (0.25 + 0.25 and 0.5) on structural and homology evidence.

**Table S1. Converting homology evidence to same-domain probability.**

| PFAM | CATHGENE3D | SSF | OTHERS | Prob | PFAM | CATHGENE3D | SSF | OTHERS | Prob |
|---|---|---|---|---|---|---|---|---|---|
| rest | same | same | same | 0.915 | same | same | different | rest | 0.405 |
| same | same | rest | rest | 0.885 | rest | rest | same | different | 0.368 |
| rest | rest | same | same | 0.885 | same | different | same | rest | 0.362 |
| same | same | same | same | 0.874 | different | rest | same | rest | 0.339 |
| same | rest | same | same | 0.864 | same | rest | different | rest | 0.333 |
| rest | same | rest | same | 0.844 | same | rest | rest | different | 0.316 |
| different | same | rest | same | 0.836 | rest | different | same | different | 0.31 |
| same | same | same | rest | 0.835 | different | different | same | same | 0.301 |
| rest | same | different | same | 0.832 | rest | different | different | same | 0.3 |
| rest | same | same | different | 0.823 | rest | same | rest | different | 0.291 |
| same | rest | rest | same | 0.816 | different | different | rest | same | 0.264 |
| same | same | rest | same | 0.813 | different | different | same | rest | 0.26 |
| rest | rest | rest | same | 0.803 | same | different | rest | same | 0.252 |
| rest | same | same | rest | 0.798 | different | rest | same | different | 0.25 |
| same | rest | different | same | 0.784 | rest | rest | rest | rest | 0.237 |
| different | same | same | same | 0.774 | same | different | rest | rest | 0.229 |
| rest | same | rest | rest | 0.768 | same | rest | same | different | 0.202 |
| same | same | different | same | 0.754 | different | different | same | different | 0.175 |
| rest | different | same | same | 0.738 | same | different | same | different | 0.17 |
| different | rest | same | same | 0.698 | same | rest | different | different | 0.145 |
| different | rest | rest | same | 0.687 | same | different | rest | different | 0.138 |
| different | same | same | different | 0.684 | different | rest | rest | rest | 0.138 |
| different | same | rest | different | 0.683 | different | different | different | same | 0.136 |
| same | same | same | different | 0.682 | rest | different | rest | rest | 0.101 |
| different | same | same | rest | 0.675 | rest | rest | different | rest | 0.101 |
| same | same | rest | different | 0.658 | rest | rest | rest | different | 0.092 |
| different | same | rest | rest | 0.657 | different | rest | rest | different | 0.09 |
| different | same | different | same | 0.635 | different | rest | different | rest | 0.084 |
| different | rest | different | same | 0.633 | rest | rest | different | different | 0.08 |
| rest | same | different | different | 0.604 | rest | different | different | rest | 0.077 |
| same | rest | rest | rest | 0.604 | different | rest | different | different | 0.075 |
| same | same | different | different | 0.567 | same | different | different | rest | 0.071 |
| rest | rest | same | rest | 0.566 | same | different | different | same | 0.059 |
| rest | same | different | rest | 0.566 | different | different | rest | rest | 0.059 |
| same | different | same | same | 0.563 | rest | different | rest | different | 0.051 |
| rest | different | rest | same | 0.506 | different | different | different | rest | 0.047 |
| rest | rest | different | same | 0.5 | same | different | different | different | 0.044 |
| different | same | different | rest | 0.496 | rest | different | different | different | 0.044 |
| same | rest | same | rest | 0.486 | different | different | rest | different | 0.029 |
| different | same | different | different | 0.466 | different | different | different | different | 0.026 |
| rest | different | same | rest | 0.411 | | | | | |

We analyzed how InterPro domain annotations should be converted to $P_{HOMO}$. Based on each of the four sources of homology evidence mentioned above, residue pairs in a protein may be in three categories: 1) belonging to the same domain, 2) belonging to different domains, and 3) the rest, i.e., including residues not assigned to any domain. We wondered how these states could be converted to $P_{HOMO}$. We partitioned residue pairs in proteins from the **DPAM-IPR benchmark set** into categories according to the four sources of homology evidence. We estimated the expected $P_{HOMO}$ for each category as the fraction of residue pairs in this category that belong to the same domains according to DPAM, and the results are shown in **Table S1.** As expected, $P_{HOMO}$ for a pair of residues is positively correlated with the number of databases that suggest them to be in the same domain.

Similar to DPAM, DPAM-IPR partitioned residues into domains by agglomerative clustering, grouping residues with high $P_{COMB}$ into the same domains. Briefly, we partitioned residues in a protein into non-overlapping and consecutive segments of 5 residues. We computed the average $P_{COMB}$ for every pair of segments. Initially, each 5-residue segment is treated as a candidate domain. Then, we sorted segment pairs reversely by $P_{COMB}$; we iterated through the list of segment pairs and progressively merged candidate domains linked by a pair of segments satisfying the following criteria: $inter\_P_{COMB} \cdot cutoff_M \geq min(intra1\_P_{COMB}, intra2\_P_{COMB})$ and $P_{COMB} > cutoff_P$. In the above formula, $intra1\_P_{COMB}$ is the average $P_{COMB}$ for segment pairs within the first candidate domain; $intra2\_P_{COMB}$ is the average $P_{COMB}$ for segment pairs within the second candidate domains; $inter\_P_{COMB}$ is the average $P_{COMB}$ for segment pairs between the two candidate domains; $cutoff_P$ and $cutoff_M$ are two parameters to be optimized.

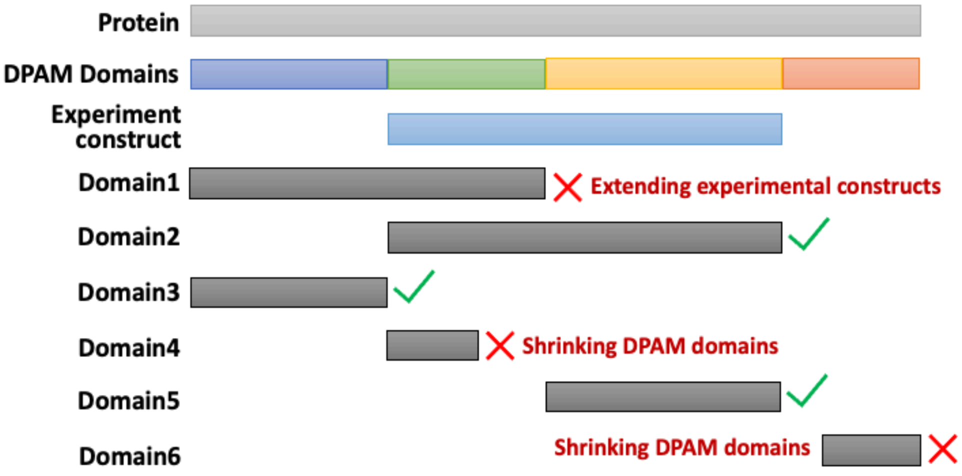


**Figure S5. Rationale to use the DPAM-IPR benchmark set for parameter optimization.** DPAM domains define the minimal boundaries while the experimental constructs define the maximal boundaries. Shrinking DPAM domains or extending experimental constructs will be penalized, while extending DPAM domains or shrinking experimental constructs will be tolerated.

Using the DPAM-IPR benchmark set, we performed a grid search to find the optimal $cutoff_P$ and $cutoff_M$. Proteins in this set were, on the one hand, segmented into structural domains by DPAM. On the other hand, at least a part of each protein was used in an experimental construct to solve a 3D structure. We reasoned that DPAM domains define the minimal boundaries, whereas the experimental constructs mark the maximal boundaries for the domains we want to find (**Figure S5**). Therefore, we considered residue pairs in the same DPAM domains to be true same-domain pairs; we considered a residue within an experimental construct and another residue outside this construct as true different-domain pairs. For $cutoff_P$ in the range of 0.2–0.8 (grid size: 0.1) and $cutoff_M$ in the range of 1.0–2.0 (grid size: 0.1), we computed the fraction of true same-domain residue pairs correctly predicted by DPAM-IPR (TPR) and the fraction of true different-domain residue pairs correctly predicted by DPAM-IPR (TNR). We chose the $cutoff_P$ and $cutoff_M$ values that resulted in the highest sum of TPR and TNR, i.e., 0.4 and 1.6.

After agglomerative clustering, we kept domains with at least 30 residues, and we further refined the boundaries of these domains. We attempted to shorten or extend each domain's boundary based on the contacts of each residue (other residues that are less than 6Å away in 3D space but more than 6 residues away in sequence) in this domain. If a residue at the domain boundary had more contacts out of this domain than within this domain, we excluded it; in contrast, we added a residue to extend the domain boundary if it had more contacts with residues within this domain than those outside this domain. Starting from residues at both ends of a domain, we repeated this procedure until no residues were removed or added.

#### M2.3. Pairs of interacting domains derived from multi-domain AFDB models

Equipped with DPAM-IPR, we set out to identify interacting domains from AFDB models (**Figure S6**). We downloaded the pre-clustered AFDB50 set from the Steinegger lab at: https://foldseek.steineggerlab.workers.dev/afdb50.tar.gz, which contains 53.7M models. We downloaded the archived InterPro data released around the same time as most of the AFDB models, from which we extracted "Domains", "Families", and "Homologous superfamilies" in these proteins annotated by PFAM, CATH-Gene3D, SSF, SMART, PROFILE, and CDD databases. We found 12.4M multiple domain proteins that contain at least two domains whose overlap is less than half the size of either domain. We applied DPAM-IPR to parse these 12.4M AFDB models into domains, resulting in 28.4M domains. We identified inter-residue contacts within each protein using the following criteria: 1) separated by at least 6 residues in between and 2) with minimal inter-residue distance < 6Å. We found 1.6M pairs of interacting domains using the following criteria: 1) average PAE for residues in either domain less than 8Å; 2) average PAE for residue pairs between the two domains less than 8Å; 3) more than 25 inter-domain contacts; 4) the number of inter-domain contacts higher than the numbers for intra-domain contacts in both domains. These criteria ensured that the 3D structures of both domains, as well as their interactions, are confidently predicted.

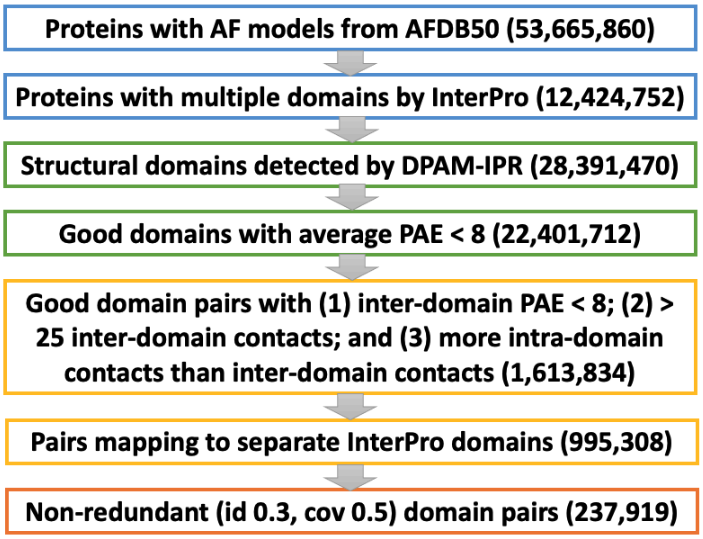


**Figure S6. Our procedure to prepare a positive training dataset with interacting domains extracted from multi-domain proteins in AFDB.**

We were concerned that some of these domain pairs might belong to the same evolutionary units, and their coevolutionary signals and interactions could be stronger than true PPIs. To remove such pairs, we filtered the 1.6M domain pairs to keep only those mapping to separate InterPro domains. Specifically, for each pair of interacting DPAM-IPR domains, we required that 1) an InterPro domain show over 50% overlap with the first domain and a 0% overlap with the second; 2) a different InterPro domain show 0% overlap with the first domain and over 50% overlap with the second. Nearly 1M domain pairs passed this filter, and they constitute our DDI training set. We manually inspected several dozens of DDIs found by our procedure, and find that they closely resemble interactions between proteins (**Figure S7**). We clustered these DDIs into 238k clusters at 30% sequence identity cutoff by placing pairs AB and A'B' into the same cluster if (1) A belongs to the same MMseqs cluster (--min-seq-id 0.3 -c 0.8 --cov-mode 0) as A' and (2) B belongs to the same MMseqs cluster as B'. We term the resulting training dataset the **positive DDI set**.

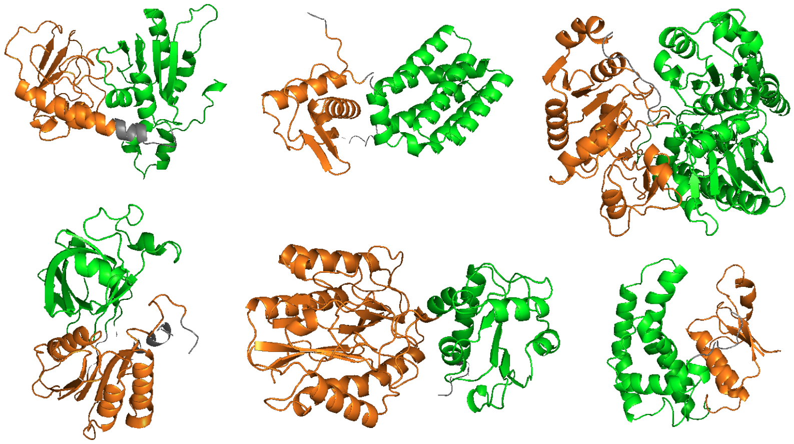

**Figure S7. Random examples of interacting domain pairs from AFDB models found by our pipeline.**

We also assembled a **negative DDI set** (**Figure S8**) by randomly pairing domains from the **positive DDI set**. Such random pairs need to be from the same organism but show no evidence of interacting. We analyzed the taxonomy information for AFDB models in the **positive DDI set** and focused on 211,995 sequences from 502 taxa in this set, where each taxon has more than 100 sequences. We mapped these sequences to entries in the STRING database using MMseqs, keeping only the 140,251 sequences (66%) that can be mapped to closely related STRING sequences (query or hit coverage > 0.75, identity > 50%). The STRING database focuses on genetic interactions, and it provides information for functionally related protein pairs. We found 2.4M pairs of AFDB models from the same taxa that are not functionally related (never mapping to genetically interacting protein pairs in STRING, and each protein has at least 5 other genetically interacting partners). We clustered these domain pairs as we did for the **negative PPI set**, and we randomly selected 240k clusters with 363k domain pairs to use as the **negative DDI set** for AI model training.

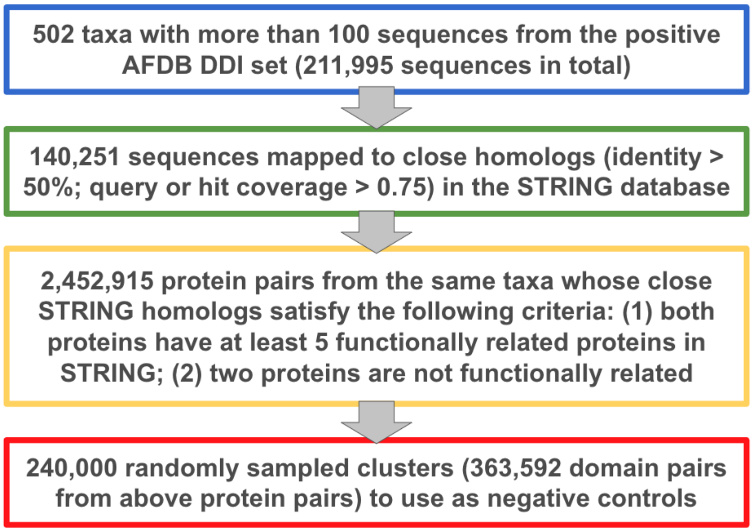


**Figure S8. Our procedure to prepare a negative training dataset by randomly pairing functionally unrelated domains from the positive DDI training set.**

#### M2.4. Preparing input data for the training datasets

For every entry in the training datasets, we prepared its paired MSA and 3D coordinates. For each binary protein complex in the **positive and negative PPI sets**, we extracted the 3D coordinates of the two chains from the corresponding PDB entries. For **the positive and negative DDI set**, we extracted the coordinates for each domain pair from the corresponding AFDB models. The extracted 3D structures for the **positive PPI** or **DDI** sets will be in the "bound conformation" determined in experimental structures or predicted by AF2, and the relative distances and orientations between residues in the two proteins/domains will be used to calculate the loss during training. In contrast, the two molecules in the negative sets, extracted from different PDB entries or AF2 models, will have random relative positions and orientations. This is not a problem because for the negative set, the distances between residues in different molecules were always assumed to be in the largest distance bin, while the relative orientations between residues in different molecules were not used.

Because many PDB chains (in the same or different PDB entries) have identical sequences, to save time for multiple sequence alignment (MSA) construction, we merged identical sequences and obtained a non-redundant set of protein sequences representing all PDB chains in the **positive and negative PPI set**. Similarly, we also obtained a non-redundant set of protein pairs. We built MSA for each sequence in the non-redundant set by searching against the UniRef30 database (obtained from https://wwwuser.gwdg.de/~compbiol/uniclust/2023_02/) with HHblits (*29*) (-realign_max 100000 -maxseq 1000000 -maxfilt 100000 -min_prefilter_hits 1000 -all). We found that the HHblits option “-all” is important. By default, HHblits filters the output MSAs at 90% sequence identity; turning this option on suppresses this behavior and allows all taxa to be present in the MSAs and more complete pairing of sequences from the same taxa to generate paired MSAs (pMSAs). We scored each hit in the output MSA of HHblits by its similarity to the query, and this similarity is evaluated by the total blosum62 score (*30*) of the aligned positions between the query and a hit. We selected a single hit showing the highest similarity to the query from each taxon. We further constructed a pMSA for each protein pair in the non-redundant set by concatenating sequences of the same taxa. We filtered the pMSAs by HHfilter (-id 95 -cov 50), and the resulting MSAs were used as inputs to train AI networks.

We built input MSAs for the **positive and negative DDI sets** in the same way, except that the “-all” option for HHblits was disabled due to disk space constraints. This reduced the depth of pMSA for the DDI sets, but we reasoned that it might be beneficial for models to learn to work with shallower MSAs. To replicate PPIs as much as possible in our **positive DDI set** made of domain pairs from the same protein, we built MSAs for each domain separately and generated the pMSA in the same fashion as the **positive PPI set**: instead of pairing domain sequences in the **positive DDI set** according to UniProt accessions, we selected the closest hit to the query in each taxon and paired them by taxon.

#### M3. Developing Artificial Intelligence networks for PPI prediction

##### M3.1. Simplifying the RoseTTAFold2 network and performance evaluation

We started from the current RF2 network and made a number of changes to streamline its architecture and speed up PPI prediction. **First,** we removed the 3D structure track. This was initially intended to simplify the network, but we later found that a better performance for PPI prediction can be achieved without the 3D structure track (see **M3.2**). **Second,** we reduced the number of attention-based blocks to 12 and the number of recycles to 3. **Third,** we removed loss functions designed to evaluate the accuracy of predicted 3D structures. In our final network, we only used loss functions for the following predictions: 1) randomly masked residues in the input MSA (masked token prediction loss, weight 1.0), 2) the relative distances between residues (distogram prediction loss, weight 2.0), 3) the relative orientations between residues (orientogram prediction loss, weight 2.0), and 4) disordered residues in experimental structures (disorder prediction loss, weight 0.1). **Fourth,** we did not use any 3D structures as templates because we found that using such templates in training decreases the network's ability to distinguish true PPIs from random pairs (see **M3.2**). Instead, we initialized the 2D pairwise features from randomly generated coordinates. **Fifth,** we removed the code for handling symmetry, because we always used a 1:1 stoichiometry for PPI screening. We previously explored whether including the stoichiometry of obligate homo-oligomeric proteins in the process of PPI modeling by AF2 can improve the success rate of identifying their interacting partners. We found that providing the correct homo-oligomeric stoichiometry actually reduced the accuracy for PPI prediction, possibly due to that modeling the homo-oligomers introduced additional complications for the AF. Even more, modeling a complex with stoichiometries other than 1:1 would greatly increase the inference time and required GPU memory. **Sixth,** we revised the data loader for model training to process different types of training data (monomers, PPIs, and DDIs). We named the revised network **RF2-ppi**, and we deposited this network and the trained weights to GitHub at: https://github.com/CongLabCode/RoseTTAFold2-PPI.

In order to determine the best training routine and model architecture, we randomly selected 3,600 **positive control** pairs and 36,000 **negative control** pairs and used them to monitor the performance of different versions of **RF2-ppi**. We referred to these selected sets as the **positive benchmark set** and **negative benchmark set**. We used precision and recall curves to evaluate **RF2-ppi**'s ability to distinguish the **positive benchmark** set from the **negative benchmark** set. We converted **RF2-ppi** outputs (distograms of inter-residue distances for residue pairs between two proteins) to the interaction probability between proteins (see **M5.3**), and we ranked the protein pairs in the benchmark sets reversely by the interaction probability and counted the numbers of true positives and false positives at each rank, allowing us to obtain a precision and recall curve for performance evaluation. To represent the low signal-to-noise ratio associated with *de novo* PPI screens, we split the **positive benchmark** set with 3,600 true PPIs into 100 parts and calculated a precision versus recall curve for each part against all the negative pairs (positive: negative = 36: 36,000). In order to utilize all the benchmark data for performance evaluation, we averaged the numbers of true positives over the 100 parts at each score cutoff to compute the overall precision and recall.

#### M3.2. Testing training dataset and model architectures

We studied the impact of three different types of training datasets, monomeric PDB chains, DDIs (positive and negative), and PPIs (positive and negative) on training **RF2-ppi** to distinguish true human PPIs from random pairs. We expected that the monomeric PDB chains (MONO set) are not as crucial for our task as the DDI and PPI set, and thus, we relied on the training set for RF2 made of 1) single chains from PDB entries released prior to Aug 2021 and 2) AF2 models for UniProt50 representatives. The DDI and PPI training datasets were newly prepared for this study as described above. For both the DDI and PPI sets, we mixed the interacting (positive) and non-interacting (negative) pairs at 1:1 ratio; an equal ratio was chosen to ensure the network would not overpredict or under-predict PPIs. For the MONO set, we took crops made of 256 continuous residues in the protein sequences. For the DDI or PPI sets, we took crops made of 256 residues near the DDI or PPI interfaces.

We were concerned that a significant fraction of the PPI training set are pairs of human proteins, and including them might appear to result in good performance due to information leakage. Therefore, we first excluded human homologs from the PPI set (see **M2.1**), and tested whether training **RF2-ppi** with the **entire PPI set** and the **filtered PPI set** show different performance in distinguishing true human PPIs from false ones. We used 12,800 training examples per epoch and randomly sampled 1 example per cluster. After training for 60 epochs, we found that the two training sets did not result in much difference in **RF2-ppi**'s performance (**Figure S9**), suggesting that the ability for **RF2-ppi** to distinguish true PPIs from random pair is not a result of memorizing the training data. Instead, such a network can learn the fundamental rules about residue-residue interaction, protein folding, and coevolution. Although this analysis suggested that information leakage will not be a problem in training **RF2-ppi**, we still relied on the **filtered PPI sets** in our study.

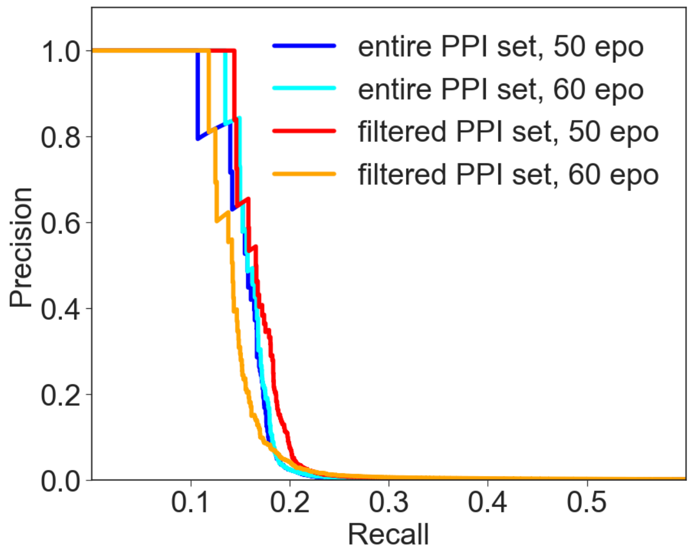


**Figure S9. Precision versus recall curves for RF2-ppi trained by the entire PPI sets (positive: negative = 1:1) or the filtered PPI sets (positive: negative = 1:1) where pairs of human protein homologs were removed.**

We next tested the performance of different training datasets and their combinations, including 1) the MONO set, 2) the DDI set, 3) the PPI set, 4) the MONO and DDI sets mixed at equal ratios, 5) the MONO and PPI sets mixed at equal ratios, 6) the DDI and PPI sets mixed at equal ratios, 7) the MONO, DDI, and PPI sets mixed at equal ratios. After 60 epochs, the performance of **RF2-ppi** trained with different datasets is shown in **Figure S10**. **RF2-ppi** showed terrible performance when training with the MONO set alone, because such training increased RF2-ppi's tendency to predict a high interaction probability for non-interacting pairs as well. It is worth noting that the poor performance of the MONO set could be related to the fact that we took continuous crops of 256 residues, so the model mostly saw interactions within domains. If we also took discontinuous crops from the MONO set, we might reach better performance by training with the MONO set alone. The DDI set alone showed better performance than the PPI set, and the best performance was achieved when the DDI set was combined with the PPI set (red curve in **Figure S10**), the MONO set (green curve in **Figure S10**), or both of them (lime curve in **Figure S10**).

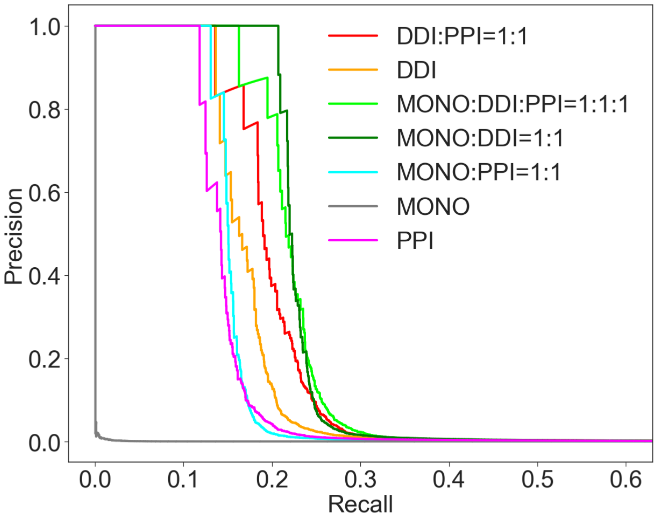


**Figure S10. Precision versus recall curves for RF2-ppi trained with different datasets and their combinations for 60 epochs.**

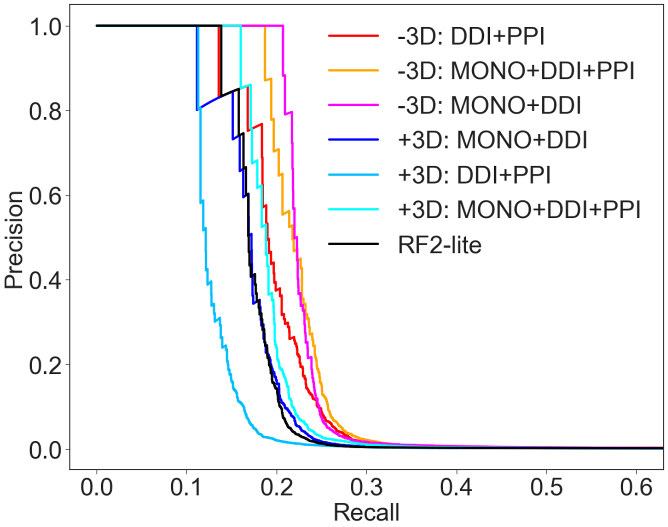


**Figure S11. Adding 3D structure features to RF2-ppi decreases its performance after training on various datasets for 60 epochs.**

We tested various modifications to the **RF2-ppi** architecture to improve its performance. We tried to add the 3D structure track and the loss functions associated with 3D structure prediction accuracy, resulting in a network similar to RF2-lite (*31*), a simplified version of RF2 with 8 attention-based blocks instead of 36 for full RF2. We recently used RF2-lite to identify PPIs in bacterial pathogens (*32*). Surprisingly, the addition of 3D structures to the network decreased its performance regardless of what training datasets were used (**Figure S11**). Similarly, RF2-lite (black curve in **Figure S11**) shows worse performance than **RF2-ppi** without the 3D structure track. RF2-lite also shows worse performance than the **RF2-ppi** network with 3D features (RF2-ppi+3D) trained on the combination of the MONO, DDI, and PPI sets for 60 epochs. RF2-lite was trained for 300 epochs on the exact MONO set and an older version of PPI set (without removal of human homologs). The superior performance of RF2-ppi+3D trained with much fewer epochs also demonstrates the power of the DDI training set.

After 60 epochs, it was not clear whether or not the addition of the PPI set to the combination of MONO and DDI sets would result in a better **RF2-ppi** network. With a 3D structure track, adding the PPI set on top of the MONO and DDI sets clearly improved its performance; however, we observed a slightly opposite trend for **RF2-ppi** without 3D structure features. We thus continued to train **RF2-ppi** without a 3D structure track for more epochs, and we found that in later epochs, training with the combination of MONO, DDI, and PPI sets showed better performance than training with the combination of MONO and DDI sets. We therefore chose to focus on the combination of all three sets for further explorations. We tested whether changing other hyperparameters of the **RF2-ppi** network would improve its performance, such as a larger crop size, a larger number of sequences in the seed MSA, and a larger number of sequences in the extended MSA. While these modifications significantly increased the time required for training and inference, they did not show visible improvement over the original crop size (256), seed MSA depth (128), and extended MSA depth (1024) that we inherited from RF2. Thus we did not change these parameters.

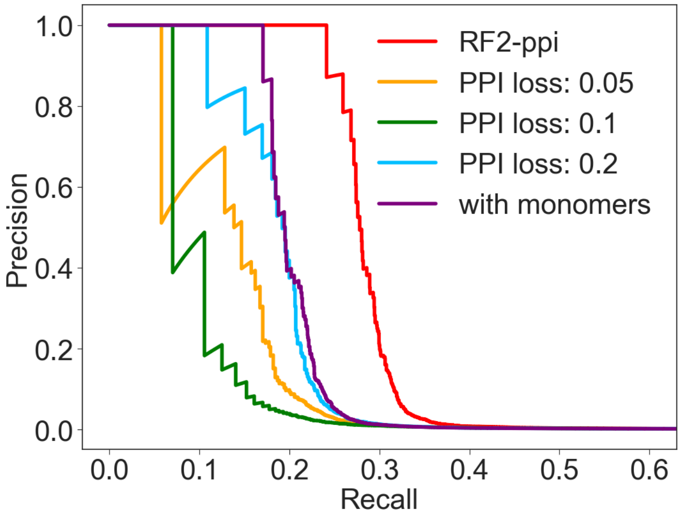


**Figure S12. Directly predicting whether two proteins/domains interact and adding a loss function for that decrease the performance of RF2-ppi. Using the monomer structures during training and inference also decreases the performance of RF2-ppi (purple curve).**

We also tested if directly predicting whether two proteins or domains should interact and adding a loss function for that (PPI loss) would help **RF2-ppi** to distinguish true interactions from false ones. We made such predictions based on the maximal interaction probabilities for residues between two molecules, and the inter-residue interaction probabilities based on the sum of probabilities for distogram bins below 12Å. We started with the **RF2-ppi** network trained without the PPI loss for 80 epochs, added PPI loss at different weights, and resumed the training. We found that adding a PPI loss significantly reduced the performance of RF2-ppi (**Figure S12**) in distinguishing true interactions from false ones at a signal-to-noise ratio of 1:1000. This is likely related to the fact that we trained the network at a signal-to-noise ratio of 1:1, and adding the PPI loss pushed the network to adapt to this much higher signal-to-noise ratio, making it less suitable for *de novo* PPI screens at very low signal-to-noise ratio.

We wondered whether providing the monomeric structures of potential interacting partners would help **RF2-ppi** to distinguish them from random pairs. During training, we extracted the monomer structures from the complex structure (i.e. bound confirmation) for each interacting protein/domain pair to initialize the pairwise features, i.e. distance and relative angles between residues features (distograms and orientograms) within each protein/domain. During validation, we used the predicted 3D structures of individual human proteins in AFDB to initialize the pairwise features. However, this attempt resulted in worse performance (purple curve in **Figure S12**). The lack of success is possibly related to the fact that we used the "bound conformations" for monomers during training, significantly simplifying the task for the AI network. Furthermore, in real applications, such bound conformations are not available and so providing these during training may bias the network against unbound conformations of the monomers. We continued to train the **RF2-ppi** network with the combination of MONO, DDI, and PPI sets for more epochs. We found the performance stopped improving after 140 epochs, and thus we took the model at 140 epochs as our final model (**Figure S13**).

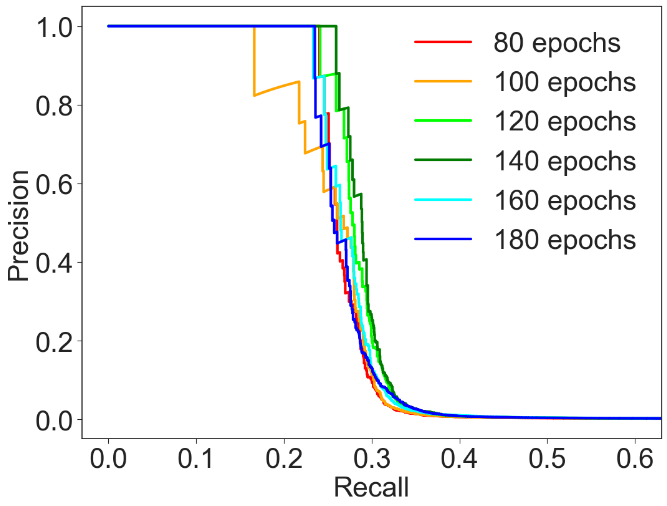


**Figure S13. Performance of RF2-ppi as a function of the number of training epochs.**

#### M3.3. Comparing the performance of RF2-ppi against other tools

Compared to our previous AI networks for proteome-wide PPI screens, i.e., the RF-2track network (*33*) used for yeast and the RF2-lite (*31*) network used for human pathogens, **RF2-ppi** shows much better performance (**Figure S14A**) in distinguishing the **positive benchmark set** from the **negative benchmark set**. We were concerned that this improved performance might be related to the fact that we used these benchmark sets to guide our decisions in selecting the training datasets and network architecture. Thus we tested the performance of these AI networks on 3,600 proteins selected from the **positive validation set** and 36,000 newly selected proteins from the large **negative control set** (negative validation set). The caveat of the **positive validation set** is that PPIs in this set are not as strongly supported and a remarkable fraction of them might be non-specific interactions (false positives) from experimental studies; indeed, our networks showed much worse recall on this validation set than the benchmark set. Nevertheless, these validation sets not used in the process of our method development clearly demonstrated the superior performance of **RF2-ppi** over the previous networks (**Figure S14B**).

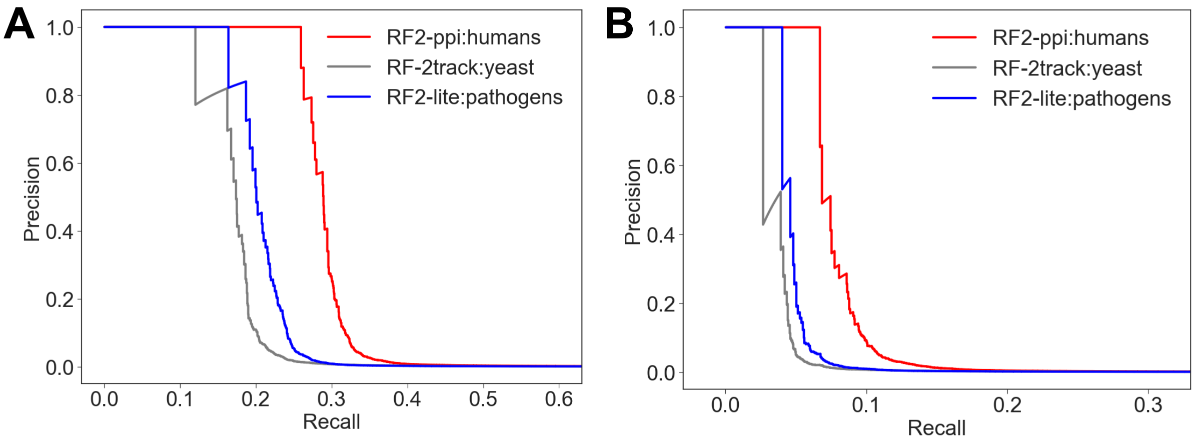


**Figure S14. RF2-ppi shows significantly better performance than our previous tools for distinguishing (A) positive controls from the negative controls, and (B) positive validation set from the negative controls.**

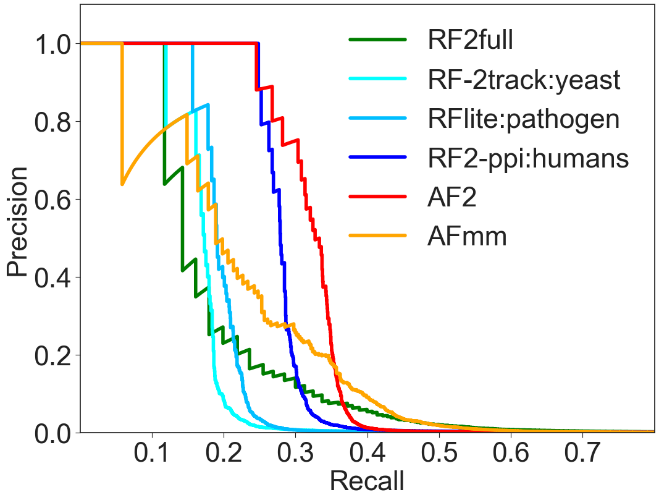


**Figure S15. Comparison of performance between RF2-ppi and other more complicated AI networks for protein/complex structure modeling in distinguishing true PPIs from false ones at a signal-to-noise ratio of 1:1000.**

Finally, we compared **RF2-ppi**'s ability in distinguishing true PPIs from false ones against the much slower AI networks, including the full version of RF2 (RF2full), AF2, and AFmm (**Figure S15**). At a signal-to-noise ratio of 1:1000, which approximates the situation of a *de novo* PPI screen, RF2-ppi's performance is only worse than AF2, and much better than RF2full or AFmm. Both AFmm and RF2full are trained with PPIs from PDB and optimized for predicting protein complex structures, and thus they might tend to over-predict interactions for random protein pairs and show worse performance when the noise level is high. However, when the noise level is low (for example, at 1:10), these more sophisticated AI networks do perform better than **RF2-ppi** (**Figure S16**) in identifying the true PPIs.

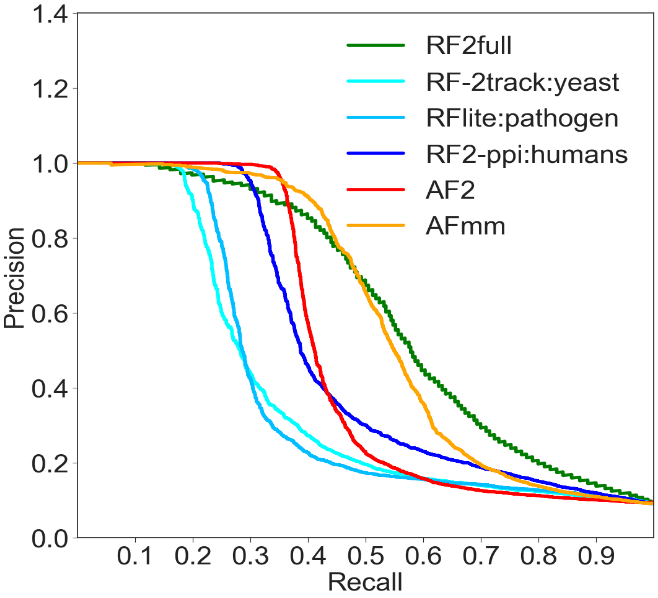


**Figure S16. Comparison of performance between RF2-ppi and other more complicated AI networks for protein/complex structure modeling in distinguishing true PPIs from false ones at a signal-to-noise ratio of 1:10.**

### M4. Generating multiple sequence alignments from genomic data

#### M4.1. Stage1: gathering genomic datasets for Eukaryotes

##### Gathering draft genomes from NCBI genome database

We used NCBI's command-line tool to gather genomic datasets and their accessory information from the NCBI genome and Sequence Read Archive (SRA) databases, following instructions provided at https://www.ncbi.nlm.nih.gov/datasets/docs/v2/download-and-install/. We collected information about eukaryotic genomes (till May 2023) using the following command: $ datasets summary genome taxon Eukaryota --exclude-atypical -as-json-lines. The “exclude-atypical” flag excludes the problematic assemblies detected by NCBI. From the resulting json file, we extracted the contig N50, scaffold N50, and the number of protein coding genes annotated from each genome. The N50 value represents the length of the contig or scaffold such that 50% of the entire assembly’s length is contained in contigs or scaffolds of that length or longer. N50 is a major indicator of genome assembly quality, and one cannot conveniently predict protein sequences from assemblies with a very low N50, because the coding sequences of many proteins might be split into different contigs or scaffolds. Thus, we only kept genomes with scaffold N50 or contig N50 above 50,000.

We extracted the TaxID of each genome assembly and assigned each assembly to a taxonomic lineage (phylum, class, order, family, genus, species) based on tables downloaded from the NCBI taxonomy database at: https://ftp.ncbi.nlm.nih.gov/pub/taxonomy/. We selected the best genome assembly for each species: if a species contains assemblies that are marked as “reference” by NCBI, we considered these “reference” assemblies as candidates; if not, all assemblies were regarded as candidates. From these candidates, we selected the assembly with the largest scaffold or contig N50 among those with assembly size above 0.9 · median assembly size for this species. As a result, we selected 9,337 genome assemblies representing 9,337 species, with the largest number of genomes from the Chordata, Arthropoda, and Ascomycota (**Table S2**). Data for the selected genomes were downloaded from NCBI using command-lines as follows: $ datasets download genome [assembly_accession] --include genome,protein,gff3.

**Table S2. Statistics of selected representative genomes for eukaryotic species.**

| Phylum | # genomes with proteomes | # genomes without proteomes | # genomes |
|---|---|---|---|
| Chordata | 357 | 1,820 | 2,177 |
| Arthropoda | 143 | 1,403 | 1,546 |
| Ascomycota | 938 | 1,773 | 2,711 |
| Streptophyta | 191 | 832 | 1,023 |
| Basidiomycota | 356 | 350 | 706 |
| Others | 466 | 708 | 1,174 |
| All | 2,451 | 6,886 | 9,337 |

Most of these genomes are not annotated with protein-coding sequences, and some are associated with only several protein-coding genes. Because complete eukaryotic genomes should at least encode several thousands of proteins, we considered genome assemblies with at least 4,000 protein-coding genes to be associated with proteomes. 2,451 of these 9,337 genomes are associated with proteomes. We used these proteomes to assist us in assembling and aligning orthologous sequences for human proteins from diverse eukaryotic species. It is important to note that these proteomes are not necessarily complete or of high quality, and thus we still attempted to directly predict protein sequences from the genomes that are associated with proteomes (see **M4.2**).

#### Gathering genomic and transcriptomic reads from NCBI SRA database

For other Eukaryotes without assembled genomes, we set out to use the unassembled genomic and transcriptomic reads from NCBI SRA database, which contained around 30 Petabytes of data in May 2023. First, we downloaded the metadata of these sequencing reads using the NCBI E-utility tools with the following command: $ esearch -db sra -query "Eukaryota[Organism] NOT txid9606[Organism:exp] NOT txid9612[Organism:exp] NOT txid6239[Organism:exp] NOT txid4522[Organism:exp] NOT txid9031[Organism:exp] NOT txid9544[Organism:exp] NOT txid7165[Organism:exp] NOT txid9913[Organism:exp] NOT txid7227[Organism:exp] NOT txid9823[Organism:exp] NOT txid10116[Organism:exp] NOT txid7955[Organism:exp] NOT txid4565[Organism:exp] NOT txid4530[Organism:exp] NOT txid4577[Organism:exp] NOT txid4513[Organism:exp] NOT txid3702[Organism:exp] NOT txid5833[Organism:exp] NOT txid4932[Organism:exp] NOT txid10090[Organism:exp]" | efetch -format runinfo. This command extracted all records from Eukaryotes excluding the 20 species as specified by their TaxIDs. Based on the information we obtained by browsing the SRA website, these 20 species are associated with the largest numbers of SRA datasets. Because assembled genomes are available for these 20 species, we did not need their genomic or transcriptomic reads. However, these 20 most frequent species constitute 75% of the SRA datasets, and excluding them significantly reduced the time to download the metadata for the rest.

From these metadata, we extracted information about each SRA dataset. We selected whole transcriptomic and whole genomic reads sequenced by illumina, which not only generate high-quality data but also constitute most of the SRA datasets. Specifically, we used the following criteria to select the desirable datasets based on their metadata. **Instrument:** HiSeq X Five, NextSeq 2000, Illumina HiSeq X Ten, Illumina Genome Analyzer II, Illumina HiSeq 1000, Illumina Genome Analyzer IIx, NextSeq 550, Illumina HiSeq 1500, Illumina HiSeq X, Illumina HiSeq 3000, HiSeq X Ten, NextSeq 500, Illumina NovaSeq 6000, Illumina HiSeq 4000, Illumina MiSeq, Illumina HiSeq 2000, or Illumina HiSeq 2500; **Library Source:** TRANSCRIPTOMIC or GENOMIC; **Library Strategy:** RNA-Seq, WGS, or OTHER; **Library Selection:** Oligo-dT, PolyA, other, other, unspecified, cDNA, RANDOM, PCR. We only focused on datasets of moderate sizes: small ones will not be sufficient to cover the entire genomes or transcriptomes; while extremely large ones will be a challenge for data storage and downstream processing. As such, we only used datasets with at least 100M bases; we discarded transcriptomic datasets with larger than 100G bases or genomic datasets with larger than 300G bases.

We also extracted the "**Organism Name**" of each dataset, and assigned it to a taxonomic lineage based on tables downloaded from the NCBI taxonomy database. We reasoned that the necessary amount of sequencing to cover the entire genomes or transcriptomes vary by taxonomic lineages. To derive lineage-specific cutoffs for the amount of sequencing data, we computed the median amount of genomic and transcriptomic data, respectively, for each phylum ($Gm_{ph}$ and $Tm_{ph}$), class ($Gm_{cl}$ and $Tm_{cl}$), order ($Gm_{or}$ and $Tm_{or}$), family ($Gm_{fa}$ and $Tm_{fa}$), and genus ($Gm_{ge}$ and $Tm_{ge}$). We computed the cutoff for the amount of genomic ($Gcut$) or transcriptomic ($Tcut$) data for a species as $Gcut = 0.95 \cdot min(Gm_{ph}, Gm_{cl}, Gm_{or}, Gm_{fa}, Gm_{ge})$ and $Tcut = 0.95 \cdot min(Tm_{ph}, Tm_{cl}, Tm_{or}, Tm_{fa}, Tm_{ge})$. For each species without genomic assemblies extracted from NCBI genome database, we selected the largest genomic dataset from SRA with bases less than 300G and above $Gcut$ and the largest transcriptomic dataset with bases less than 100G and above $Tcut$. As a result, genomic datasets for 23,733 species and transcriptomic datasets for 16,820 species were selected.

Due to the limitation in disk space, we only focused on species from two largest phyla that are relatively close to humans, Chordata (4,585 genomic datasets and 2,055 transcriptomic datasets, representing 6,053 species) and Arthropoda (4,697 genomic datasets and 4,792 transcriptomic datasets, representing 9,069 species). Other phylums that are close to humans, Echinodermata and Hemichordata, have very few (around 200) SRA datasets and thus were not used. We downloaded these SRA reads using sra-toolkit with commands like: $ fastq-dump [accession] --split-3  --skip-technical –fasta –quite.

#### M4.2 Stage2: assembling orthologs from draft genomes associated with proteomes

We processed 2,451 genomes (representing 2,451 species) associated with annotated protein sequences, i.e., proteomes. From each genome, we used multiple strategies to find candidate orthologous sequences for each human protein, including: 1) direct detection of orthologs from the proteome using human proteins as queries; 2) indirect detection of orthologs from the proteome using proteins of other model organisms as queries; 3) direct assembling from the genome using human proteins as queries; and 4) indirect assembling from the genome using proteins of other model organisms as queries. The two direct strategies were used for all genomes, while the indirect strategies using queries of other model organisms were only applied to species in two of the largest phylums, Arthropoda and Ascomycota.

The reference protein sets of *Drosophila melanogaster* and *Saccharomyces cerevisiae* (yeast), were used to facilitate ortholog detection and protein sequence assembly from Arthropoda and Ascomycota genomes, respectively. We reasoned that using reference proteins from more closely related species might increase the accuracy for sequence assembly and ortholog detection, because aligning more divergent sequences or distinguishing orthologs from paralogs between more divergent species might be challenging. The reference protein sets of *Drosophila* and yeast were further mapped to their human orthologs based on reciprocal best hit criteria (see **M4.3**); thus the candidate orthologs assigned to *Drosophila* or yeast proteins will be propagated to their human orthologs. The large evolutionary distance between human and *Drosophila*/yeast and the fact that humans have experienced several whole genome-wide duplications in relation to these lower Eukaryotes will conflate ortholog detection. Nevertheless, the fact that these best-studied model organisms have complete protein sets and accurate protein sequences will likely make it easier to find orthologs between them and humans.

##### Detecting orthologs from proteomes

To detect orthologs of a reference protein from a target proteome (**Figure S17A**), we first found homologs for each reference protein using BLAST, keeping hits with e-value < 0.0001 and identity > 0.5 · top-hit identity. BLAST may separate the alignment between the query and a hit into multiple **High Scoring Pairs** (HSPs, i.e., aligned segments between a query and a hit) if the hit experiences large insertions or deletions relative to the query, which is not uncommon for eukaryotic sequences due to alternative splicing. Thus, we designed routines to compare multiple HSPs and merge compatible ones. We merged pairs of HSPs if they satisfied the following criteria: 1) overlap in covered query residues by both HSPs is smaller than half of their coverage in the query; 2) overlap in covered hit residues by both HSPs is smaller than half of their coverage in the hit; 3) merging the two HSPs can extend the query coverage by both HSP by at least 30 residues. If two HSPs should be merged, in their overlapping region, we chose the one displaying higher sequence similarity (measured by total blosum62 scores) between the query and the hit. This HSP merging routine could combine several HSPs into one alignment between the query and a hit, and if there were more than one merged alignment for a hit, the one including the HSP showing the lowest e-value was kept. As a result, we obtained only one alignment for each query-hit pair.

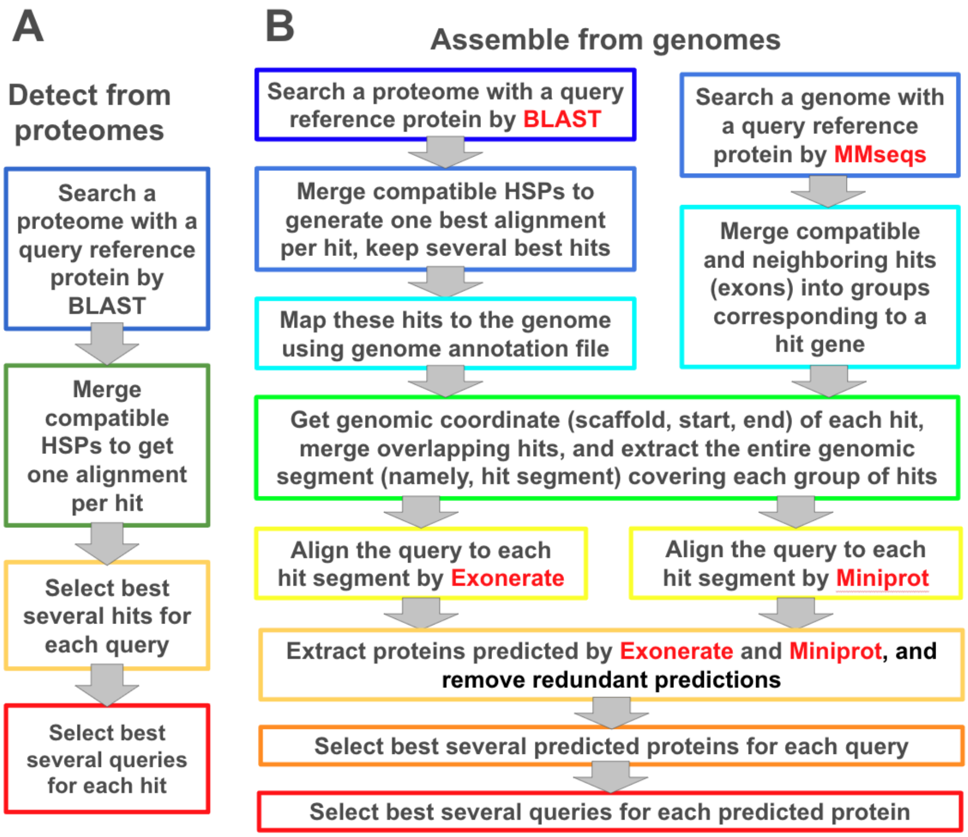


**Figure S16. Pipelines (A) to detect candidate orthologs for each reference protein from proteomes and (B) to assemble candidate orthologs directly from genomes.**

Among the hits for each query, we used a relaxed reciprocal best hit criteria to select candidate orthologs, allowing each query to have multiple best hits showing comparable similarities to the query. **First,** we found pairs of hits that overlap remarkably in the query residues they mapped to (overlap > 0.5 · query coverage by either hit), and we removed one of them, e.g., hit1, if it showed lower similarity to the query than the other hit, i.e., hit2 by the following criteria: 1) over the entire alignment, the total blosum62 score for hit1 · 1.5 < the total blosum62 score for hit2; 2) over the query residues covered by both hits, the average blosum score for hit1 · 1.1 < the average blosum score for hit2. **Second,** for each hit from a target proteome, we analyzed the query reference proteins aligned to this hit. We compared every pair of query proteins mapping to overlapping regions in the hit (overlap > 0.5 · hit coverage by either query) and removed the query showing lower similarity to the hit than the other, according to the same criteria as described above. All candidate orthologs were used to build the MSA for a query human protein and the closest hit was selected based on statistics calculated from the final MSA (see **M4.5**).

#### Assembling orthologous sequences from genomes

We were concerned with the quality and completeness of the annotated proteins in these draft genomes, and therefore, we designed a pipeline (**Figure S17B**) to directly assemble protein sequences from the genomes using a set of reference proteins (from human, *Drosophila*, or yeast). In this pipeline, we first identified the genomic segments encoding homologs of a query protein. On the one hand, we found homologs of a query by BLAST against the proteome. We merged the compatible HSPs to generate one alignment per hit as described above. We identified the best several hits for each query that satisfied the following criteria: 1) coverage to query by a hit is larger than the maximal query coverage among all hits; and 2) the total blosum62 score between the query and a hit is larger than the maximal total blosum62 score among all hits. We looked up where each selected hit mapped to in the genome based on the genome annotation file we downloaded from NCBI (GFF format) for each proteome, and these genomic regions (from the start site of a coding gene to the end site) were used as **homologous genomic segments**.

On the other hand, we directly found **homologous genomic segments** for each query by mapping it to the genome with MMseqs. We choose MMseqs due to its superior speed for aligning thousands of query proteins to a genome. For optimal speed, we prepared all query proteins in a reference species (human, *Drosophila*, or yeast) into a database using the “createdb” command in MMseqs. Similarly, we prepared another database with all the scaffolds from a target genome. We searched the database with query proteins against the one with scaffolds of a target genome using the following command: `$ mmseqs search [query_database] [genome_database] [output_prefix] [temporary_directory] --start-sens 3 --sens-steps 2 -s 7 -a 1 --threads 32`. MMseqs aligns each query protein to open-reading frames (ORFs) in a target genome, and these aligned ORFs likely correspond to exons or partial exons of a gene encoding a homologous protein to the query.

We processed the MMseqs output to group multiple ORFs aligned to a query into a potential protein-coding gene. The alignment between a segment in the query and a hit ORF can be considered as a HSP (high scoring pair). We kept HSPs with e-value < 0.001 and considered pairs of HSPs compatible to each other if they satisfied the following criteria: 1) their overlap in aligned query residues is shorter than half of the aligned query residues of both HSPs; 2) their overlap in aligned genomic bases is shorter than a quarter of the aligned genomic bases of both HSPs; 3) the hit ORFs are on the same strand of the same scaffold; 4) the genomic distance between the two hit ORFs is shorter than the 95% quantile of gene size (distance between the first and the last base in the coding sequence) in the target genome; and 5) the order of aligned regions in the query is consistent with the order of aligned regions in the hit ORFs (**Figure S18**).

Starting from the HSP showing the lowest e-value, we merged an HSP into a group if it is compatible with all existing HSPs in the group; otherwise, we started a new group with this HSP. Each HSP group represents a gene encoding a homolog to the query. We evaluated each HSP group's similarity to the query based on the total blosum62 score ($Sco_{HSPgrp}$) of the aligned residues in these HSPs and the total number of covered query residues by them ($Cov_{HSPgrp}$). We kept HSP groups showing $Sco_{HSPgrp} > 0.67 \cdot max(Sco_{HSPgrp})$ and $Cov_{HSPgrp} > 0.67 \cdot max(Cov_{HSPgrp})$. We extracted a minimal continuous genomic segment that completely covered the HSPs in each group, which represented a **homologous genomic segment** to the query.

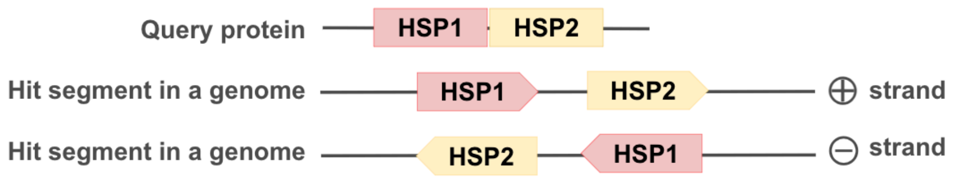


**Figure S18. HSPs that are consistent in their orders on the query protein and hit genome.**

We analyzed the **homologous genomic segments** for each query protein found by the above approaches. We merged segments 1) from the same strand of the same scaffold and 2) showing overlap in their genomic coordinates, the resulting merged segments were aligned to the query protein using exonerate (*34*) and miniprot (*35*). We parsed outputs of these tools to extract the coding sequences for homologs of the query. In contrast to MMseqs, exonerate and miniprot are aware of splicing sites, and thus they are able to align a protein sequence to multiple exons, skipping long introns.

However, we observed that both exonerate and miniprot still frequently split the alignment between a query and a homologous genomic segment into multiple HSPs covering different regions of the query protein. Therefore, we compared pairs of HSPs from each program and merged HSPs satisfying the following criteria. First, their overlap in aligned query residues is shorter than half of the aligned query residues of both HSPs. Second, the aligned regions in the target genome have no overlap in their genomic coordinates. Third, the arrangement of these HSPs on the genome are consistent with their order on the query protein: 1) if they are on the same scaffold, their distance should be than 200,000 bp and the order of the hits on the genome should be consistent to the aligned regions in the query protein (as in **Figure S18**); 2) if they are on different scaffolds, both of them should be near the end (< 20,000 bp away) of the scaffolds and consistent with the assumption that the two scaffolds can be concatenated to form the coding sequence of a homolog to the query protein (as in **Figure S19**). When two HSPs were merged, in the query regions covered by both HSPs, the HSP showing higher similarity between the query and the hit was taken.

In addition, we observed a decay in alignment quality towards the beginnings and ends of HSPs, which is likely a result of failing to find the true exons. To fix these problems, we calculated the average blosum62 score for each exon in each HSP, and if an exon at the edge (the beginning or the end) showed average blosum62 score lower than ⅔ of the scores for the exons in the middle, we shrunk the alignment and excluded this exon.

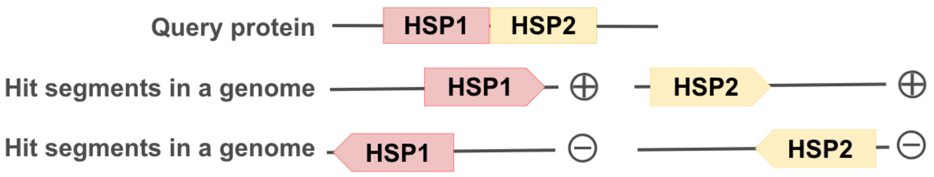


**Figure S19. HSPs that are consistent in their arrangement on the query protein and hit genomic segments from different scaffolds of a draft genome.**

Among the homologs for each query found by exonerate and exonerate, we used a relaxed reciprocal best hit criteria to select candidate orthologs as described above, allowing each query to have multiple candidate orthologs with comparable similarities. First, we found pairs of coding sequences with >50% overlap in the query residues they mapped to, and we removed one of them (sequence1) if it showed lower similarity to the query than the other (sequence2) by the following criteria: 1) over the entire alignment, the total blosum62 score of sequence1 · 1.5 < the total blosum62 score for sequence2; 2) over the query residues covered by both sequences, the average blosum62 score for sequence1 · 1.1 < the average blosum62 score for sequence2. Second, for pairs of homologous coding sequences of different queries showing >50% overlap in their genomic coordinates, we removed one of the queries (query1) if it showed lower similarity to the coding sequence than the other (query2) by the following criteria: 1) over the entire alignment, the total blosum62 score for query1 · 1.5 < the total blosum62 score for query2; 2) over the genomic positions that were mapped to by both queries, the average blosum62 score for query1 · 1.5 < the average blosum62 score for query2. The selected coding sequences for each query by these criteria were translated into protein sequences and considered as candidate orthologs of the query. All the candidates were used to build the MSA for each query human protein and the closest hit to the query was selected based on statistics calculated from the final MSA (see **M4.5**).

#### Comparing the "from proteome" and the "from genome" strategies

We wondered whether direct assembly of sequences from the genomes would help us to find the orthologous proteins or segments (e.g. an exon) that were missed in the set of annotated proteins. For each query, we examined its alignment to the best candidate ortholog (the hit showing the highest total blosum62 score over all aligned positions) found either by BLAST search against a proteome (the "from proteome" strategy) or by exonerate/miniprot search against a genomic segment (the "from genome" strategy). We partitioned the aligned positions by these two strategies into bins based on the blosum62 score between the query and hit residues. We found that to assemble orthologs from Chordata species for human proteins, the "from genome" strategy generated more aligned residues in each blosum62 score bin, suggesting that this strategy allowed us to assemble more complete sequences and obtain better alignments (**Figure S20A**). We observed the same trend for assembling sequences from Arthopoda species using *Drosophila* proteins as the reference (**Figure S20B**). However, the "from genome" strategy generated less aligned residues in each blosum62 score bin than the “from proteome” strategy when human proteins were used as reference to assemble orthologs from non-Chordata species (**Figure S20C**). Similarly, direct assembling of sequences from Ascomycota genomes using Yeast proteins as references also did not generate better alignments than the “from proteome” strategy (**Figure S20D**).

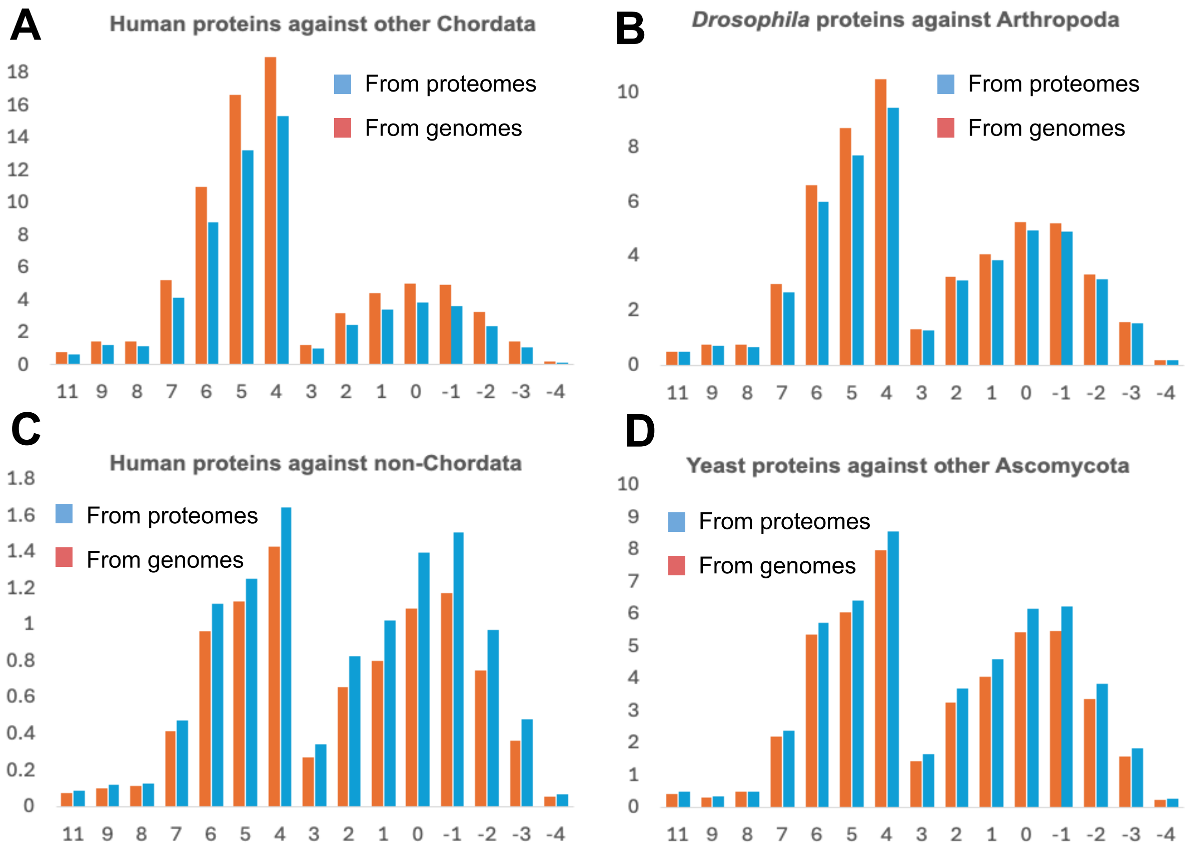


**Figure S20. The distribution of blosum62 scores for aligned positions between reference proteins and their orthologs detected from proteomes (blue bars) or assembled from genomes (red bars).** The X-axis is the blosum62 score between a pair of aligned residues, and Y-axis is the percentage of residue pairs. The two peaks correspond to the scores between identical residues (high blosum62 scores) and different residues (low blosum62 scores)

These results showed that when the reference protein set is closely related to the target genome, the “from genome” strategy is expected to obtain better alignments than the “from proteome” strategy. When the evolutionary distance between the reference and the target is large, such as between humans and non-Chordata species, assembling coding sequences in the target genomes by aligning them to the reference proteins is challenging. In contrast, the previously annotated proteins in these target genomes were likely based on RNA-seq evidence or homologs from more closely related species, resulting in a more complete set of annotated proteins and more complete protein sequences. These results suggest that direct assembling of protein sequences from draft genomes is an efficient strategy to obtain deeper MSAs for human proteins. However, for target genomes showing large evolutionary distances to humans, the proteomes of more closely related species should be used as references.

#### M4.3 Stage3: assembling orthologs from draft genomes

We gathered candidate orthologs assembled by different strategies in the previous stage. Some of these sequences were assembled using the *Drosophila* or yeast proteins as references. To assign them to human proteins, we assigned each *Drosophila* and yeast protein to its human orthologs (allowing multiple orthologs) based on the relaxed reciprocal best hit strategy described above (**Figure S17A**). We aligned each query human protein to its candidate orthologs using HMMER (*36*) with the following command: $ phmmer -o /dev/null --noali --notextw --incE 1e-2 -E 1e-2 --cpu 4 -A [output] [human_query] [all_candidate_orthologs]. Based on the output alignment for each human protein, we computed the similarity score ($Sim_{prot}$) for each candidate ortholog (hit) as the sum of positive blosum62 scores of aligned residues (gaps or aligned positions with negative blosum score did not contribute). For each query protein, we selected its best hit with the highest $Sim_{prot}$ from each species, genus, family, order, class, and phylum, and the similarity scores for these best hits were denoted as $Sim_{spe}$, $Sim_{gen}$, $Sim_{fam}$, $Sim_{ord}$, $Sim_{cla}$, and $Sim_{phy}$.

For each species processed in the previous stage, we generated a revised proteome to be used as a reference for orthologous sequence assembly in this stage. We built this revised proteome based on the following ideas: 1) we selected the best hit from this species for each human protein, if available; 2) if a best hit is not available in this species (possibly due to gene loss in evolution or in the process of genome assembly and annotation), we replaced it with the best hit from higher taxonomic ranks; 3) if the best hit in a higher taxonomic rank shows remarkably (1.2 times) higher similarity to the query, we used it to replace the best hit in this species. In practice, we computed preference scores for the best hits in this species and higher taxonomic ranks as: $Pref_{spe} = Sim_{spe} \cdot 1.2^5 + 0.5$; $Pref_{gen} = Sim_{gen} \cdot 1.2^4 + 0.4$; $Pref_{fam} = Sim_{fam} \cdot 1.2^3 + 0.3$; $Pref_{ord} = Sim_{ord} \cdot 1.2^2 + 0.2$; $Pref_{cla} = Sim_{cla} \cdot 1.2 + 0.1$; $Pref_{phy} = Sim_{phy}$. For each query protein and each proteome assembled in the previous stage, we selected the best hit with maximal preference score and added it to the revised proteome. These revised proteomes were used as references for assembling proteins from other draft genomes without annotated proteins in this stage.

For each draft genome (target) without annotated proteins, we found its closely related reference proteome based on taxonomic information. We went up the taxonomic ranks of a target genome, from genus to phylum, to find the rank with available reference proteomes in the same rank. At that taxonomic rank, we selected the reference proteome covering the largest number of human residues in the alignments made by phmmer (see above) between every human proteome and each reference proteome. Both the closely related reference proteome and the human proteome were used to assemble candidate orthologs from the target genome, using a pipeline (**Figure S21**) similar to the one used in the previous stage. However, since the target genome is not associated with a proteome, we could not use previously annotated proteins to help us to find the **homologous genomic segments** for a query protein.

We gathered the candidate orthologs assembled from each target genome using the human proteome or the selected, closely related, proteome as a reference. Since each of the additional reference proteomes was made of orthologs of human proteins, assembled sequences based on these references could be easily assigned as candidate orthologs of human proteins. We again aligned each query human protein to its candidate orthologs using HMMER. Based on the output alignments, we counted the number of residues in the human proteins that were covered by the orthologs assembled by different strategies. We found that for Chordata species, using human proteins as queries resulted in slightly (1.005 times) higher coverage of human protein residues by the assembled sequences. However, for non-Chordata species, assembling sequences with more closely related references resulted in remarkably (1.14 times) higher coverage over human proteins (i.e. more complete assembled proteomes) on average.

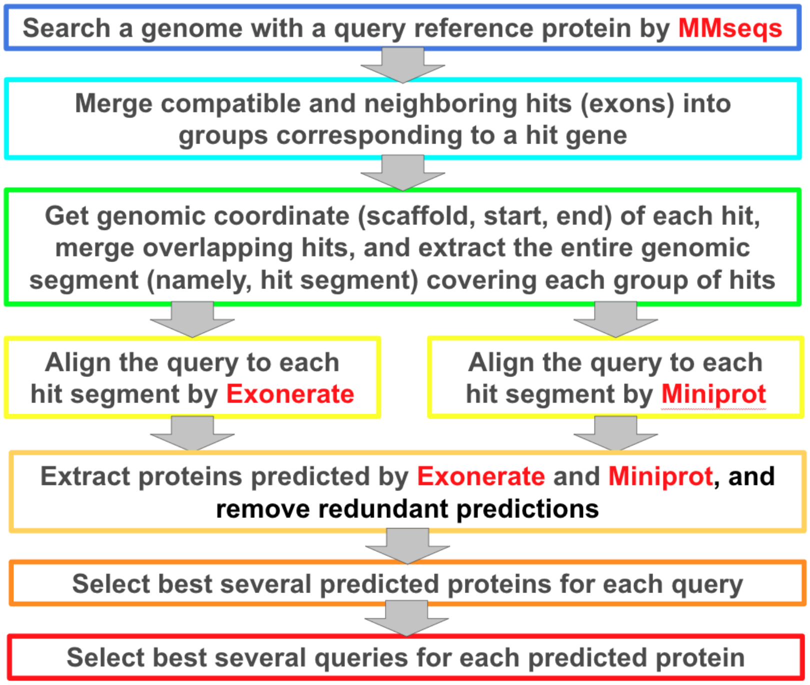


**Figure S21. A pipeline to assemble sequences of candidate orthologs from draft genomes without annotated proteins.**

#### M4.4 Stage4: assembling orthologs from genomic and transcriptomic reads

We gathered the proteomes assembled in the previous stages (**M4.2** and **M4.3**). We revised these proteomes using the same strategy as described in M4.3. Briefly, for each human query protein, we selected its best hit showing the highest identity within each taxonomic rank. We generated revised proteomes by replacing proteins with such best hits if the best hits show considerably higher similarity to the human orthologs. These revised proteomes were used as reference proteomes to assemble sequences from genomic or transcriptomic reads. For each genomic or transcriptomic dataset (target), we found its closely related reference proteome based on taxonomic information (see **M4.3** for details).

Because transcriptomic reads are mostly not expected to contain introns, we used whole protein sequences as references (reference protein set) to align to these reads and assemble transcript sequences. In contrast, genomic reads are expected to contain introns, and using the whole protein sequences containing multiple exons might cause the alignment to incorrectly extend to introns. Therefore, to search against the genomic reads, we first split the reference proteins into exons, taking advantage of the fact that the splicing sites are usually conserved among closely related species (*37*, *38*). We used these exon sequences (reference exon set) to align to genomic reads of a target species and assemble exon sequences, which were then concatenated to get assembled protein sequences from a target species. The strategies we used to assemble sequences from a target species is summarized in **Figure S22**.

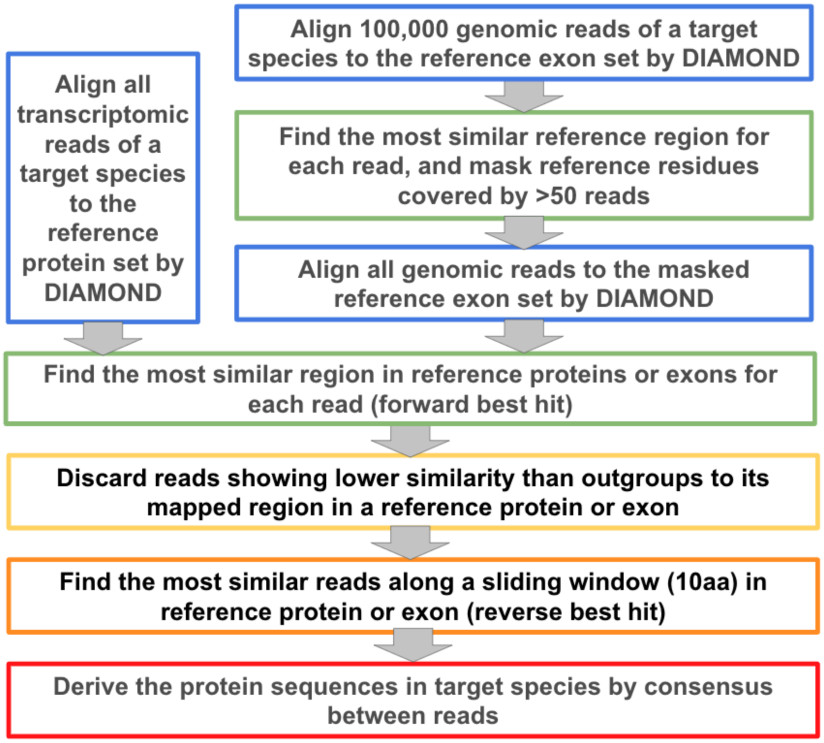


**Figure S22. The strategy to assemble orthologous sequences from transcriptomic and genomic reads using closely related reference exon and protein sets**

#### Finding the most similar reference protein/exon for each sequencing read

We aligned the genomic reads of each target species to the reference exon set; we aligned the transcriptome reads of each target species to the reference protein set. We used DIAMOND (*39*), a fast program optimized for handling large amounts of reads, for this task. We made each reference set into a database using the following command: $ diamond makedb -- in [reference_basename].fasta -d [reference_basename]. We then aligned the reads to the references using commands like: $ diamond blastx --query [reads_in_fasta_format] --db [reference_basename].dmnd -l 1 -k 0 --comp-based-stats 1 --masking 0 --out [output_prefix].tab -f 6 qseqid sseqid evalue pident qlen qstart qend slen sstart send qseq_gapped sseq_gapped --evalue 0.01 --unal 0 -p 28.

We processed the DIAMOND results to keep the alignments between a read and its most closely related queries. If a read was aligned to multiple queries, we compared each pair of read-query alignments and removed the alignment (alnA) showing remarkably worse quality than the other alignment (alnB) according to either of the following two criteria based on e-value and sequence identity: 1) $evalue_{alnA} > evalue_{alnB}^{1.2}$ and $identity_{alnA} < identity_{alnB}$ or 2) $evalue_{alnA} > evalue_{alnB}$ and $identity_{alnA} < identity_{alnB} - 10$. This process is equivalent to the "forward best hit" criterion used in ortholog detection. In the later stage, we only kept the reads showing the highest similarity to each query, equivalent to the "reverse best hit" criterion used in ortholog detection. They together constitute the "reciprocal best hit" strategy and ensure that only the reads from the orthologous protein of the reference exon/protein will be used to assemble the exon/protein sequences in each target species.

The large volume (about 100T at this stage) of our datasets created additional difficulties. Sometimes a reference protein or exon might map to repeats in the genome of a target species, and such reference protein/exon will be aligned to millions of reads, generating outputs that challenge both our file storage system and our speed to process the DIAMOND outputs. To solve this problem, we first took a sample of 100,000 reads from each genomic dataset, respectively, and we aligned this sample to the reference and processed the outputs as described above. Based on the results, we computed the sequencing depth at each position (total number of aligned reads covering that position), and we masked positions with depth above 50 by replacing the amino acid at these positions with “X”. We then used the entire set of genomic reads to align to the masked reference sequences and processed the outputs as described above.

#### Filtering aligned reads based on statistics derived from outgroups

In rare cases where the orthologs for a **reference** protein or exon were lost in evolution or not covered by the sequencing reads, non-orthologous reads that cannot map to other exons might be mistakenly identified as orthologs. We, therefore, introduced another set of criteria to filter the reads in a **target** species by similarity to the reference in addition to the reciprocal best hit criterion. A generic sequence identity cutoff is not ideal because different proteins may evolve at very different rates. We derived sequence similarity cutoffs through comparisons between reference protein/exon sets to **outgroups** whose evolutionary distance to the **reference** is expected to be larger than the **target**. These outgroups were selected among the reference proteome sets.

We first selected one **representative proteome** for each taxonomic lineage (genus, family, order, class, and phylum), preferring the reference proteome covering the largest number of human residues in the alignments made by phmmer (see above) between every human proteome and each reference proteome. For each pair of **reference** and **target** species, we found the lowest taxonomic rank at which they belong to the same lineage. The **representative proteomes** belonging to any other lineage at this rank are candidate outgroups. We went up the taxonomic rank from the shared lineage between the **reference** and the **target**, and at each rank, we checked if there are candidate outgroups, until we found such candidates and used them as **outgroups**.

For each reference-target pair, we aligned the reference exon set to the protein set of each outgroup using commands like: $ mmseqs search [reference_exons] [outgroups_proteins] [output_prefix] /tmp/mmseqs/ --start-sens 3 --sens-steps 2 -s 7 -a 1 --threads 28. We found the top hit for each reference exon and computed $outScore_{ij}$, where $i$ stands for an outgroup, and $j$ stands for a position in the query (from the reference proteome). $OutScore_{ij}$ is the blosum62 score between the query residue and the hit residue at this position; if the a query position is aligned to a gap, we considered $outScore_{ij}$ to be -2. For query exons without hit, all positions received $outScore_{ij}$ of -2. If there are multiple outgroups, we gathered the $outScore_{ij}$ scores of different outgroups for each position, $OUTScore_j = median(OutScore_{ij})$ were used as the outgroup similarity score for a position, $j$.

The **outgroup similarity scores** ($OUTScore_j$) were used to filter the alignments between the reference proteome and sequence reads from a target species. For each alignment, we computed the cumulative $OUTScore_j$ over the aligned reference residues, and this cumulative score is used as a cutoff ($Cutoff_{OUT}$) to determine if this alignment should be kept. If the cumulative blosum62 score for this alignment is lower than $0.95 \cdot Cutoff_{OUT}$, we discarded this alignment. This procedure allowed us to discard low-quality alignments between the reference and a target showing sequence similarity lower than between this reference and outgroups.

#### Selecting reads with the highest similarity the reference proteins/exons

We wanted to choose reads with the highest sequence identity from a target species for each reference exon or protein, similar to the reverse best hit strategy used in ortholog detection. However, two critical adaptations are necessary for applying this strategy to unassembled reads. First, since each read frequently covers part of a reference exon or protein, we can only compare reads covering the same region. Second, highly similar reads originating from the same genomic or transcriptomic region must be merged to remove redundancy. We converted the pairwise alignments between a reference exon or protein and reads of a target species mapped to it and passed the previous sequence similarity filters into an MSA. The pairwise alignments were built by aligning the amino acid sequences, and we replaced each amino acid with its corresponding codon to derive the alignment of nucleotide sequences.

We then introduced a 30 bp sliding window moving by one codon at a time to this MSA. Within a window, we clustered all the 30 bp segments from reads into groups of similar sequences using the following procedure. We ranked reads by their identity to the query from high to low. The first read initiated a cluster. Subsequently, a new read was compared to the first sequence of each cluster and assigned to the first cluster whose first sequence had no more than one mismatch from the current read. If a new read could not be assigned to existing clusters, a new cluster was initiated with this read as the first member.

Since most of the Chordata and Arthropoda genomes are diploid, we expect one or two clusters in each window if the genome is homozygous or heterozygous in this window, respectively. More than two clusters indicate contaminating reads from paralogous genes or other specimen contaminants such as bacteria, or fungi during specimen collection or library preparation. Reads from paralogs will likely show lower sequence identity to the reference exon; contaminations from other resources will likely show much lower sequencing depth because randomly introduced gDNA contamination is not likely to be more abundant than endogenous gDNA. Therefore, for each cluster ($i$), we computed the number of reads, $N_{reads}(i)$, in this cluster and the average number of mismatches to the query exon, $C_{mismatch}(i)$. We considered a cluster to be good if (1) its $N_{reads}(i)$ was at least half of $max(N_{reads}(i))$ among all clusters; (2) its $C_{mismatch}(i)$ was no larger than $min(C_{mismatch}(i)) + 2$ among all clusters.

If the number of "good clusters" was no more than two, we marked the reads not included in the "good clusters" as bad; otherwise, we marked all reads as bad. We discarded reads that were marked as bad. We used the fraction of bad reads aligned to a reference protein to indicate whether this reference protein is problematic or difficult to assemble from reads. If such a fraction is above 25%, we discarded this reference protein and all the reads aligned to it. After all these filters, the dominant nucleotide showing the highest frequency at each position in the MSA of reference exon or protein was used to construct the coding sequence from the target species. If such a dominant nucleotide did not exist, we replaced that position with a gap. The exon sequences were further translated to amino acid sequences, and sequences of exons from the same protein were concatenated to obtain the protein sequence of each target species.

##### M4.5 Stage5: generating multiple sequence alignments for human proteins

We gathered candidate orthologs of each human protein assembled from draft genomes and sequencing reads. We aligned each query human protein to its candidate orthologs using HMMER (*36*). We used four rounds of iterative HMMER searches with e-value cutoffs (--incE option) of 1e-10, 1e-7, 1e-4, and 1e-1 and bitscore cutoffs (-T option) of $0.75L$, $0.5L$, $0.25L$, and 0, where $L$ is the length of the query human protein. Homologs found in each round of sequence search were used to construct the sequence profile for each query human protein using HMMER hmmbuild, which was used to align more homologs in the next round. HMMER also has the tendency to split the alignment between a query and a hit protein into multiple HSPs. We compared multiple HSPs and merged pairs of HSPs if they satisfied the following criteria: 1) overlap in covered query residues by both HSPs is smaller than half of their coverage in the query; 2) overlap in covered hit residues by both HSPs is smaller than half of their coverage in the hit; 3) merging the two HSPs can extend the query coverage by both HSP by at least 20 residues. If two HSPs should be merged, in their overlapping region, we chose the one displaying higher sequence similarity (measured by total blosum62 scores) between the query and the hit. This HSP merging routine could combine several HSPs into one alignment between the query and a hit, and if there were more than one merged alignment for a hit, the one showing a higher total blosum62 score was kept.

We obtained the MSA for each query protein, and selected the best hit from each source (either a draft genome or a SRA dataset, labeled by its NCBI accession). We first identified the set of sources with a clear single best hit to the human query by the following criteria: 1) this hit is aligned to >50% of query residues; 2) this hit is the best hit, showing the highest total blosum62 score, $blosum_{BEST}$, among hits from this source; 3) other hits from this source either show total blosum62 score lower than $0.8 \cdot blosum_{BEST}$ or show higher than 95% sequence identity to the best hit among positions in the MSA covered by both this hit and the best hit. Using these single best hits, we obtained a sequence profile (with frequency of each amino acid at each position) for the query protein. Using the sequence profile, we then scored the hits from sources without clear single best hits. We focused on hits aligned to at least 30 query residues and showing query coverage higher than the maximal coverage of each source, and we then selected the hit showing the highest score to the sequence profile based on the blosum62 matrix.

After the previous step, the MSA for each human protein contained the best single hit from each source. We were concerned that the sequences assembled from SRA reads might contain a large fraction of gaps due to the low sequencing depth of some species or show low sequence identity to the query because non-orthologous reads were included. Therefore, we filtered these MSAs to remove such sequences. For each source ($i$) in a MSA, we computed the query coverage ($Qcov_i$) and average blosum score over aligned positions ($Score_i$). We calculated the 10% quantile of $Qcov_i$ for hit sequences originated from draft genomes (not SRA reads) in each phylum and class, i.e., $Qqua10_{ph}$ and $Qqua10_{cl}$. Similarly, we calculated the 10% quantile of $Score_i$ for hit sequences originated from draft genomes (not SRA reads) in each phylum and class, i.e., $Squan10_{phy}$ and $Squan10_{cla}$. We discarded hits that originated from the SRA reads if they showed worse quality than the hits from draft genomes in the same class or phylum by the following criteria: 1) $Qcov_i < min(Qqua10_{ph}, Qqua10_{cl})$ or 2) $Scov_i < min(Squa10_{ph}, Squa10_{cl})$. Such filters removed 38% of sequences that originated from SRA reads.

The human proteome contains a large fraction of intrinsically disordered and poorly conserved regions. Finding and aligning the orthologous sequences for such regions is challenging, especially in species with large evolutionary distances from humans, because these regions are frequently tolerant to insertions and deletions and show compositional biases. We first identified conserved sequence domains and globular structural domains in the query human proteins based on Pfam domains annotated in the InterPro database and structural domains we detected from AF2 models of these proteins based on DPAM. These sequence and structure domains contain 6.97M residues, 66% of all 10.54M residues in the 20,296 human proteins included in our analysis. The remaining residues mostly belong to intrinsically disordered regions (IDRs).

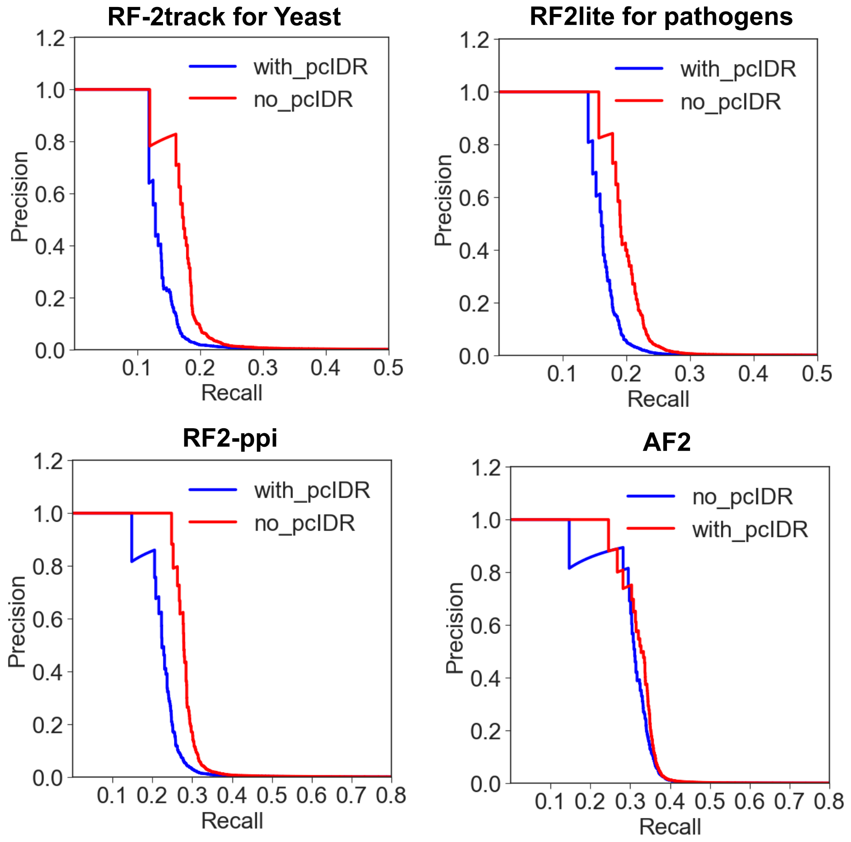


**Figure S23. Removing poorly conserved intrinsically disordered regions (pcIDR) from sequence alignments improves the performance of different tools in distinguishing true PPIs from false ones.**

We then analyzed the filtered MSAs from the previous steps to identify poorly conserved IDRs. We calculated the fraction of non-gap residues ($GapRatio_i$) and average blosum62 scores ($AveScore_i$) for each position in Chordata sequences. For each human protein with sequence or structure domains, we calculated the 25% quantile of $GapRatio_i$ and $AveScore_i$ (based on the Chordata sequences) of the residues belonging to these domains, which we denoted as $GRqua25_{dom}$ and $ASqua25_{dom}$. We then analyzed the non-domain regions using a sliding window of 10 residues: if the average $GapRatio_i$ is above $GRqua25_{dom}$ and the average $AveScore_i$ is above $ASqua25_{dom}$, we considered this window to be a relatively conserved IDR. For human proteins without sequence or structure domains, we considered a 10-residue window to be relatively conserved if its average $GapRatio_i$ and $AveScore_i$ are both above the 25% quantile of these values among the relatively conserved IDRs from proteins with domains. Neighboring relatively conserved IDR windows in sequence space were merged if there were less than 5 residues in between, and the merged segments were considered as **inter-domain conserved motifs**. The **inter-domain conserved motifs** consist of 0.80M residues (7.6%), and the remaining 2.77M residues (26.3%) were considered as poorly conserved IDRs.

We hypothesized that excluding positions from IDRs that are poorly conserved even (expected to be even less conserved in other lineages)from the MSAs of human proteins may reduce the noise in our MSAs due to poor quality of alignments in such regions, leading to superior performance in distinguishing true PPIs from false ones. However, excluding these 26.3% of human residues might remove some motifs mediating PPIs, and leading to decreased performance. Thus, we tested the performance of various AI tools in distinguishing the positive control set from the negative control set based on the full MSAs and MSAs without such poorly conserved IDRs. We found that removing poorly conserved IDRs resulted in better performance for most tools (**Figure S23**), and thus we discarded these regions from our MSAs and only used the remaining conserved or structured regions in our proteome-wide PPI screen.

#### M4.6 Evaluating the performance of our multiple sequence alignments

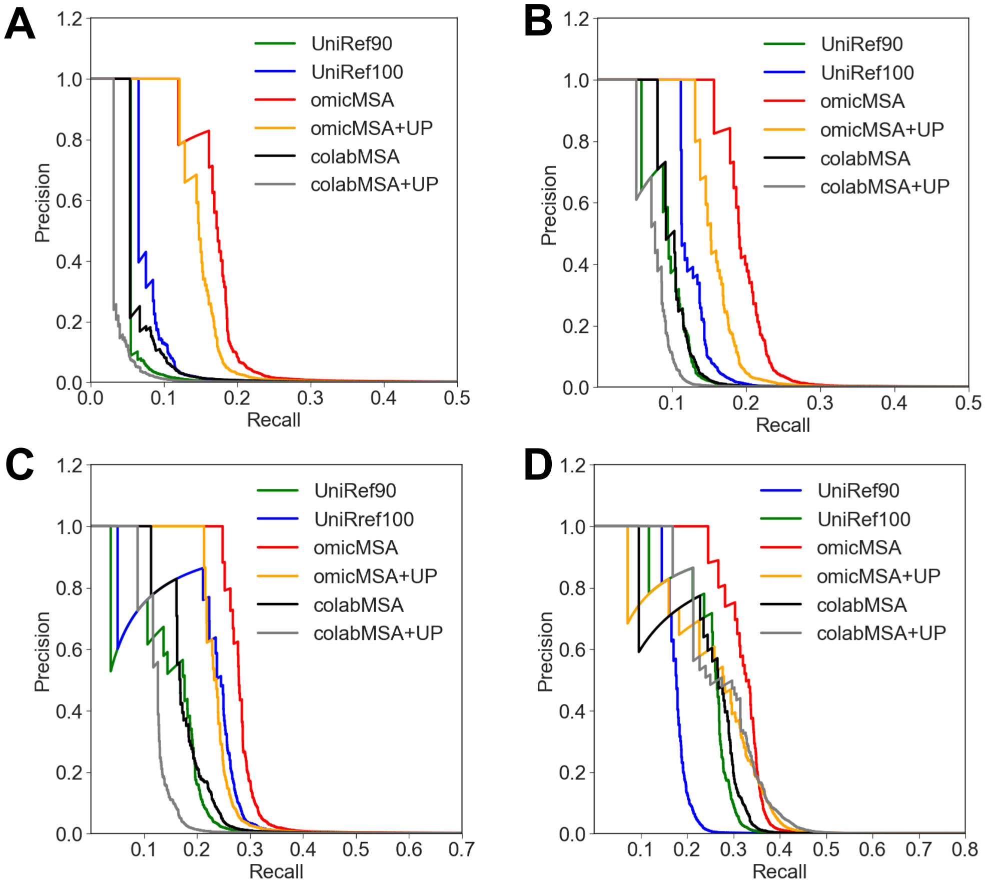


**Figure S24. Comparing the performance of our omicMSA against other common strategies for MSA construction.** We tested how these MSAs perform with different AI networks, including **(A)** RF-2track used for PPI screen in Yeast, **(B)** RF2lite used for PPI screen in pathogen, **(C)** RF2-ppi developed in this study, and **(D)** AF2. +UP means unpaired sequences are added to the pMSAs.

Our sequence alignments for human proteins were built directly from these large genomic datasets, thus we called it **omicMSAs**. In our omicMSA, we included one sequence per source data (e.g., the draft genome of a species or the genomic reads of a species) for each human protein. To build pMSAs, we concatenated the sequences from the same source. We compared omicMSAs against MSAs built by widely-used strategies (HHblits and colabfold/MMseqs) to test how they affect the performance of different tools. These widely-used strategies were used to build MSAs of single proteins, and pMSAs were obtained by concatenating the closest hits of the two query proteins in each taxon.

The common approach first is HHblits against UniRef, a widely used strategy to prepare MSA inputs for RF2 and AF2. By default, HHblits filters the output MSAs at 90% sequence identity, and we named this strategy UniRef90. However, filtering the single-protein MSAs at 90% before pairing them significantly reduces the fraction of sequences that can be paired because different sets of taxa might be kept in these two MSAs. To fix this problem, we disabled the sequence identity filter by turning on the "-all" option in HHblits and termed this alternative strategy UniRef100. Sequences from metagenomic data are frequently used to obtain deeper MSAs and improve structure prediction (*40*). Thus, we exploited the MSA building pipeline of ColabFold (*41*), which combines UniRef and metagenomic sequences. We only used sources or taxa present in both MSAs to make the pMSAs. However, the default MSA building pipeline of ColabFold also added sequences from taxa present only in one of the two proteins at the end of the pMSA, i.e., unpaired sequences. We wondered if adding unpaired sequences help to distinguish true PPIs from the false ones, and thus we added unpaired (UP) sequences in pMSAs made from omicMSAs (omicMSA + UP).

We tested the performance of pMSAs built with these different strategies in combination of different AI networks, including RF-2track, the 2-track version of RoseTTAFold we used for PPI screen in yeast, RF2-lite, a lightweight version of the full RF2 network we used for PPI screen in bacterial pathogens, RF2-ppi, the new AI network developed in this study, and AF2, which remains the best in accuracy for distinguishing true PPIs from false ones at low signal-to-noise ratios. Regardless of what AI network is used, omicMSA shows remarkably better performance than MSAs built with other strategies. Adding unpaired sequences decreased the performance of different AI networks, although people showed previously that this strategy could improve the accuracy of predicted complex structure (*42*). We reason that degradation in performance can be attributed to the fact that unpaired sequences do not provide coevolutionary signals between the two proteins for these models, and thus they are not helpful for distinguishing true PPIs among random pairs.

#### M5. Developing a hierarchical PPI prediction pipeline

##### M5.1 Preparing proteins for PPI screening

As described above, we took the 20,296 predicted 3D structures of human proteins modeled in single frames (see **M1**) and detected domains and inter-domain conserved motifs in each protein (see **M4.5**). These domains and inter-domain conserved motifs were included in our PPI screen. We referred to such domains or inter-domain motifs as **elements** for simplicity; 36 proteins did not contain any such elements and were excluded from our pipeline. Our ability to model large protein complexes is limited by the memory of available GPUs. Thus, we attempted to split larger proteins into segments with minimal inter-residue contacts and flexible relative orientations.

In order to split proteins into segments, we first calculated the cumulative length of elements in each protein. If the cumulative length is less than 750 residues, we do not attempt to split a protein into segments. Otherwise, we obtained the three parameters between pairs of elements in a protein, including, the total number of inter-element contacts (separated by at least 10 residues in sequence and inter-residue distance < $8\text{Å}$ in 3D space), i.e., $Ncontact$, the minimal length of the two elements ($Lmin$), and the average PAE of residues between elements ( $AvePAE$). We merged pairs of elements if $Ncontact/Lmin + Ncontact/100 > AvePAE/8$, indicating the two elements have a large number of residue-residue contacts or their relative orientation, reflected by $AvePAE$, is not flexible. We attempted to split long proteins into 2-4 segments and avoid splitting elements while ensuring that the cumulative length of elements in each segment remains below 750 residues.

We excluded 732 proteins that cannot be split into shorter (<750 aa) segments. We collected residues belonging to domains or conserved motifs in each segment, and these were the residues to be modeled in our PPI screens. These residues are not always continuous in sequence due to the exclusion of poorly conserved IDRs, and we filled gaps that were10 residues or less. We obtained a total of 21,127 segments, and they represent 19,528 (95%) human proteins and constitute 191 million pairs.

##### M5.2 Prioritizing pairs based on subcellular localization information

To maximize the number of true PPIs we can detect with limited computer resources, we prioritized human protein pairs based on their subcellular localization and strength of coevolution signals between them. We downloaded metadata (in JSON format) of all human proteins from UniProt and extracted the Uniprot keywords in the category of “cellular component” (CC). We considered protein pairs to be from the same cellular locality if they share any CC keywords; we considered them to be from different localities if they both have CC keywords but show no overlap; the rest contain proteins of unknown locality. We found that confident PPIs from databases (UniProt, BioGRID, and STRING) tend to be from the same subcellular localities (**Table S3**). For example, the 85.7% pairs in our positive control set (confident PPIs from 3 databases) are from the same localities.

Therefore, prioritizing pairs in the same subcellular localities significantly reduced the scale of our computation without a significant decrease in the number of true PPIs that can be detected. Among the 191M pairs we sought to screen, we focused on the 53.8M (28.2%) pairs sharing CC keywords; the majority of true PPIs to be are expected to be in this set. In addition, we screened the 57.4M pairs that include proteins without any assigned CC keywords. Such poorly annotated entries likely correspond to poorly characterized human proteins. Although these pairs do not have a very low chance to be true PPIs, they may connect poorly characterized proteins to well-characterized partners, shedding light on their functional characterization. In total, 111.2M (53.8M + 57.4M) protein pairs were selected based on this subcellular locality filter.

**Table S3. Partitioning of protein pairs in different sets by subcellular localities.**

| | Same locality | Different localities | Unknown locality |
|---|---|---|---|
| All protein pairs | 58.5M (28.4%) | 85.9M (41.7%) | 61.5M (29.8%) |
| Pairs in our screen | 53.8M (28.2%) | 79.4M (41.7%) | 57.4M (30.1%) |
| Confident PPIs (3 databases) | 3,419 (85.7%) | 209 (5.2%) | 360 (9.0%) |
| Confident PPIs (2 databases) | 18,461 (83.2%) | 1,591 (7.2%) | 2,132 (9.6%) |
| Confident PPIs (1 database) | 111,412 (67.4%) | 31,439 (19.0%) | 22,464 (13.6%) |

#### M5.3 Prioritizing pairs based on coevolution signals

We previously implemented Direct Coupling Analysis (DCA)(*43*), a statistical method to detect coevolution between positions in a MSA, using the Tensorflow framework. This implementation is 5 times faster than RF2-ppi when running on GPUs. Therefore, we used DCA to rapidly analyze pairs selected by subcellular localization. We filtered the pMSA built from omicMSA at 90% sequence identity using HHfilter (-id 90) (*18*). Filtered pMSA showed slightly better performance than the entire pMSA, increased compute efficiency for DCA, and reduced the file size. We converted the filtered pMSA into matrices made of integers (see the script below) to save space and stored thousands to tens of thousands of matrices into the same file to avoid having too many files and confronting the file system.

```
def parse_a3m(seqs): # seqs is a list of aligned sequences parsed from a fasta file.
    alphabet = np.array(list('ARNDCQEGHILKMFPSTWYV-'), dtype='|S1').view(np.uint8)
    seq_num = np.array([list(s) for s in seqs], dtype='|S1').view(np.uint8)
    for i in range(alphabet.shape[0]):
        seq_num[seq_num == alphabet[i]] = i
    seq_num[seq_num > 20] = 20
    return seq_num
```

We then applied DCA to matrices converted from pMSAs using the following script.

```
import os, sys, time
import numpy as np
```

```python
import tensorflow as tf

def tf_cov(x,w=None):
    if w is None:
        num_points = tf.cast(tf.shape(x)[0], tf.float32) - 1
        x_mean = tf.reduce_mean(x, axis=0, keep_dims=True)
        x = (x - x_mean)
    else:
        num_points = tf.reduce_sum(w) - tf.sqrt(tf.reduce_mean(w))
        x_mean = tf.reduce_sum(x * w[:, None], axis=0, keepdims=True) / num_points
        x = (x - x_mean) * tf.sqrt(w[:, None])
    return tf.matmul(tf.transpose(x), x)/num_points

myinput = sys.argv[1]
mygpu = sys.argv[2]
os.environ['CUDA_VISIBLE_DEVICES'] = mygpu
config = tf.ConfigProto(gpu_options = tf.GPUOptions(per_process_gpu_memory_fraction=0.95))
config.gpu_options.allow_growth = True

# each line contain first_ID, second_ID, first_length
fp = open(myinput, 'r')
pair2len1 = {}
for line in fp:
    words = line.split()
    pair = words[0] + '__' + words[1]
    pair2len1[pair] = int(words[2])
fp.close()

npz = np.load(myinput + '.npz')
with tf.Graph().as_default():
    x = tf.placeholder(tf.uint8,shape=(None,None),name='x')
    x_shape = tf.shape(x)
    x_nr = x_shape[0]
    x_nc = x_shape[1]
    x_ns = 21
    x_msa = tf.one_hot(x,x_ns)

    x_cutoff = tf.cast(x_nc,tf.float32) * 0.8
    x_pw = tf.tensordot(x_msa, x_msa, [[1,2], [1,2]])
    x_cut = x_pw > x_cutoff
    x_weights = 1.0/tf.reduce_sum(tf.cast(x_cut, dtype=tf.float32),-1)

    x_feat = tf.reshape(x_msa,(x_nr,x_nc*x_ns))
    x_c = tf_cov(x_feat,x_weights) + tf.eye(x_nc*x_ns)*4.5/tf.sqrt(tf.reduce_sum(x_weights))
    x_c_inv = tf.linalg.inv(x_c)
    x_w = tf.reshape(x_c_inv,(x_nc,x_ns,x_nc,x_ns))
    x_wi = tf.sqrt(tf.reduce_sum(tf.square(x_w[:,:-1,:,:-1]),(1,3))) * (1-tf.eye(x_nc))

    with tf.Session(config=config) as sess:
        results = {}
        for pair in npz.files:
            start_time = time.time()
            msa = np.copy(npz[pair])
            try:
                wi = sess.run(x_wi,{x:msa})
                results[pair] = wi[:len1, len1:].astype(np.float16)
            except tf.errors.ResourceExhaustedError as e:
                pass
            end_time = time.time()
        np.savez_compressed(myoutput + '.npz', **results)
```

We then applied average product correction (APC) to the original DCA scores ($s_{i,j}$) using the following formula: $dca_{i,j} = s_{i,j} - (\sum_{m=1}^{m=L1} s_{i,m}) \times (\sum_{n=1}^{n=L1} s_{n,j}) / (\sum_{m=1}^{m=L1} \sum_{n=1}^{n=L1} s_{m,n})$, where $m$ and $n$ are residues in the first protein; $dca_{i,j} = s_{i,j} - (\sum_{m=1}^{m=L2} s_{i,m}) \times (\sum_{n=1}^{n=L2} s_{n,j}) / (\sum_{m=1}^{m=L2} \sum_{n=1}^{n=L2} s_{m,n})$, where $m$ and $n$ are residues in the second protein; $dca_{i,j} = s_{i,j} - (\sum_{m=1}^{m=L2} s_{i,m}) \times (\sum_{n=1}^{n=L1} s_{n,j}) / (\sum_{m=1}^{m=L2} \sum_{n=1}^{n=L1} s_{m,n})$, where $m$ is any residue in the first protein and $n$ is any residue in the second protein. We derived the DCA score for a pair of protein by $DCA_{ProtPair} = max(dca_{i,j})$, where $i$ is any residue in the first protein and $j$ is any residue in the second protein.

**Table S4. The ability for $DCA_{ProtPair}$ to prioritize candidate PPIs from different sets.**

| DCA score | Negative controls | Confident PPIs by 1 database | Confident PPIs by 2 databases | Positive controls | Positive controls (RF2-ppi > 0.9) | Positive controls (AF2 > 0.9) |
|---|---|---|---|---|---|---|
| > 0.8 | 0.002% | 1.0% | 2.2% | 4.0% | 6.3% | 8.3% |
| > 0.5 | 0.03% | 1.5% | 3.4% | 6.1% | 12.2% | 12.7% |
| > 0.3 | 1.8% | 5.8% | 8.5% | 12.5% | 28.7% | 25.8% |
| > 0.28 | 2.9% | 8.1% | 10.8% | 14.9% | 32.3% | 29.3% |
| > 0.26 | 4.3% | 11.2% | 14.2% | 17.9% | 37.2% | 33.3% |
| > 0.24 | 6.3% | 15.2% | 18.3% | 22.6% | 43.8% | 38.8% |
| > 0.22 | 8.9% | 20.2% | 23.9% | 28.4% | 51.5% | 45.5% |
| > 0.2 | 12.2% | 26.1% | 30.3% | 34.1% | 57.6% | 51.5% |
| > 0.18 | 16.3% | 32.9% | 38.3% | 41.3% | 64.1% | 58.3% |
| > 0.16 | 21.5% | 40.9% | 47.7% | 49.6% | 71.8% | 67.0% |
| > 0.14 | 27.9% | 49.7% | 57.8% | 58.5% | 79.8% | 75.3% |
| > 0.12 | 35.5% | 58.7% | 67.9% | 68.0% | 85.9% | 82.1% |
| > 0.1 | 44.3% | 68.1% | 77.5% | 77.3% | 90.4% | 87.7% |
| > 0.08 | 54.2% | 77.1% | 85.3% | 85.3% | 93.3% | 91.3% |
| > 0.06 | 65.0% | 85.0% | 91.4% | 92.5% | 98.7% | 96.9% |
| > 0.04 | 76.7% | 91.4% | 95.7% | 96.0% | 99.9% | 99.1% |
| > 0.02 | 88.2% | 96.6% | 98.4% | 99.1% | 100.0% | 100.0% |

We obtained $DCA_{ProtPair}$ for all the 111.2M pairs selected based on subcellular localizations, and we tested how $DCA_{ProtPair}$ can distinguish protein pairs in the positive control set from those in the negative control set (yellow cells in **Table S4**). A high $DCA_{ProtPair}$ cutoff can significantly enrich the positive controls over the negative controls. For example, while 4% of positive control pairs show $DCA_{ProtPair}$ above 0.8, only 0.002% of negative control pairs show such a high score. However, a high $DCA_{ProtPair}$ will exclude a sizable fraction of true interactions in the positive control set that can be detected by the more accurate and more sensitive AI networks (green cells in **Table S4**). Therefore, we selected a low cutoff of $DCA_{ProtPair} > 0.12$ to allow the majority (over 80%) of true interactions showing interaction probability above 0.9 by RF2-ppi or AF2 to pass while greatly reducing the total number of pairs for subsequent screening stages.

##### M5.4 Using RF2-ppi to identify PPIs

Our *de novo* proteome-wide screening pipeline selected 43.6M protein pairs after the subcellular localization and coevolution (DCA) filters applied previously. We were concerned that a remarkable fraction of true PPIs would be lost due to these pre-filters. In addition, we reason that utilizing the extensive prior knowledge about human PPIs should help us to obtain a more complete picture of the human interactome. Therefore, we used both the functional (genetic) interactions from the STRING database and physical interactions collected from STRING (physical), BioGRID, and UniProt to perform an experiment-guided screen, adding all the pairs extracted from these databases (see **M1**) directly to PPI prediction at this stage.

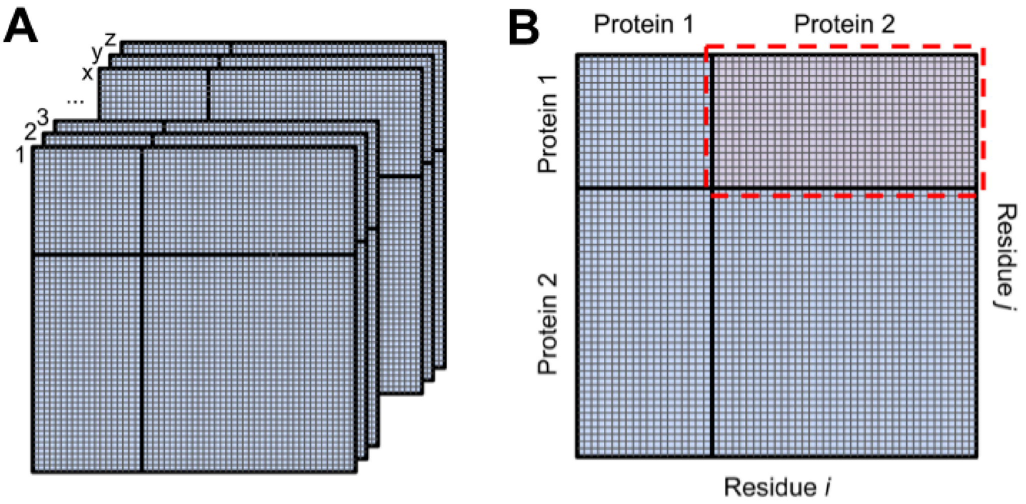


**Figure S25. Illustration of our procedure to process the RF2-ppi or AF2 outputs. (A)** The 3D matrix of dimension (L1 + L2) by (L1 + L2) by number of distance bins (37 for RF2-ppi binned by 2-20Å / 0.5Å). **(B)** The summed 2D matrix (d < 12Å) from panel A. The submatrix inside the red dashed line contains the contact probability for residues between two proteins. *Minorly adapted from the supplemental material of our previous manuscript 10.1126/science.abm4805.*

We filtered the pMSAs generated from omicMSAs at 90% sequence identity using HHfilter (-id 90), which were used as inputs to RF2-ppi. RF2-ppi uses a matrix of residue indices as its input feature, and we introduced a residue indexing gap of 200 residues between different chains/domains during training. Similarly, we provided the length of the first protein to RF2-ppi, which will in turn, introduce a 200 residue index gap in the residue index matrix. RF2-ppi produces predicted probabilities for the $C_ß$- $C_ß$ distances between all pairs of residues $i$ and $j$ which are binned between 0-20Å. Residue-residue contact probability was calculated as the sum of the probabilities for the distance bins under 12Å for each $i$, $j$ residue pair (**Figure S25A**). For any given pair of proteins, the matrix ($M_{RF2ppi}$) of contact probability is of the shape

$(L1 + L2)$ by $(L1 + L2)$, where $L1$ and $L2$ are the length of the first and second proteins, respectively. To investigate inter-protein contacts, we extracted the submatrix $Minter_{RF2ppi} = M_{RF2ppi}[:L1][L1: L1 + L2]$ shown in the red box of **Figure S25B**. The highest residue-residue contact probability in this matrix $Minter_{RF2ppi}$ was used to as the interaction probability ($RF_{ProtPair}$) for a pair of proteins.

We studied how interaction probabilities from RF2-ppi can further enrich true PPIs while filtering out random pairs. We tested various RF2-ppi interaction probability cutoffs and monitored the fraction protein pairs from different datasets that will pass each cutoff (**Table S5**). A high interaction probability cutoff can substantially enrich the positive controls over the negative controls. For example, while only 0.01% of the negative control pairs that passed our previous $DCA_{ProtProb}$ filter show $RF_{ProtPair}$ above 0.95, only 21% of the positive control pairs show such a high score. A high $RF_{ProtPair}$ will miss a small fraction of true interactions in the positive control set that can be detected by the slower and slightly more accurate AF2 (green cells in **Table S5**). Balancing the available computational resources and the recall of our screening pipeline, we chose a low cutoff of $RF_{ProtPair} > 0.3$ for our *de novo* screen, and a lower cutoff of $RF_{ProtPair} > 0.25$ for pairs with previous experimental evidence for physical or genetic interactions. These cutoffs select about 5% of pairs screened by RF2-ppi, while allowing the majority (about 80%) of true interactions showing interaction probability above 0.9 by AF2 to progress to the next stage.

**Table S5. The ability for RF2-ppi to distinguish true PPIs from false ones.**

| RF2-ppi interaction prob | Positive controls | Confident PPIs by 1 database | Confident PPIs by 2 database | Negative controls (DCA > 0.12) | Positive controls (AF2 > 0.9) |
|---|---|---|---|---|---|
| 0.95-1 | 21.17% | 6.52% | 1.14% | 0.01% | 62.58% |
| 0.9-0.95 | 22.58% | 7.14% | 1.29% | 0.04% | 65.67% |
| 0.85-0.9 | 22.99% | 7.61% | 1.44% | 0.07% | 66.64% |
| 0.8-0.85 | 23.54% | 8.03% | 1.58% | 0.11% | 67.96% |
| 0.75-0.8 | 23.90% | 8.45% | 1.73% | 0.15% | 68.58% |
| 0.7-0.75 | 24.23% | 8.76% | 1.90% | 0.22% | 69.11% |
| 0.65-0.7 | 24.64% | 9.12% | 2.09% | 0.30% | 69.81% |
| 0.6-0.65 | 25.25% | 9.55% | 2.31% | 0.42% | 70.87% |
| 0.55-0.6 | 25.94% | 10.14% | 2.59% | 0.58% | 71.58% |
| 0.5-0.55 | 26.65% | 10.80% | 2.94% | 0.79% | 72.82% |
| 0.45-0.5 | 27.23% | 11.60% | 3.43% | 1.13% | 73.26% |
| 0.4-0.45 | 28.08% | 12.63% | 4.16% | 1.65% | 74.05% |
| 0.35-0.4 | 29.60% | 14.20% | 5.28% | 2.54% | 75.55% |
| 0.3-0.35 | 31.99% | 16.54% | 7.17% | 4.17% | 77.76% |
| 0.25-0.3 | 35.68% | 20.84% | 10.68% | 7.41% | 80.14% |
| 0.2-0.25 | 43.53% | 29.14% | 17.84% | 14.42% | 83.41% |
| 0.1-0.2 | 77.34% | 64.79% | 53.56% | 49.36% | 94.97% |
| 0-0.1 | 100.00% | 100.00% | 100.00% | 100.00% | 100.00% |

#### M5.5 Using AF2, AFmm, and ColabFold for PPI modeling

We used AF2 to evaluate the probability of interaction for the ~2.2M protein pairs that passed our RF2-ppi filter (see **M5.4**). AF2, when coupled with our omicMSA, still outperforms RF2-ppi. Therefore, to resolve a more complete human proteome, we also applied AF2 to all the candidate physical interactions we collected from UniProt, BioGRID, and STRING (physical) databases, resulting in a total of ~3.1M protein pairs that were evaluated at this stage. For this large-scale screen, we only used AF2 model 3, which performed the best for complex structure prediction based on our previous benchmark (*33*).

Similar to our previous studies (*31*, *33*, *44*, *45*), we deployed AF2 to predict protein-protein complex structures using pMSAs generated from our omicMSA filtered at 90% sequence identity. We added 200 residue gap between the residue indices of the first protein and the second protein, to allow two chains to move freely in relative to each other during the modeling process. We disabled template searches in AF2 to ensure existing protein complexes in the PDB do not bias our results. In line with our RF2-ppi interaction score, we extracted the predicted distograms (probabilities for the $C_{ß}$- $C_{ß}$ distances) from AF2, and estimated the inter-residue contact probability as the sum of the probabilities for the distance bins under 12Å (**Figure S25A**). The highest inter-residue contact probability for residue pairs between the two proteins (red box in **Figure S25B**) was used to as the interaction probability ($AF2_{ProtPair}$) of this protein pair.

In addition, we also exploited the ColabFold pipeline to model complex structures using the AF2 and AFmm networks. We converted our filtered pMSA into a format compatible with ColabFold by adding two names separated by tab and stripping white space characters (this is how the ColabFold pipeline recognizes paired sequences) for each paired sequence in the pMSA. We obtained the docker image of the ColabFold pipeline and used it to run AF2 network with the following command: $ apptainer run --nv -B [absolute path to the colabfold_afweights folder]:/cache -B $(pwd):/work [absolute path to colabfold_1.5.5-cuda12.2.2.sif] colabfold_batch --model-order 3,5,1,2,4 --num-models 5 --num-recycle 3 --save-all --model-type alphafold2_ptm [folder with input msas] [folder to store outputs]. Similarly, to use the AFmm network via the ColabFold pipeline, we used the following command: apptainer run --nv -B [absolute path to colabfold_afweights]:/cache -B $(pwd):/work [absolute path to colabfold_1.5.5-cuda12.2.2.sif] colabfold_batch --model-order 3,5,1,2,4 --num-models 5 --num-recycle 3 --save-all --model-type alphafold2_multimer_v3 [folder with input msas] [folder to store outputs]. Since the fine details of the predicted complex structures are not crucial for this study, we used only three recycles (instead of 20 recycles by default) of AFmm to considerably reduce compute time. We turned on the "--save-all" flag of ColabFold to output the intermediate matrices, which include the predicted distograms that we used to derive inter-residue contact probabilities and the interaction probabilities between protein pairsas described above. However, there is a drawback of turning on this flag; that is that the volume of the output data is very large, requiring timely processing of the output data.

#### M5.6 Selecting the final set of predicted PPIs

In this study, we applied our pipeline onto four datasets: 1) protein pairs sharing subcellular localizations (keywords in the CC category), namely, the **CCcom set**; 2) pairs including proteins without subcellular localization annotations, namely, the **CCunk set**; 3) pairs showing evidence of genetic interactions according to the STRING database, namely, the **GENETIC set**; 4) pairs showing evidence of physical interactions according to established PPI databases, namely, the **PPIDB set**. For the first two datasets, we applied a pipeline consists of DCA, RF2-ppi, and AF2; for the third dataset, we applied pipeline made of RF2-ppi and AF2; for the fourth dataset, we used both RF2-ppi and AF2 to evaluate all protein pairs. We reasoned that if RF2-ppi strongly supports the interaction between a pair of proteins, the lack of support from AF2, especially since we only used a single model of AF2, should not exclude this pair from our final dataset. Therefore, we have five possible computational pipelines applied to different datasets: 1) DCA (cutoff: 0.12) → RF2-ppi (cutoff: 0.3) → AF2; 2) DCA (cutoff: 0.12) → RF2-ppi; 3) RF2-ppi (cutoff: 0.25) → AF2; 4) RF2-ppi only; and 5) AF2 only. These datasets and tailored screening pipelines for each dataset constitute a total of **9 strategies** as summarized in **Table S6.**

Next, we aimed to identify a method to estimate the accuracy of each strategy, and determine the cutoffs that maintain a high precision to obtain a final set of predictions. In our previous studies, we estimated the precision of predicted interactions in the final dataset based on the estimated precision of the benchmark set. However, such an estimate assumes that the signal-to-noise ratio in the benchmark set represents that in a proteome-wide screen. However, given the number of true PPIs in humans (roughly estimated to be between 74k and 200k (*46*)) is hard to estimate, the true signal-to-noise ratio in a *de novo* PPI screen is also hard to estimate. In addition to the *de novo* screen, we also performed knowledge-guided screens among candidate PPIs from various databases which should have substantially lower signal-to-noise, further complicating accuracy estimation.

Inspired by the widely used false discovery rate (*47*) estimation in statistics, in this study, we opt to estimate the precision by the following procedure. First, we estimate the fraction of protein pairs in the negative control set that will pass our cutoffs in the PPI screen pipeline, which we denote as $Rate_{false}$. Second, if a pipeline screened a total of $N_{input}$ pairs, and in the worst possible scenario where all these pairs do not interact, one would still expect $N_{false} = N_{input} \cdot Rate_{false}$ pairs to pass the cutoffs. Third, if the total number of PPIs predicted by a pipeline is $N_{output}$, the difference between $N_{output}$ and $N_{false}$ should correspond to the number of true predictions. Thus, we use the following equation to estimate the precision of our screen: $(N_{output} - N_{false}) / N_{output} = 1 - N_{input} \cdot Rate_{false} / N_{output}$.

The aforementioned estimate is accurate for our *de novo* screen because most protein pairs used as the inputs of this pipeline do not interact. Additionally, we screened the entire set of 140M **negative control pairs** (see **M1**) through our *de novo* screen pipeline, and thus we are able to compute an accurate estimate of $Rate_{false}$ for this pipeline. For knowledge-guided screens, the starting set contains a considerable rate of true PPIs, our method will underestimate the precision in these applications. Additionally, because we skipped the DCA step in knowledge-guided screen to maximize recall, we must estimate a $Rate_{false}$ without a

DCA filter. Therefore, we introduced two randomly selected subsets of the **negative control pairs**: one with 200,000 pairs to estimate the $Rate_{false}$ of the "RF2-ppi (cutoff: 0.25) → AF2" and "RF2-ppi only" pipelines, and another with 50,000 pairs to test the "AF2 only" pipeline.

We next searched for the cutoffs that corresponded to our desired precision of 90% to select the predicted PPIs from each strategy. As in our previous studies (*33*, *48*), we noticed that some proteins tend to have a large number of predicted PPIs and these proteins may represent a false positive hub that shows a higher predicted interaction probability with any other proteins due to some intrinsic property. For each strategy, we determined the criteria to identify and remove such false positive hubs: first, we marked a protein as a candidate false positive hub for RF2-ppi or AF2 predictions if it showed a predicted interaction probability above $Cutoff_{hub}$ with $N_{partner}$ other proteins based on RF2-ppi or AF2; second, from the predicted set, we removed pairs that include such candidate false positive hub proteins if the predicted interaction probability for this pair was below $Cutoff_{high}$, where $Cutoff_{high}$ is a higher cutoff to capture the most confident predictions regardless of candidate false positive hub proteins. $Cutoff_{hub}$, $N_{partner}$, and $Cutoff_{prob}$ are parameters that we optimized for each screening strategy to maximize the number of predicted PPIs at a 90% precision. We performed grid searches of $Cutoff_{hub}$ (0.5 to 0.95, step size 0.5), $N_{partner}$ (5 to 100, step size 1), and $Cutoff_{prob}$ (0.95 to 1, step size 0.001) to find the optimal parameters for each strategy as listed in **Table S6**.

Based on these optimized parameters, we found the interaction probability cutoff corresponding to a 90% precision for different strategies. These cutoffs were then applied to the final filter (either RF2-ppi or AF2) in each strategy. Due to the different initial signal-to-noise ratios of each dataset, the interaction probability cutoffs vary between them. For *de novo* screens, a higher cutoff is needed to exclude the dominating noise, while for knowledge-guided screens, the cutoffs for RF2-ppi or AF2 will be lower because the experimental evidence has drastically reduced the initial noise level.

**Table S6. The cutoffs to select the final predicted set aiming at 90% precision.**

| Strategy | RF2ppi cutoff | False positive hub cutoffs* |
|---|---|---|
| CCcom → DCA(0.12) → RF2-ppi | 0.994 | 0.8; 26; 0.997 |
| CCunk → DCA(0.12) → RF2-ppi | 0.999 | 0.9; 21; 1 |
| GENETIC → RF2-ppi | 0.888 | 0.8; 46; 0.967 |
| PPIDB → RF2-ppi | 0.681 | 0.7; 50; 0.967 |
| | **AF2 cutoff** | **False positive hub cutoffs*** |
| CCcom → DCA(0.12) → RF2-ppi(0.3) → AF2 | 0.941 | 0.6; 31; 0.993 |
| CCunk → DCA(0.12) → RF2-ppi(0.3) → AF2 | 0.993 | 0.75; 36; 1 |
| GENETIC → RF2-ppi(0.25) → AF2 | 0.853 | 0.9; 13; 0.986 |
| PPIDB → RF2-ppi(0.25) → AF2 | 0.693 | 0.9; 22; 0.964 |
| PPIDB → AF2 | 0.926 | |

*False positive hub cutoffs: the three values separated by semicolons represent the optimal $Cutoff_{hub}$, $N_{partner}$, and $Cutoff_{prob}$ for each strategy.

The set of predicted PPIs from all different strategies were merged, resulting in 20,853 predictions. In an attempt to obtain a confident 3D model for all predicted PPIs, the predicted PPIs supported by only RF2-ppi were subsequently modeled with 5 models of AF2 and AFmm, respectively (see **M6.2**). If none of these 10 models for a predicted PPI showed a predicted interaction probability above 0.7, we excluded this pair. A total of 2,546 pairs were excluded, which we expect to further increase the precision of our predicted set. Our final set is comprised of 18,307 predicted PPIs; among them, 1,228 pairs are in the positive control set (3,988 in total), indicating a final recall of 30.8% (1228/3988).

### M6. Analyzing predicted PPIs

#### M6.1 Integrating predicted PPIs with knowledge in databases

As described in **M1**, we obtained information about PPIs from STRING, BioGRID, and UniProt databases. Because we only processed 19,528 human proteins through our PPI screening pipeline (see **M5.1**), we filtered PPIs obtained from these external sources to only include pairs formed by these 19,528 proteins. Additionally, we identified human protein pairs that can be mapped to interacting PDB chains. We added PDB entries released up to May 2024, and repeated the procedure described in **M2.1** to identify interacting PDB chains. We compared human sequences used in our screen to sequences of PDB chains using the sensitive mode of MMseqs (-s 7). If a pair of human proteins are homologous to two interacting PDB chains, we consider that the PDB chain pair can serve as a structure template for the human protein pair.

We identify protein pairs with structural templates and split them into two categories: 1) with orthologous PDB templates if both proteins can be aligned to two interacting PDB chains with identity > 0.5 and hit (the PDB chain) coverage > 0.8 and 2) with homologous PDB templates if the pair does not have orthologous templates but both proteins can be aligned to two interacting chains with MMseqs e-value < 0.00001. A total of 3.72M pairs of proteins used in our screen found homologous (not orthologous) PDB templates, while only 9,725 found orthologous PDB templates. There can be remote templates for a PPI that cannot be detected by MMseqs. However, if the sequences have diverged sufficiently beyond the recognition of a sensitive search by MMseqs, the interaction interface between two proteins might have diverged as well, making it meaningful to ignore such templates and model the complex structure *de novo*.

We compared our final set of predicted PPIs against the PPIs from these different resources, which allowed us to estimate the fractions of predicted PPIs that are supported by other evidence (**main Figure 4A**). In addition, we found that the overlap between our predicted set and another PPI source correlates with the confidence of that set. While a large fraction (~30%) of PPIs with orthologous PDB templates or *bona fide* PPIs from the positive control set can be predicted by our *in silico* pipeline, a much smaller fraction can be predicted from other less confident PPI sources. We reason that if the ability of our computational pipeline to predict PPIs from different sets remains constant, the fraction of PPIs we identify ($f_{AI/DB}$) reflects the fraction of PPIs that are true, i.e., precision, in each database ($pre_{DB}$), which can estimated by the following formula: $pre_{DB} \cdot rec_{AI} = f_{AI/DB}$ (**main Figure 3D**). This assumption allowed us to estimate the accuracy of different PPI sources. It is worth noting that our estimates likely represent the lower limit of expected precision of these PPI databases because large databases likely include more weak interactions that are harder to detect than the positive controls.

We downloaded information about human proteins in JSON format from the UniProt website. From this file, we obtained various functional annotations of human proteins that were used in our analyses. We extracted UniProt keywords associated with each protein and grouped these keywords into the following categories: cellular components (CC), biological processes (BP), and molecular functions (MF) based on information collected from the following webpage: https://www.uniprot.org/keywords?query=*. We used overlap in these keywords to analyze if predicted PPIs tend to share common cellular components and biological pathways (**main Figure 3E; M5.2**). In addition, each PPI inherited the UniProt keywords of the two partners, allowing us to classify PPIs by these keywords and identify the enriched keywords among novel predictions. The enrichment of each keyword was evaluated using a binomial test, where p = the fraction of all predicted PPIs associated with this keyword, m = the number of novel PPIs associated with this keyword, and N = the number of novel PPIs (**main Figure 4B**).

#### M6.2 Generating and analyzing 3D models for predicted PPIs

For every predicted PPI, we exploited the ColabFold pipeline to generate 5 AF2 models and 5 AFmm models (see **M5.5**). We used these 3D models to identify the inter-protein contacts (interaction probability > 0.5 and inter-residue distance < 6Å). Residues participating in such contacts were considered as interface residues. We integrated the inter-protein contacts in 10 models (5 from AF2 and 5 from AFmm) to identify consistently predicted contacts present in $\geq 50\%$ of models. The model containing the largest number of such consistently predicted contacts was selected as the representative structure model for each predicted PPI.

We compared the structural features of interfaces for predicted PPIs and interacting PDB chain pairs that are orthologous to human proteins (see **M6.1**). Interface residues in predicted PPIs were identified as above, whereas the interface residues in PDB chain pairs were identified only by inter-residue distances (< 6Å). We classified these interface residues based on the type of regions they come from, including globular domains, flexible helices, intrinsically disordered regions, and transmembrane (TM) helices (**main Figure 3F**). Globular domains in human proteins were defined by the DPAM pipeline (*16*) in our previous work (*21*). TM helices were defined based on UniProt annotations extracted from the JSON file (see **M6.1**). Residues outside the globular domains or TM helices were further classified into flexible helices or intrinsically disordered regions based on secondary structures in the AF2 model of human monomeric proteins. The secondary structure of each residue was assigned by DSSP, and any segment of more than 5 consecutive residues annotated as “H”, “G”, or “I” (G and I represent narrower and wider helices, respectively) was considered a helix. We subsequently classified

PPIs based on the features of their interfaces: 1) we consider a PPI to be mediated by globular domains if more than 50% of interface residues in both proteins belong to such regions; 2) we consider a PPI to involve flexible helices if more than 50% of interface residues in at least one protein belong to such regions; 3) we consider a PPI to involve disordered regions if more than 50% of interface residues in at least one protein belong to such regions; and 4) we consider a PPI to be mediated by TM helices if more than >25% interface residues in at least one protein belong to such regions. These categories of PPI interfaces are not always mutually exclusive (the 4th category can overlap significantly with the 1st) and some PPI interfaces may not fall into any descrete category based on these definitions.

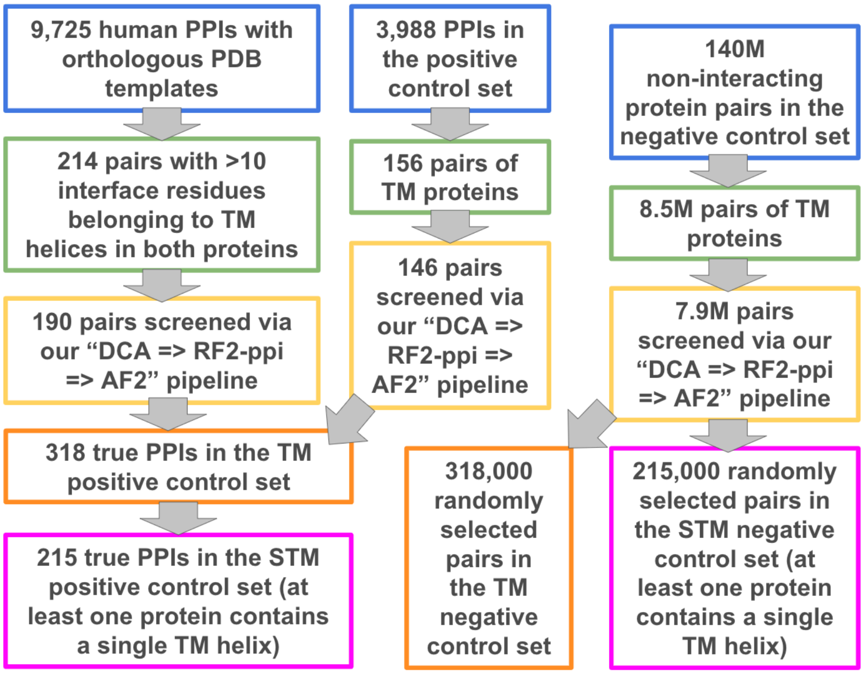


**Table S26. Our procedure to prepare additional benchmark sets to test the performance of our pipeline on transmembrane (TM) proteins.**

From the above structural feature comparisons, we found that a higher fraction of predicted PPI interfaces involves TM helices than interacting PDB chains (**main Figure 3F**). In order to study whether this is an artifact of our *in silico* PPI screen pipeline, we prepared new benchmark sets consisting of TM proteins (**Figure S26**). We assembled pairs of true interacting TM proteins (positives) from two sources. The first is the set of human protein pairs with orthologous PDB templates, from which we selected pairs with more than residues from TM helices in both proteins forming inter-protein contacts (distance < 6Å). The second is the 3,988 *bona fide* PPIs in our positive control set, from which we selected pairs of TM proteins (according to UniProt annotations). A total of 318 interacting TM proteins that went through our "DCA(0.12) → RF2-ppi (0.3) → AF2" pipeline were selected as the **TM positive control set**. From this set, we further selected pairs where at least one protein contained only a single TM helix, resulting in the **single transmembrane (STM) positive control set** consisting of 215 pairs. From the 140M negative control pairs that went through our "DCA(0.12) → RF2-ppi (0.3) → AF2" pipeline, we identified pairs of TM proteins, and randomly sampled 318,000 pairs to use as the **TM negative**

**control set.** Similarly, we created a**STM negative control set** where at least one TM protein contains only one TM helix; this set consists of 215,000 pairs.

We evaluated the performance of our pipeline based on these benchmark sets. We combined protein pairs in the **TM positive control set** and the **TM negative control set**. We ranked pairs that had not passed our DCA cutoff (0.12, stage1 pairs) at the bottom and pairs that progressed to the AF2 stage (stage3 pairs) at the top; the remaining pairs which stopped at the RF2-ppi stage (stage2 pairs) were placed in the middle. Within each stage, we ranked pairs according to the score ($DCA_{ProtPair}$, $RF_{ProtPair}$, $AF_{ProtPair}$, see **M5**) used in that stage. This way, we can evaluate the joint performance of the three stages in distinguishing positive controls from the negative controls. We computed the precision and recall at each rank and obtained the precision versus recall curve shown in **main Figure 3G**. We repeated this procedure on different benchmark sets: the **STM positive control set** versus **STM negative control set**, as well as the **positive benchmark set** versus the **negative benchmark set** (see **M3.1**), i.e., our primary benchmark sets for performance evaluation (**main Figure 3B**).

In order to understand how the 3D models of PPIs might help us understand protein function. We extracted the functional sites of each protein annotated in the UniProt JSON file. Specifically, we extracted the following features: “modified residue”, “active site”, “binding site”, “glycosylation”, and “lipidation”; we considered “modified residue”, “glycosylation”, and “lipidation” as post-translational modification (PTM) sites and the rest as functional sites. We examined if these sites overlap with interface residues both in structure models of predicted PPIs and experimental structures of PPIs (with orthologous PDB templates, **main Figure 3H**).

In order to identify disease-associated mutations on the PPI interfaces, we downloaded files containing information about single amino acid variations (SAVs) in humans and their association with diseases from UniProt at: https://ftp.uniprot.org/pub/databases/uniprot/current_release/knowledgebase/variants/humsavar.txt and https://ftp.uniprot.org/pub/databases/uniprot/current_release/knowledgebase/variants/homo_sapiens_variation.txt.gz. We compared the wildtype amino acid at each SAV position against the amino acid in our sequences to ensure these SAVs matched our protein sequences. If the fraction of mismatches in a protein was above 5%, we discarded the SAVs of that protein; as a result, SAVs associated with 246 proteins (1.3%) were discarded. In addition, we downloaded AlphaMissense predictions from https://console.cloud.google.com/storage/browser/_details/dm_alphamissense/. Similarly, we checked if the reference sequences used by AlphaMissense matched our sequences, and discarded 3 proteins (0.01%) showing >5% mismatched residues. We considered three types of evidence that support the pathogenicity of a SAV, including: 1) predicted as pathogenic by AlphaMissense; 2) annotated as “pathogenic” or “likely pathogenic” by UniProt, and 3) linked to diseases by UniProt. We included SAVs supported by at least two of the above criteria, resulting in a total of 10.4M SAVs that are potentially pathogenic. We mapped these SAVs to the interface residues of predicted and experimental protein complexes (**main Figure 3H**).

To present the protein complex models (**main Figure 4**), we extracted the residue pairs with contact probabilities greater than 0.5. For rendered complexes, we only marked the top 25 representative (one per residue) high-scoring inter-protein contacts (distance < 8Å) with a bar to connect the interacting residue pairs. If a residue at the PPI interface is associated with pathogenic SAVs, we show this residue as yellow spheres and display neighboring side chains;

however we do not show the inter-protein interaction bars in such cases for visual clarity. Further, we removed highly disordered and highly flexible regions (in the case of coiled-coils) from the rendered structure if they were not part of any interface interactions for visual clarity. These removed regions were primarily at the C and/or N terminus of proteins. Full structures containing these trimmed regions can be obtained from our website at http://prodata.swmed.edu/humanPPI/_bulk_download or https://conglab.swmed.edu/humanPPI/humanPPI_download.html. In some cases (**main Figure 5**) we overlay experimentally determined structures from the PDB onto our modeled higher order complexes to show how our predictions can add new components to known complexes. These experimental structures are shown as gray transparent surface renderings.

#### M6.3 Identifying high-order protein complexes and new components to known complexes based on our predicted PPIs

We integrated the 18,307 predicted PPIs and the 9,725 PPIs with orthologous PDB templates, two PPI sources showing high precision (**main Figure 3D**), to identify high-order protein complexes. We found that some human proteins, such as ubiquitins, are included in many PDB entries, forming contacts with other proteins although they do not function together with these proteins. We are more interested in the networks formed by predicted PPIs; however, including such interaction hubs in experimental structures might bias the results towards experimental evidence which was undesirable for our analysis. We identified such proteins that might contribute to the aforementioned problems by the following criteria: 1) less than ⅓ of the interactions derived form orthologous PDB templates are supported by AF2 interaction probabilities > 0.5, and 2) the total number of PDB-derived interacting partners with AF2 interaction probabilities ≤ 0.5 is above 5. We removed PPIs involving these proteins unless they are supported by moderate AF2 interaction probabilities (> 0.5).

After this filter, we tried several widely used methods implemented in Cytoscape (*49*) to cluster proteins using their interactions as linkages. We found most of these methods are not suitable for identifying high-order protein complexes because they assume that each protein should only belong to one cluster when in reality, a protein might serve as a component of multiple protein complexes. In addition, some multifunctional proteins can interact with many other proteins. Including such proteins in clustering might result in large clusters of functionally irrelevant proteins. We thus developed our own method to find clusters of proteins with a high density of connections (interactions) between proteins, and we require each protein in a cluster to interact with at least two other components of the same cluster (**Figure S27**).

We initially found clusters of 3 or 4 proteins where each protein is interacting with another two proteins, i.e., trimers and tetramers. We found that homologous proteins frequently interact with the same partner. While it is relatively common for paralogs, especially closely related ones, to share interacting partners (*50*, *51*), it is unlikely that these homologs are simultaneously present in a protein complex. Instead, they may represent an alternative component of a complex. Therefore, we only considered trimers and tetramers without any homology between their component proteins (detected by MMseqs with the “-s 7” flag and showing e-value < 0.00001).

As a result, we found 3,733 trimers and 2,123 tetramers, which we used as seeds for larger clusters.

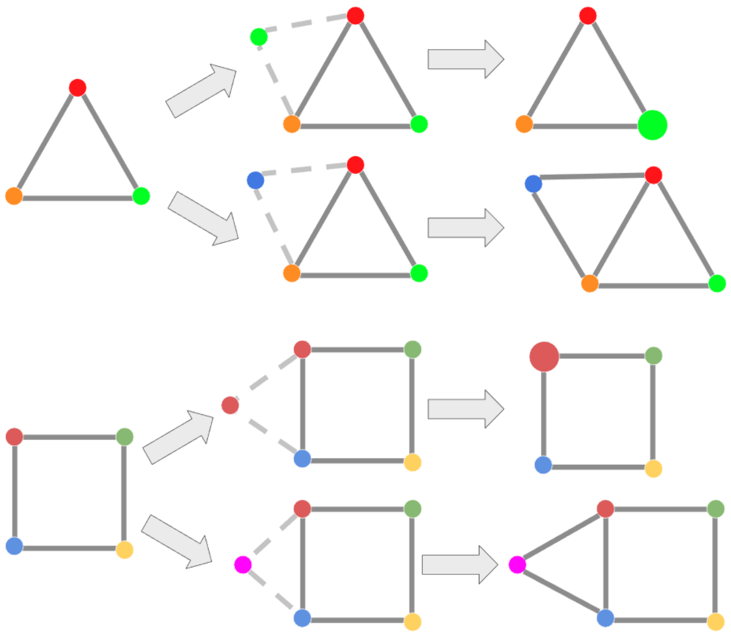

**Figure S27. Illustration of our procedure to identify putative clusters of proteins.** Each cluster is represented as a graph; proteins are represented as nodes in a graph; edges in the graph represent an interaction between two proteins. Homologous proteins are coded in the same color. Dashed lines representing the interactions between by a protein to be included in a cluster and existing proteins in a cluster. If a new protein is homologous to another protein in the cluster, it will be merged to the same node as its homolog. If a new protein is not homologous to any existing protein, it will be used as a new node in the graph.

We converted each seed into a graph, where each node in the graph can store one protein or multiple homologous proteins. We added additional proteins to each cluster, starting with the protein showing the highest cumulative AF2 interaction probability with existing nodes in a cluster: if a node contains multiple homologous proteins, the highest AF2 interaction probability among these proteins is taken as the value for this node. We only add a protein to a cluster if it interacts with at least two nodes in the cluster. If a new protein is homologous to another protein in the cluster, it will be merged to the same node as its homolog. If a new protein is not homologous to any existing proteins, it will be used as a new node in the graph. After adding all proteins interacting with at least two nodes in the cluster, we compared the clusters initiated by different seeds. We considered two clusters to be redundant if the shared nodes (nodes with any common proteins) between them is larger than 80% for the smaller cluster, and we removed the smaller cluster (with fewer nodes) in such cases. As a result, 379 clusters were identified, which can be downloaded at https://prodata.swmed.edu/humanPPI/bulk_download or https://conglab.swmed.edu/humanPPI/humanPPI_download.html. We manually studied these clusters with the help of Cytoscape, focusing on cases rich in novel predictions without strong prior evidence, i.e., not considered confident by any PPI databases and do not have orthologous PDB templates (**main Figure 5**).

To identify potential novel components to well-known human protein complexes, we extracted information about human proteins in the Complex Portal database (*51*) from the UniProt JSON file. We identified a putative new component to each complex if it satisfies the following criteria: 1) the new component is predicted to interact with at least two components; 2) over half of the predicted interactions of this new component are known components of this complex; 3) interactions between known components and the new component are not supported by strong prior evidence, i.e., not considered confident by any PPI databases and do not have orthologous PDB templates. We identified 83 proteins that are potentially new components to 77 complexes in the Complex Portal database, and the full result can be downloaded at https://prodata.swmed.edu/humanPPI/bulk_download or https://conglab.swmed.edu/humanPPI/humanPPI_download.html. We manually studied these cases with the help of Cytoscape and highlight a number of representative cases (**main Figure 5**).

### Supplemental Results

#### R1. Features of PPIs that cannot be predicted by current methods

Despite our improvements to both the input MSAs and AI networks to predict PPIs, a large fraction of PPIs remain challenging to distinguish from random pairs. We observed the same phenomenon for PPI screens in bacteria and yeast: at a 90% precision, we can only achieve a recall around 30% of our positive control PPIs. We wondered what sequence and structural features separate PPIs that can be confidently predicted from those that cannot. We identified 5,749 pairs of human proteins whose complex structures have been determined experimentally (see https://prodata.swmed.edu/humanPPI/bulk_download or https://conglab.swmed.edu/humanPPI/humanPPI_download.html. In this analysis, we only used PDB complexes of the exact human proteins, and structures of human orthologs were not used. Among these pairs, we compared a set of 2,181 pairs (**the predicted set**), showing predicted interaction probability above 0.9 by AF2 against another set of 2,434 pairs (**the missed set**), showing a predicted interaction probability below 0.1 by AF2.

The predicted set shows higher pMSA depth than the missed set (**Figure S28A**), suggesting that coevolutionary signals gleaned from deep pMSAs are important for predicting true, but possibly weak, interactions. Coevolution at PPI interfaces is a result of purifying selection: mutations disrupting PPIs will reduce the fitness and require compensatory mutations to survive natural selection. To understand whether selection pressure at the PPI interfaces affects our ability to detect PPIs, we estimated the selection pressure for interface residues using the fraction of nonsynonymous single nucleotide variations (SNVs) observed in human populations. Because synonymous SNVs are mostly not expected to cause phenotypic or fitness effects in humans, the rate of synonymous SNVs at a position can serve as an indicator of mutation rate at that position. Comparing the nonsynonymous SNV rate against the synonymous SNV rate indicates the selection pressure imposed on the nonsynonymous SNVs (*52*). A higher ratio of nonsynonymous SNVs indicates a weaker purifying selection (*52*).

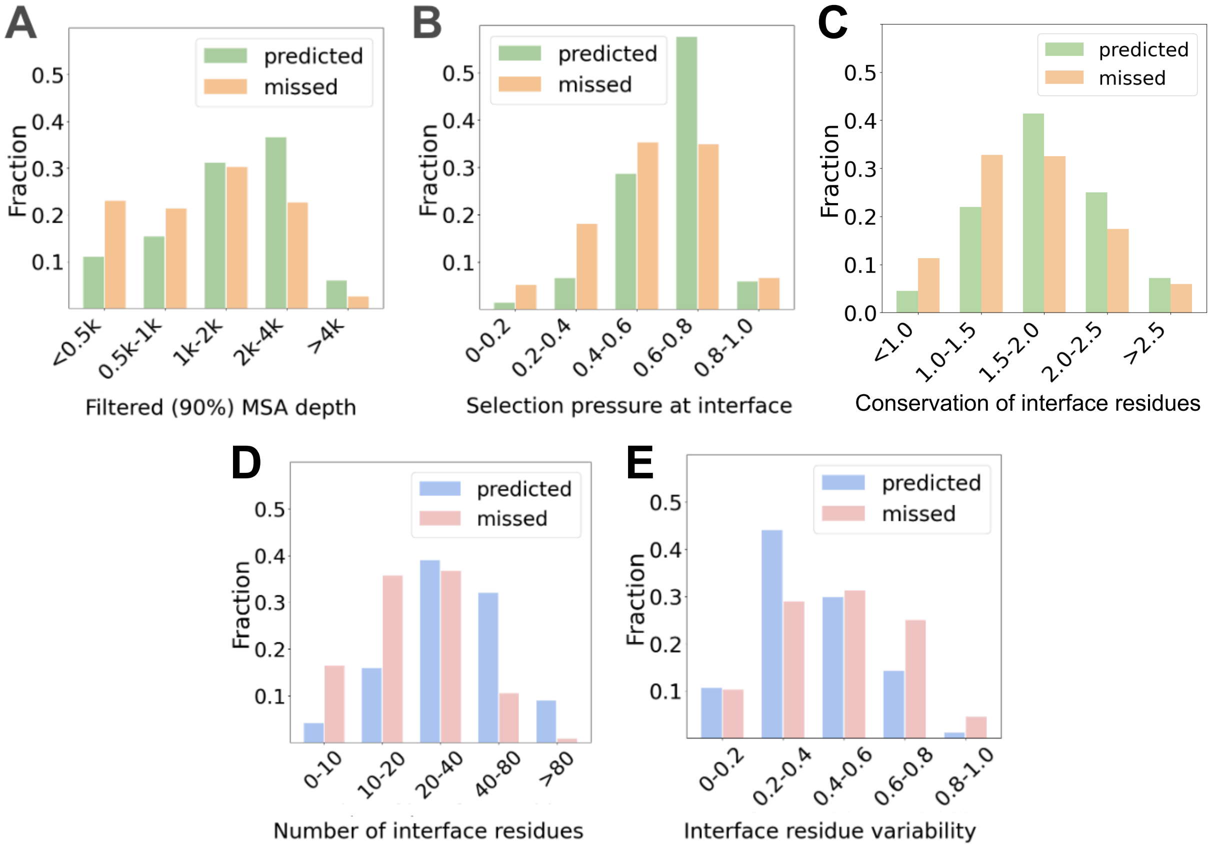


**Figure S28. Factors that affect whether a true PPI can be confidently predicted by AF2.** Predicted PPIs show the following trends compared to the PPIs that are missed: **(A)** deeper MSAs after filtering at 90% sequence identity; **(B)** moderate purifying selection pressure of interface residues; **(C)** lower conservation of interface residues estimated by entropy (X-axis, higher entropy indicates lower conservation) in the pMSA; **(D)** larger numbers of interface residues (distance to another protein < 6Å); **(E)** smaller variability in interface residues among different PDB structures, as estimated by the fraction of alternative structures where an inter-protein contact (distance < 6Å) is maintained.

We obtained information about genetic variations (in VCF format) in human populations from the gnomAD database (*53*), which contain the observed SNVs in the human genome and their frequency obtained from over 700,000 whole exome sequencing results. We mapped the SNVs in the human genome onto proteins used in our study based on genomic mapping data (UP000005640_9606_proteome.bed) provide by UniProt at https://ftp.ebi.ac.uk/pub/databases/uniprot/current_release/knowledgebase/genome_annotation_tracks/UP000005640_9606_beds. This mapping also allowed us to examine whether a SNV causes change in the protein sequence (nonsynonymous). We noticed that synonymous SVNs seem to be underrepresented in the gnomAD data, which is probably related to the fact that not all synonymous SVNs are reported by researchers who sequence human exomes/genomes because they are mostly not phenotypic. Nevertheless, such a systematic bias towards nonsynonymous SNVs is not a problem given we are only interested in the relative selection pressure (not the absolute values) between different PPI interfaces.

We identified contacts at each PPI interface as all residue pairs between the two proteins with distance less than 6Å. Residues participating in such contacts are considered as interface residues. For each PPI, we counted the number of synonymous and nonsynonymous SNVs mapped to these interface residues, i.e., $C_{syn}$ and $C_{nonsyn}$. We estimated the selection pressure happening at a PPI interface by $C_{nonsyn}/C_{syn} + C_{nonsyn}$. Interestingly, PPIs are the easiest to predict when purifying selection pressure is moderate (**Figure S28B**). We reason that when the purifying selection is too weak, it will not leave a strong trace in the sequences. In contrast, when purifying selection is too strong, residues at the PPI interface will be completely conserved, preventing the detection of coevolution. Consistent with this observation, the predicted set displays slightly lower conservation (estimated by entropy of amino acid frequencies at a position in the omicMSA, i.e $H(X) =- \sum_{i=1}^{i=n} P(x_i)log_2P(x_i)$) for residues at the PPI interface (**Figure S28C**) than the missed set.

We next analyzed the structural features of predicted and missed PPIs. As we expected, the predicted PPIs show larger interfaces (**Figure S28D**), likely related to the fact that such interfaces contain more coevolving residues and result in stronger affinity between the two proteins. Additionally, the interface between a pair of interacting proteins is not always conserved among species and some PPIs possess more than one possible interface. We hypothesize that high variability in PPI interfaces may prevent their successful modeling and detection. Focusing on 908 PPIs from the predicted set and 419 from the missed set for which there are more than 10 orthologous PDB templates. We mapped the human sequences to these PDB sequences and determined residue pairs that are in contact (inter-residue distance < 6Å). We evaluated the interface variability based on the average fraction of templates where each inter-protein contact is present. The predicted set displays smaller variability in their interfaces (**Figure S28E**), supporting our hypothesis.

### R2. Biological insights revealed from predicted PPIs

#### R2.1 PPIs of proteins involved in cancer

##### FANCM and SMC6

FANCM (Fanconi Anemia Complementation Group M) and SMC6 (Structural Maintenance of Chromosomes 6) are both proteins involved in DNA repair and genome maintenance, but they have distinct roles and functions. FANCM is a member of the Fanconi anemia (FA) protein complex, which plays a crucial role in the repair of DNA interstrand crosslinks (ICLs) (*54*). FANCM acts as a sensor and initiator of the FA pathway, recognizing and binding to ICLs, which then triggers the recruitment and activation of other FA proteins to facilitate the repair process. Multiple interaction partners were predicted for FANCM, including SMC6. SMC6 is a subunit of the SMC5/6 complex, which is involved in the maintenance of chromosome structure and integrity (*55*). The SMC5/6 complex is responsible for the resolution of DNA damage and the regulation of sister chromatid separation during cell division. FANCM is a large protein with more than 2000 amino acid residues. Our predicted model of the FANCM-SMC6 (**Figure S29**) involves the N-terminal helicase domain of FANCM, consisting of a P-loop domain (residues 71-281) and a loop region (residues 620-646) that wraps around the P-loop domain. The interface on SMC6 lies in a part of the coiled coil regions of two alpha-helices (residues 413-446 and residues 686-716). FANCM and SMC6 are both involved in the broader processes of DNA repair and genome maintenance. While FANCM and SMC6 have distinct primary functions, they are part of interconnected pathways and complexes that contribute to the overall maintenance of genome stability and the repair of DNA lesions. Experimental studies suggest that SMC5/6 complex could antagonize the potential toxic activity of yeast Mph1 helicase, the ortholog of human FANCM (*55*). Our prediction provides structural insights into the mechanism of potential suppression of FANCM helicase activity by SMC5/6 complex as the interaction interface is in the FANCM helicase domain.

##### PMS1 and FAAP24

We detect interactions between the FA complex subunit FAAP24 and PMS1. The FA complex is in the pathway of DNA repair of the Interstrand Crosslink (ICL), which are highly toxic DNA lesions that covalently link the two strands of DNA (*56*). PMS1 is in the pathway of DNA mismatch repair (MMR), which primarily deals with correcting base mismatches and small insertions/deletions that occur during DNA replication (*57*). Interactions between FAAP24 and PMS1 (**Figure S29**) provide a linkage between these two DNA repair pathways and it has been shown that MMR plays a role in sensing and initiation of ICL DNA repair (*58*, *59*).

##### ERCC8 and STK19

ERCC8, also known as CSA (Cockayne syndrome A), is in the transcription-coupled nucleotide excision repair (TC-NER) pathway (*60*). The main functions of ERCC8/CSA include recognition of RNA polymerase II stalled at DNA lesions during transcription and recruitment of other repair factors to the site of damage. Mutations in ERCC8 can lead to Cockayne syndrome, a rare genetic disorder characterized by growth failure, impaired development of the nervous system, and premature aging (*61*). Our method identifies a confident interaction between ERCC8 and STK19 (Inactive serine/Threonine Kinase 19). STK19 was recently shown to be a DNA/RNA binding protein critical in DNA damage repair and cell proliferation in the nucleotide excision repair (NER) and mismatch repair (MMR) pathways (*62*). Its predicted interaction with ERCC8 is consistent with the experimental studies. Since STK19 contains 3 packed winged helix-turn-helix domains (WH domains) and does not contain the protein kinase domain, it may not act as a protein kinase while ERCC8 contains a seven-bladed beta-propeller domain. The STK19-ERCC8 involves all three WH domains of STK19 and the fourth, fifth and sixth blades of the ERCC8 beta-propeller domain (**Figure S29**).

#### PTEN and PLEKHA1

PTEN (Phosphatase and tensin homolog) is a crucial tumor suppressor gene that plays several important roles in regulation of cell growth and division, cell cycle control, apoptosis, genomic stability, and cell migration (*63*). PTEN acts as a negative regulator of the PI3K/AKT signaling pathway that promotes cell growth and survival. It dephosphorylates PIP3 (phosphatidylinositol-3,4,5-trisphosphate), converting it to PIP2, thus counteracting the activity of PI3K. The importance of PTEN is underscored by the fact that its loss or mutation is frequently observed in various types of cancer, including breast, prostate, and brain cancers. PTEN mutations are also associated with developmental disorders and autism spectrum disorders. Understanding PTEN's functions is crucial for developing targeted therapies in cancer and other diseases where PTEN signaling is dysregulated. We predict an interaction between PTEN and PLEKHA1. PLEKHA1 (Pleckstrin homology domain-containing family A member 1, also named tandem PH domain-containing protein 1 (TAPP-1)) possesses two PH domains, which bind PIP2 and are important for targeting proteins to membrane (*64*–*66*). PLEKHA1 polymorphisms have been linked to muscular degeneration in various populations (*67*–*69*). The cellular function of PLEKHA1 has not been thoroughly studied. PLEKHA1 mainly uses the first PH domain and a beta hairpin in between the two PH domains to interact with opposite sides of PTEN, forming a large interaction interface (**Figure S29**). PLEKHA1 could help target PTEN to the membrane. By interacting with PTEN, PLEKHA1 may be able to have easy access to PTEN’s catalytic product PIP2.

#### ZNF511 and TMCO6

We predict an interaction between ZNF511 (zinc finger protein 511) and TMCO6 (Transmembrane and coiled-coil domain-containing protein 6), two poorly characterized proteins. ZNF511 has three C2H2 zinc fingers. TMCO6, despite its name, does not contain transmembrane segments according to its AF2 prediction (**Figure S29**). Instead, TMCO6 is mainly made up of alpha helical repeats. The UniProt-annotated transmembrane regions of TMCO6 (residues 338-358 and 386-406) are in fact buried hydrophobic alpha-helices in the core of the AF2 structural model. Both these two proteins have nucleus localization and are classified as prognostic markers in renal cancer with unfavorable outcome in the human protein atlas database (*70*). ZNF511 was also found to be overexpressed in prostate cancer (*71*).

#### C6orf120 and SLIT1/SLIT2

C6orf120 is a glycosylated secreted protein with an immunoglobulin-like domain. The function of C6orf120 is not known. We predict an interaction between C6orf120 and SLIT1 and SLIT2, which are two homologous long secreted proteins with multiple domains. C6orf120 interacts with its N-terminal leucine-rich repeats (**Figure S29**). Slit1 and Slit2 play critical roles in guiding axons during neural development, particularly in preventing premature midline crossing of axons (*72*). They are involved in guiding the migration of neuronal precursors during CNS development (*73*). Slit2 has been shown to inhibit growth and metastasis in some types of cancer, including fibrosarcoma and squamous cell carcinoma (*74*, *75*).

#### DSG2 and PERP

P53 is a transcriptional regulator of cell-fate decisions such as senescence, cell-cycle arrest, and apoptosis; mutations in this protein or its regulators are present in many cancers (*76*). PERP (p53 apoptosis effector related to PMP-22) is a direct downstream target of p53 with an unknown function that has been linked to apoptosis. Desmoglein-2 (DSG2) is a member of the desmosome, a family of cadherins, with mutations associated with ventricular cardiomyopathy. We predict an interaction between DSG2 and PERP (**Figure S29**) in-line with bioinformatic analyses that indicate p53 associates with DSG2. DSG2 was found to be upregulated in skin cancer (*77*) and DSG3 is over-expressed in neck, lung, and other cancers (*78*) suggesting mutations in p53 may induce overexpression of DSG2/DSG3 resulting in cancerous growth.

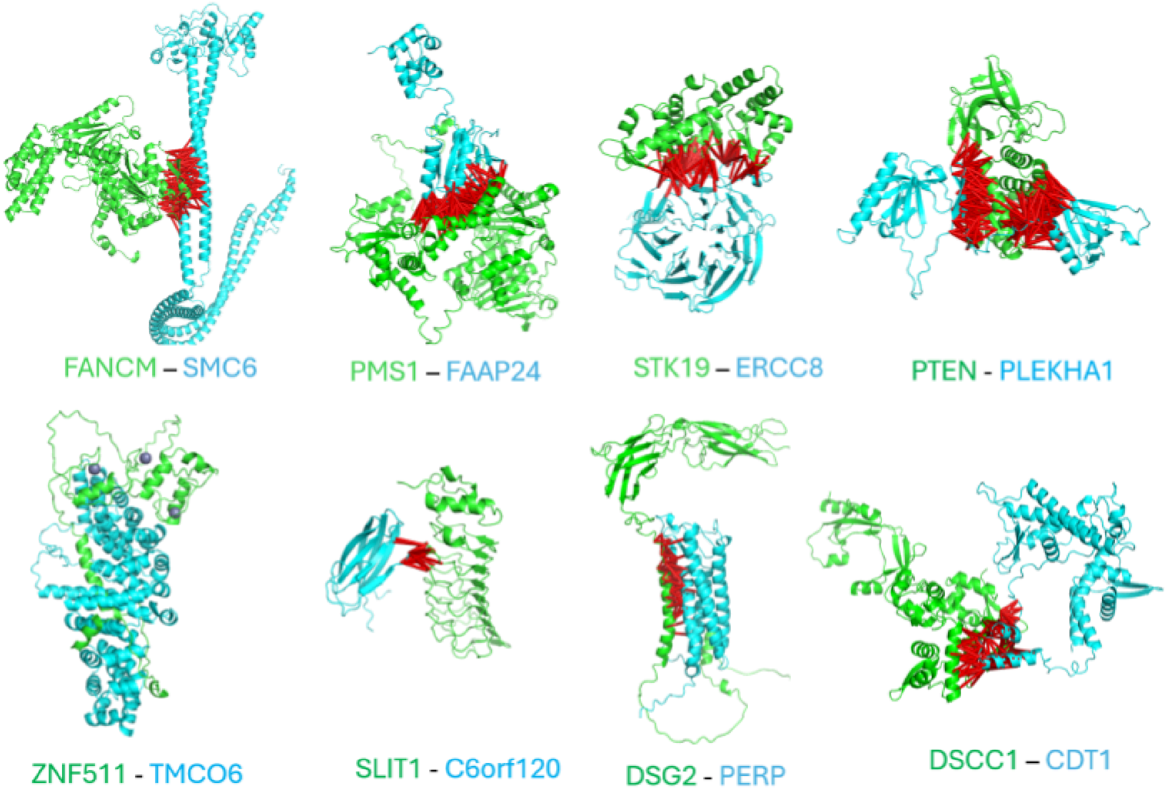


**Figure S29. Examples of PPIs of proteins involved in cancers.** The two chains are colored according to protein names displayed below each model (green / blue). Red bars connect interacting residue pairs (distance < 12Å) predicted interaction probability > 0.9 bg AF2.

#### R2.2 PPIs of G protein-coupled receptors and other membrane proteins

##### GPR143 and HPS1

GPR143, known as G-protein coupled receptor 143 or ocular albinism type 1 (OA1) receptor, is crucial for melanosome biogenesis and distribution, impacting pigmentation and vision (*79*). Mutations in GPR143 lead to Ocular Albinism Type 1, characterized by visual impairment and hypopigmentation (*80*). We predict a strong interaction between GPR143 and HPS1. HPS1 is a component of Biogenesis of lysosome-related organelles complex 3 (BLOC-3) and is crucial for the biogenesis and function of lysosome-related organelles, playing a significant role in pigmentation, blood clotting, and cellular trafficking (*80*). Mutations in HPS1 lead to Hermansky-Pudlak Syndrome Type 1, characterized by oculocutaneous albinism, bleeding tendencies, pulmonary fibrosis, and granulomatous colitis (*81*). This predicted relationship between GPR143 and HPS1 is supported by their similar disease phenotype (ocular albinism). Four mutations of GPR143 causing ocular albinism type 1 [MIM:300500] lie in the interface of GPR143-HPS1 (**Figure S30**).

##### GPR35 and GPRC5C

GPR35 and GPRC5C are both orphan GPCRs with unknown endogenous ligands (*82*, *83*), which we predict to interact via their transmembrane regions (**Figure S30**). GPR35 plays a role in the modulation of synaptic transmission and is implicated in neurogenic and inflammatory pain pathways (*84*). By interacting with the serotonin-derived ligand 5-hydroxyindoleacetic acid (5-HIAA), GPR35 promotes neutrophil recruitment to sites of inflammation, an important process in innate immune response (*85*). A recent study showed that GPRC5C promotes dormancy in hematopoietic stem cells by interacting with the extracellular ligand hyaluronic acid (*86*). The functional connection between GPR35 and GPRC5C remains to be explored.

##### GPR171 and OR56A1

Another example of a GPCR heterodimer is our predicted interaction between GPR171 and Olfactory receptor OR56A1 (**Figure S30**). GPR171 is a receptor for BigLEN, a 16-amino acid neuropeptide produced from the precursor protein proSAAS (*87*). This interaction plays a role in regulating food intake and anxiety. GPR171 is involved in pain modulation, T cell immunity regulation and TR signaling inhibition (*88*, *89*). The function of the interaction between GPR171 and OR56A1 remains to be studied experimentally.

##### HTR6 and CDH22

We also predict several other interactions involving GPCRs through their transmembrane regions. For example, HTR6 (5-hydroxytryptamine receptor 6) is a GPCR that mediates neurotransmission and plays important roles in brain functions such as neuroplasticity, memory and cognition (*90*). HTR6 was predicted to interact with CDH22 (**Figure S30**), a single-transmembrane protein in the cadherin family involved in cell adhesion in pituitary gland and brain tissues (*91*). Such an interaction provides a potential link between GPCR signaling and cell adhesion.

#### GPR152 and TMEM42

GPR152 is an understudied orphan GPCR (*92*) which we predict to interact with TMEM42, a transmembrane protein with unknown function. An HHpred search (*93*) revealed that TMEM42 is a four-TM protein homologous to a variety of transporters such as those in the family of Mg_trans_NIPA (Pfam entry: PF05653; magnesium transporter) and EamA (Pfam entry: PF00892; EamA-like transporter). These families have been classified in the DMT (drug/metabolite transporter superfamily) clan in Pfam (*94*). Its homologs have been found to contribute to bacterial multidrug resistance (*95*). TMEM42 was also predicted to form a heterodimer with its homologous protein TMEM234. Our model shows the interaction between GPR152 and TMEM42 uses a different interface than the interaction between TMEM42 and TMEM234 (**Figure S30**). The GPR152-TMEM42 interaction points to the possibility of regulation of transporter activity by a GPCR.

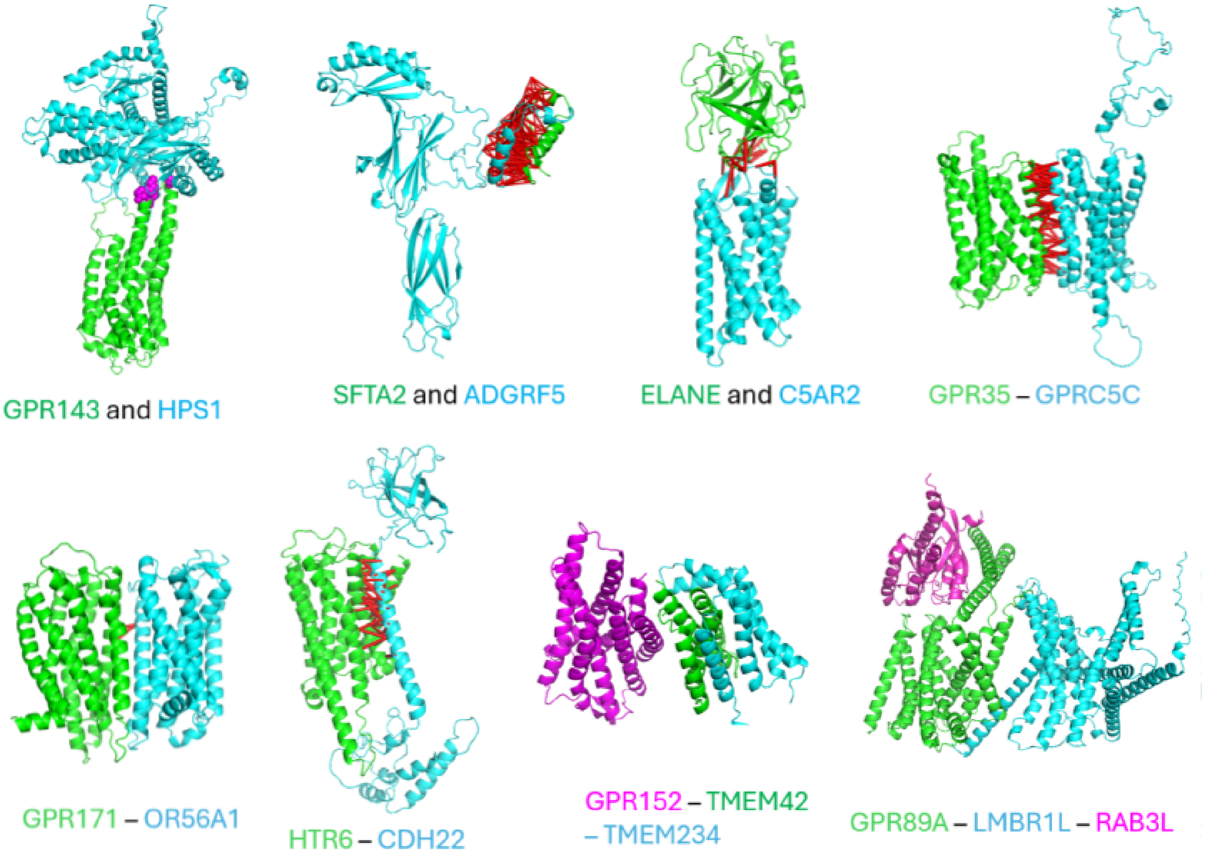


**Figure S30. Examples of PPIs of proteins involved in GPCRs and transmembrane proteins.** The two chains are colored according to protein names displayed below each model (green, blue, or purple). Purple spheres indicate interface residues associated with disease-causing SAVs. Red bars connect interacting residue pairs (distance < 12Å) predicted interaction probability > 0.9 bg AF2.

#### GPR89A and LMBR1L/RAB3L

GPR89A (Golgi PH regulator A) was annotated as a putative G protein coupled receptor. It enables voltage-gated monoatomic anion channel activity, which is crucial for the movement of ions across cell membranes (*96*). We predict that GPR89A interacts with LMBR1L, a membrane protein involved in receptor-mediated endocytosis and signal transduction. Interestingly, GPR89A and LMBR1L adopt the same fold with 9 transmembrane helices, suggesting that GPR89A is not a classic GPCR. Both GPR89A and LMBR1L have a soluble insertion with two long alpha-helices in between the fifth and sixth transmembrane helices. Their distant homologous relationship is also supported by HHpred similarity search (probability score > 0.98). GPR89A was also predicted to interact with RAB3L, a RAB family protein of small GTPase, consistent with a recent experimental study (*97*). GPR89A, RABL3 and LMBR1L are important for lymphoid development and function (*97*, *98*). GPR89A interacts with LMBR1L by using their transmembrane helices and interacts with RABL3 by using the two long helices in the soluble region (**Figure S30**).

#### R2.3 PPIs of proteins involved in immunity

##### OTUD4 and ASB8

Our method predicts strong interactions between OTUD4 (OTU Deubiquitinase 4) and ASB8 (Ankyrin Repeat and SOCS Box Containing 8, **Figure S31**), both of which are involved in the regulation of protein ubiquitination. OTUD4 is a deubiquitinating enzyme that removes ubiquitin moieties from target proteins. ASB8 is an E3 ubiquitin ligase of the SOCS box protein family, which recruits other proteins to be ubiquitinated. Both OTUD4 and ASB8 have functions related to innate immunity (*99*). OTUD4 removes K48-linked ubiquitin chains from MAVS to inhibit its degradation and thus promote innate immunity response (*100*). MAVS is an adaptor protein involved in the detection of viral RNA and downstream signal transduction by activating the protein kinase TBK1 (*100*). ASB8 induces polyubiquitination of TBK1 that targets it for degradation (*101*). The opposing roles of OTUD4 and ASB8 could help maintain a balance in the regulation of protein turnover in innate immune response.

##### FCN1 and TPSAB1/TPSAB2/HYAL1

FCN1 (Ficolin-1) is a crucial component of the innate immune system, functioning primarily in pathogen recognition and the activation of the complement system (*102*). By binding to specific carbohydrate structures on pathogens, FCN1 initiates the lectin pathway of complement activation, leading to opsonization and enhanced phagocytosis. Tryptase is a trypsin-family serine protease released by mast cells that plays a crucial role in inflammation, tissue remodeling, and allergic reactions (*103*). It contributes to the degradation of the extracellular matrix, promotion of angiogenesis, and activation of protease-activated receptors, leading to various cellular responses. Tryptase and FCN1both play important roles in the innate immune response. Our method predicted strong interactions between FCN1 and two tryptase proteins (TPSB2 and TPSAB1). Our high resolution structure model (**Figure S31**) reveals that the interaction interface covers the active site of tryptase but does not fully conceal it. The role of FCN1 on the catalytic activity of tryptase and the effect of this interaction in immune response remain to be elucidated. Interestingly, FCN1 also interacts with HYAL1 (hyaluronidase-1), a secreted enzyme involved in extracellular matrix remodeling (*104*). The interactions of FCN1 with these enzymes suggest potential roles of FCN1 in innate immune response related to the extracellular matrix. This could open new directions for research into its functions and mechanisms.

##### IFITM1 and VAMP8

We predict interactions between IFITM1 (Interferon-Induced Transmembrane Protein 1) and VAMP8 (Vesicle-Associated Membrane Protein 8). IFITM1 and VAMP8 play significant roles in cellular processes related to the immune response and vesicle trafficking (*105*, *106*). IFITM1 is primarily involved in antiviral defense (*107*). IFITM1 is induced by interferons and contributes to the cellular antiviral state by inhibiting viral entry into host cells by preventing the fusion of viral membranes with cellular membranes. VAMP8 is a SNARE protein involved in vesicle trafficking and membrane fusion (*108*) which plays roles in exocytosis, secretion of granules, and the fusion of lysosomes with phagosomes and autophagosomes. VAMP8 also plays a role in the secretion of cytokines, which are crucial for immune responses (*109*). IFITM1, being an interferon-stimulated gene, is part of the broader immune response. The interaction between IFITM1 and VAMP8 (**Figure S31**) could potentially create a negative feedback loop of interferon signaling if such an interaction negatively affects the SNARE function of VAMP8. Excessive IFITM1 proteins stimulated by interferon could interact with VAMP8 to inhibit its ability to form SNARE complexes and thus negatively affect the trafficking of interferon-bearing vesicles and interferon signaling. On the other hand, if the IFITM1-VAMP8 interaction promotes the SNARE function of VAMP8, the upregulation of IFITM1 in response to interferon signaling could enhance VAMP8's role in vesicle trafficking and cytokine secretion, creating a more robust antiviral state.

##### LOC122513141 and PLPP6

PLPP6 is a magnesium-independent polyisoprenoid diphosphatase that catalyzes the sequential dephosphorylation of presqualene, farnesyl, geranyl and geranylgeranyl diphosphates (*110*–*112*). Presqualene diphosphate is a potent inhibitor of signaling pathways contributing to polymorphonuclear neutrophils activation. PLPP6 thus play important roles in the innate immune response through the dephosphorylation of presqualene diphosphate (*110*, *113*). PLPP6 is an integral membrane protein with five transmembrane segments. Our method predicts an interaction between PLPP6 and a RING-finger containing protein (LOC122513141) with unknown function. This model reveals the interface is mediated by the C-terminal transmembrane segment of LOC122513141 (**Figure S31**). Such an interaction could mediate the degradation of PLPP6 as RING-finger proteins often act as E3 ligases.

##### KLRG1 and TLR3

KLRG1 (Killer Cell Lectin-Like Receptor Subfamily G Member 1) is a type II transmembrane protein expressed on the surface of certain immune cells, including natural killer (NK) cells and subsets of T cells (*114*). It functions as an inhibitory receptor that plays a role in regulating the immune response, particularly in modulating the activity of NK cells and effector T cells. KLRG1 binds to classical cadherins (such as E-cadherin), which helps to inhibit immune cell activation and maintain immune homeostasis. We predict strong interactions between the extracellular domains of KLRG1 and TLR3 (Toll-like receptor 3, **Figure S31**), which play critical roles in immune response by recognizing viral double-stranded RNA and inducing the production of inflammatory cytokines/chemokines and dendritic cell (DC) maturation (*115*). Such an interaction could contribute to the inhibitory role of KLRG1 by inactivating TLR3 as a pattern recognition receptor.

#### TYROBP and CD3D

TYROBP (Transmembrane Immune Signaling Adaptor TYROBP), is a signaling adaptor protein that plays a crucial role in the transmission of activation signals from various receptors on the surface of immune cells, including natural killer (NK) cells, macrophages, dendritic cells, and microglia (*116*). It is associated with several immunoreceptors such as TREM2, SIRPβ1, and KIR receptors. CD3D (CD3δ, CD3 delta) is a component of the CD3 complex associated with the T-cell receptor (TCR) on T cells. The CD3 complex is crucial for T cell activation and plays a vital role in the immune response. HHpred results suggest that TYROBP and CD3D are remote homologs as they are both single-transmembrane proteins with the cytoplasmic ITAM motif (*116*). We predict strong interactions between TYROBP and CD3D (**Figure S31**), suggesting a role of TYROBP in the function of T cell receptors.

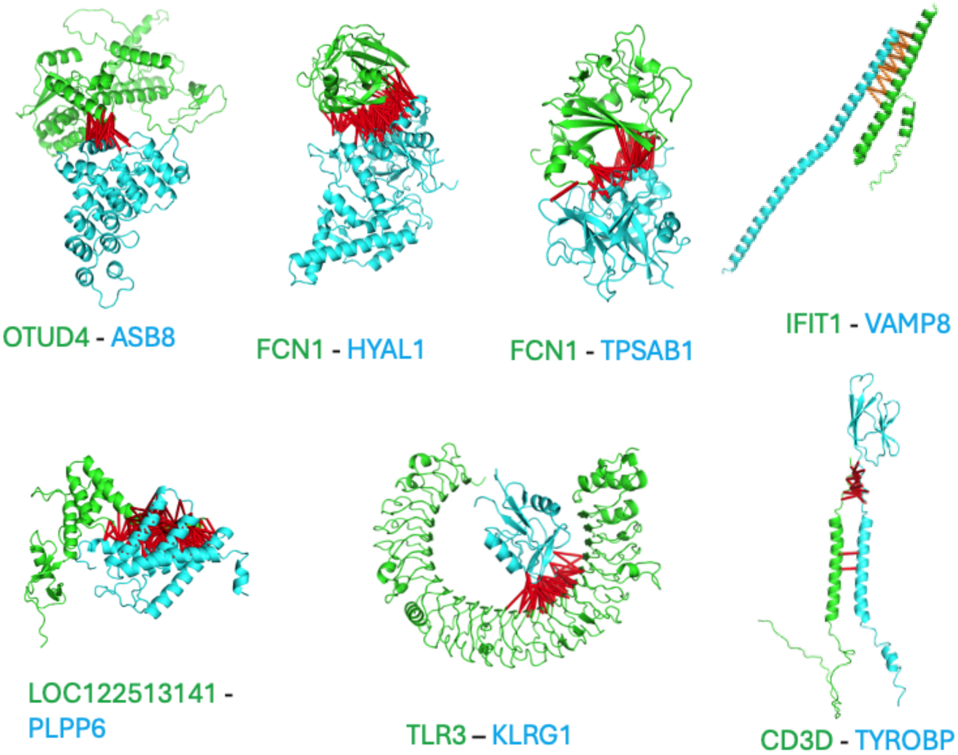


**Figure S31. Examples of PPIs of proteins involved in immunity.** The two chains are colored according to protein names displayed below each model (green, blue, or purple). Red bars connect interacting residue pairs (distance < 12Å) predicted interaction probability > 0.9 bg AF2.

#### R2.4 PPIs of mitochondrial proteins

##### COX20 and MT-CO2

COX20 is an assembly factor of mitochondrial complex IV. Our prediction revealed strong interactions between COX20 and complex IV subunit MT-CO2 (cytochrome c oxidase subunit 2) mainly through their transmembrane regions. With our high resolution structure model of the COX20-MT-CO2 interaction, we map three disease-causing mutations onto the interaction interface (**Figure S32**). The mutation M29K in MT-CO2 causes Mitochondrial complex IV deficiency (MT-C4D) [MIM:220110] (*117*), and the mutations S33L and W74C in COX20 were linked to Mitochondrial complex 4 deficiency, nuclear type 11 [MIM: 619054] (*118*, *119*). Interestingly, these three residues are spatially close and in contact with each other. These mutations likely affect the assembly process of complex IV.

##### MT-CO1 and SMIM20/CMC1/CMC2/COA4

Four complex IV (cytochrome c oxidase complex) assembly factors (SMIM20, CMC1, CMC2 and COA4) classified in the MitoCarta database (*120*) were predicted to interact with the major complex IV subunit MT-CO1 (cytochromec oxidase subunit I, **Figure S32**). These assembly factors form distinct interfaces with MT-CO1. SMIM20 and COA4 are single transmembrane proteins that mainly interact with MT-CO1 by using their transmembrane regions. CMC1 and CMC2 interact with MT-CO1 from intermembrane space. These two proteins belong to the CX9C family (*121*) characterized by two disulfide bonds formed between two α-helices.

##### CMC4 and UQCRFS1

We also predict several interactions involving mitochondrial proteins with unknown functions in the MitoCarta database (*120*). CMC4, a CX9C family protein with unknown function (*121*), was predicted to interact with UQCRFS1 (**Figure S32**), a component of mitochondrial complex III (ubiquinol-cytochrome c reductase complex), suggesting that CMC4 is a complex III assembly factor.

##### LOC128706665 and MT-CO3

A small uncharacterized protein (LOC128706665, UniProt accession: Q9HB66) has 63 amino acid residues and a single predicted transmembrane segment. This protein is annotated as “alternative protein MKKS” in UniProt as it is transcribed from a gene that overlaps with the 5’ UTR region of the MKKS (McKusick-Kaufman syndrome) gene (*122*). However, the coding region of this protein does not share any part with the coding region of the MKKS gene. This small protein was predicted to interact with mitochondrial complex IV subunit MT-CO3 by using its transmembrane segment, suggesting that it is a novel mitochondrial protein (**Figure S32**). While the coding region of this protein contains a natural variant that has been found in MKKS patients, this variant was classified as a variant of unknown significance by ClinVar and is probably not related to the MKKS disease. However, this variant was predicted to be deleterious by AlphaMissense (*123*) and is located in atransmembrane segment at the interface of the interaction model of MT-CO3 and LOC128706665. Consequently, we believe this could be a disease-causing variant by affecting the assembly of complex IV.

#### C22orf39 and ATP23

C22orf39 is a small protein with unknown function. We predict C22ord39 to interact with ATP23 (**Figure S32**), which is a thermolysin family metalloprotease (characterized by the HEXXH motif) and is an assembly factor of mitochondrial ATP synthase (*124*). Interestingly, sequence similarity searches by HHpred suggest that C22orf39 is a distant member of the CX9C family, possessing two disulfide bonds formed between two α-helices. C22orf39 could be a regulatory subunit of ATP23.

#### C8orf82 and ETFB

The electron transfer flavoprotein (ETF) enzyme complex resides in the mitochondrial matrix where it mediates the transfer of electrons from various mitochondrial flavoenzymes to the respiratory chain (*125*). This complex is made up of the ETFA (catalytic) and ETFB subunits. C8orf82, a mitochondrial protein with unknown function which we predict to form a complex with ETFB (**Figure S32**). C8orf82 could thus have functions related to the ETF complex where it may serve as an assembly factor or a regulatory subunit.

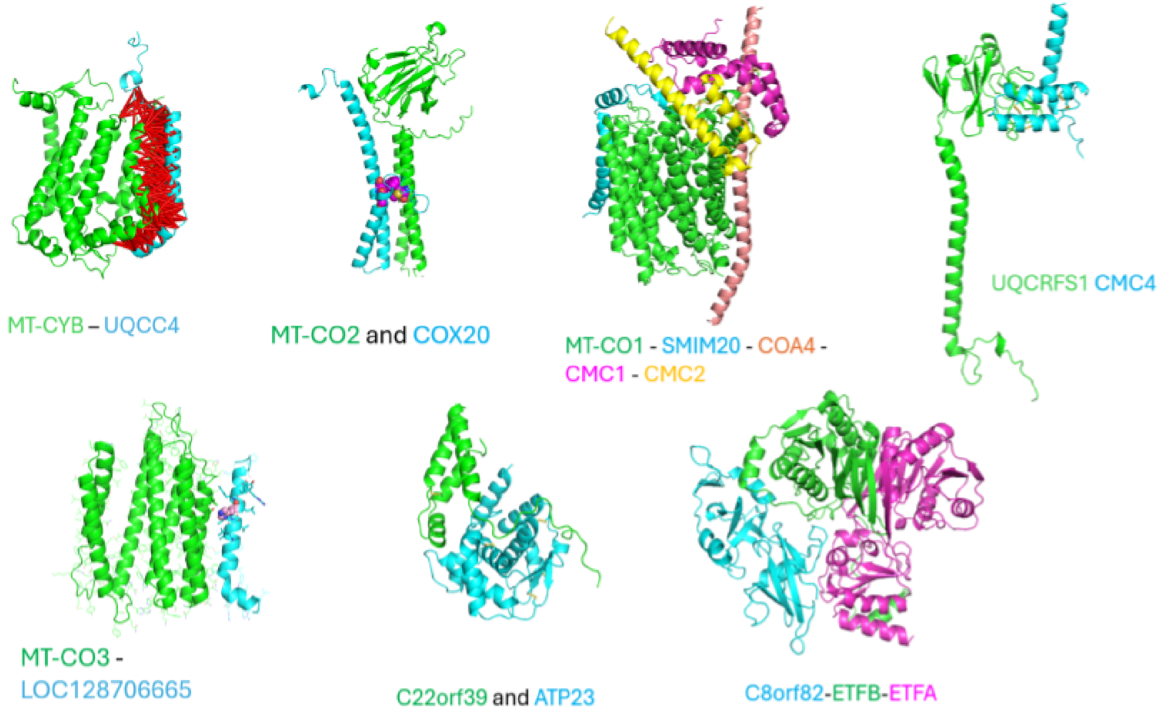


**Figure S32. Examples of PPIs of mitochondrial proteins.** The two chains are colored according to protein names displayed below each model (green, blue, orange, yellow, or purple). Purple spheres indicate interface residues associated with disease-causing SAVs. Red bars connect interacting residue pairs (distance < 12Å) predicted interaction probability > 0.9 bg AF2.

#### R2.5 PPIs of proteins involved in cilium function

##### OSCP1 and WDR54

OSCP1 was recently identified to be a novel ciliary protein, which is located at the base and axoneme of the cilium in humans and is required for cilium formation in zebrafish (*126*). WDR54 was identified as a candidate novel ciliary protein and later confirmed ciliary localization (*127*). We predict an interaction between these two proteins. WDR54 adopts the fold of a seven-bladed beta propeller, while OSCP1 is a mainly alpha helical protein. They form a large interaction interface (**Figure S33**). This interface includes a beta-hairpin in OSCP1 that extends the beta-sheet of the fifth propeller.

##### ARMC3 and CFAP70

ARMC3 (armadillo repeat-containing protein 3) is a candidate ciliary protein and that was experimentally shown to localize to cilia (*126*). Our predictions identify strong interactions between ARMC3 and CFAP70 (Cilia- and flagella-associated protein 70, **Figure S33**), a known ciliary protein that plays a role in the regulation of ciliary motility and cilium length (*128*). The functional link between ARMC3 and CFAP70 is supported by their requirements in male fertility as both their mutations caused flagellar disorganization and infertility (*129*, *130*).

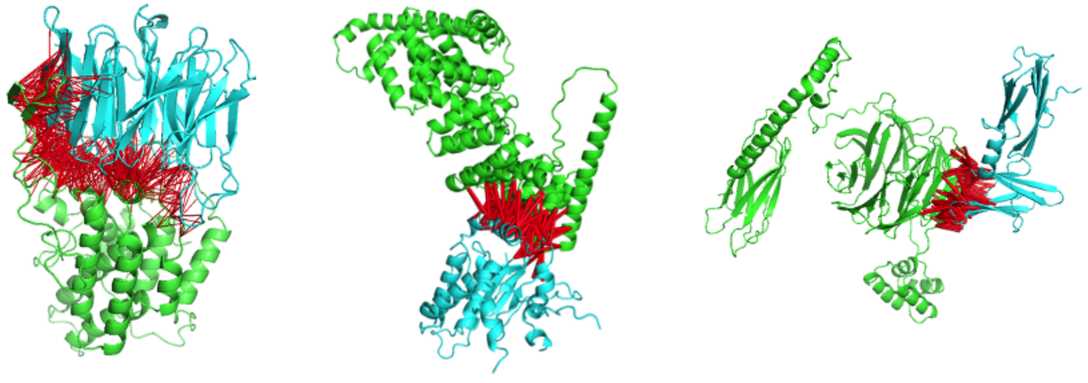


**Figure S33. Examples of PPIs of proteins involved in ciliary function.** The two chains are colored according to protein names displayed below each model (green and blue) Red bars connect interacting residue pairs (distance < 12Å) predicted interaction probability > 0.9 bg AF2.

##### BBS1 and FAIM

BBS1 (Bardet-Biedl Syndrome 1) is a core subunit of the BBSome complex, which plays a crucial role in the function of cilia (*131*, *132*). The BBSome is involved in protein trafficking to and within cilia, maintenance of ciliary structure and function, and regulation of various signaling pathways associated with cilia (*133*). Our method predicts several known interactions involving BBS1, including other subunits of the BBSome complex (BBS2, BBS4, BBS5 and BBS9), TTC8, and ARL6. Interestingly, BBS1 was also found to interact with FAIM (Fas apoptotic inhibitory molecule), a protein that has not been previously linked to ciliary functions (*134*). FAIM is composed of a duplication of two beta-sandwich domains (*135*). The interface of FAIM and

BBS1 is formed by the C-terminal beta sandwich domain of FAIM and the second and third blades of the beta-propeller domain of BBS1 (**Figure S33**). FAIM was identified as a protein that confer resistance to FAS-induced apoptosis (*136*, *137*). The strong interactions between BBS1 and FAIM suggest that FAIM could play a role in regulating the BBSome complex.

#### R2.6 PPIs with disease mutations mapped to interaction interfaces

##### C6orf226 and PEX7

C6orf226, a protein with unknown function, is predicted to have strong interactions with PEX7. PEX7 (peroxisomal biogenesis factor 7) plays a vital role in the formation and function of peroxisomes. PEX7 acts as a cytosolic receptor for peroxisomal matrix enzymes targeted to the organelle by the peroxisome targeting signal 2 (PTS2) (*138*). It is responsible for transporting several essential enzymes into peroxisomes, particularly in lipid synthesis and breakdown processes. PEX7 is also involved in the import of specific enzymes necessary for the normal assembly and function of peroxisomes. The critical nature of PEX7's function is evident from the disorders associated with its mutations. Severe mutations in PEX7 can lead to Rhizomelic chondrodysplasia punctata type 1 (RCDP1), characterized by skeletal abnormalities and intellectual disability (*138*). Some mutations in PEX7 can cause a small percentage of Refsum disease cases, affecting vision, smell, and other functions (*139*). C6orf226 is a small poorly characterized protein predicted to be mostly disordered. In the AF2 model (**Figure S34**), C6orf226 mainly adopts a loop conformation that wraps around the beta-propeller domain of PEX7, while the N- and C-termini of C6orf226 adopts small alpha-helical conformations. One likely pathogenic mutation of PEX7 according to ClinVar, H39P, lies in the interface of the predicted PEX7-C6orf226 model. This mutation is linked to Peroxisome biogenesis disorder 9B and Rhizomelic chondrodysplasia punctata type 1 (RCDP1) (*140*).

##### C12orf57 and ANKRD50

C12orf57 is a small protein linked to many disorders affecting brain function such as Temtamy syndrome (*141*), colobomatous microphthalmia (*142*), intellectual disability (*143*), and corpus callosum hypoplasia (*143*, *144*). The function of C12orf57 is unknown. We predict an interaction between C12orf57 and ANKRD50 (Ankyrin Repeat Domain 50). C12orf57 forms a helical bundle with 5 alpha-helices and interacts with the middle region of ANKR50 consisting of ankyrin repeats in our model. ANKRD50 is a key component of the SNX27–retromer–WASH super complex responsible for endocytic recycling of various membrane proteins (*145*). Retromer dysfunction has been linked to various neurodegenerative diseases (*146*). It would be interesting to test if C12orf57 also has retromer-related functions. We identify a likely disease-causing mutation at the interface of C12orf57 and ANKRD50 (**Figure S34**). This mutation (L51Q) on C12orf57 is buried inside the interface and makes several interactions with large hydrophobic residues from ANKRD50. This mutation is linked to the autosomal Temtamy syndrome characterized by intellectual disability, variable craniofacial dysmorphism, ocular coloboma, seizures, and brain abnormalities, including abnormalities of the corpus callosum and thalamus (*142*).

#### C1orf50 and AHCY

C1orf50 is a protein of unknown function conserved in Metazoa. We predict an interaction with adenosylhomocysteinase (AHCY), which was supported by large-scale experimental PPI studies in human (*147*, *148*) and fruit fly (*149*). AHCY is a hydrolase that converts S-adenosylhomocysteine to adenosine and homocysteine. AHCY mutations cause hypermethioninemia with S-adenosylhomocysteine hydrolase deficiency (HMAHCHD) [MIM:613752], a metabolic and genetic disorder. Our model (**Figure S34**) shows a disease-causing mutation (D86G) lies at the interface of the AHCY and C1orf50 interaction.

#### SCO1/SCO2 and COA6/COX17

SCO1 (Synthesis of Cytochrome c Oxidase 1) is a mitochondrial protein that plays crucial roles in the assembly of cytochrome c oxidase (COX) and copper homeostasis (*150*). It acts as a copper metallochaperone, helping to incorporate copper ions into the COX complex. SCO1 is essential for the biogenesis of the CuA site in the cytochrome c oxidase (COX) subunit Cox2. Mutations in the SCO1 gene can lead to mitochondrial disorders characterized by complex IV (cytochrome c oxidase) deficiency, resulting in conditions such as hepatic failure, encephalopathy, hypertrophic cardiomyopathy, and fatal infantile encephalopathy. We predict the interactions between SCO1 (or its paralog, SCO2) and COA6 and COX17, two other complex IV assembly factors involved in biogenesis of CuA. Several pathogenic mutations were found at the interface of SCO1-COA6 and SCO2-COX17 (**Figure S34**). Two residues involving pathogenic mutations, P174L of SCO1 and W59C of COA6, interact with each other. Both mutations cause complex IV deficiency.

#### BBS7 and WDR19

We predict strong interactions between BBS7 and WDR19. BBS7 is a subunit of the BBSome complex, and WDR19 is a subunit of the IFT-A complex. The BBSome organizes the IFT-A complex, IFT-B complex and kinesin motors into a supercomplex functioning in transportation of ciliary proteins. Our model (**Figure S34**) offers novel insights into the structural basis of the BBSome and IFT-B interactions. A pathogenic mutation, V80D in WDR19, lies in the interface of BBS7 and WDR19. This mutation is linked to Senior-Loken syndrome 8 (SLSN8) [MIM:616307], a rare autosomal recessive oculo-renal ciliopathy characterized by the association of nephronophthisis (NPHP), a chronic kidney disease, with retinal dystrophy (*151*).

#### CDAN1 and CDIN1

CDAN1 (Codanin-1) is a protein encoded by the CDAN1 gene. It has been primarily studied in the context of a genetic disorder called congenital dyserythropoietic anemia type I (CDA I) (*152*). Some key functions and roles of CDAN1 include regulation of chromatin structure and dynamics, erythropoiesis, and maintenance of the integrity and proper function of the nuclear envelope, as well as regulation of cell cycle. CDIN1 (CDAN1-interacting nuclease 1), is a protein that has been implicated in various cellular functions in cell division and growth, signal transduction, and developmental processes. Our prediction (**Figure S34**) offers structural insight into the interaction between CDAN1 and CDIN1. Several pathogenic mutations were identified in their interaction interface. Three mutations in CDAN1 (R104W, D1043V, and P1130L) are associated with anemia, congenital dyserythropoietic, 1A (CDAN1A) [MIM:224120]. Two mutations in CDIN1 (H230P and Y236C) are associated with anemia, congenital dyserythropoietic, 1B (CDAN1B) [MIM:615631].

#### TMEM218 and TMEM216

TMEM218 and TMEM216 are both transmembrane proteins associated with the primary cilia and have roles in the functioning of the ciliary transition zone, which is crucial for cilia formation and signaling (*153*). Mutations in TMEM218 can lead to ciliopathies, a group of disorders caused by ciliary dysfunction. These include conditions like Joubert syndrome and other related disorders, which are characterized by brain malformations, kidney dysfunction, and retinal degeneration. Mutations in TMEM216 are also known to cause Joubert syndrome (*154*), which is characterized by a distinctive malformation of the brainstem and cerebellum, leading to motor and cognitive impairments, and Meckel Syndrome, a severe ciliopathy that includes features such as encephalocele, polydactyly, polycystic kidneys, and liver fibrosis. We map two pathogenic mutations in position R80 (R80C and R80H) of TMEM218 (*155*) to the interface of our predicted model of TMEM218 and TMEM216 (**Figure S34**).

#### RAB28 and WDR31/BBS4

WDR31 is a recently identified new ciliary protein involved in IFT complex regulation and BBSome recruitment to the cilium (*156*). We find strongly predicted interactions between WDR31 and another ciliary protein RAB28. RAB28 is a small GTPase associated with the BBSome and intraflagellar transport (*157*). A missense variant in RAB28 G19R was predicted to be deleterious by AlphaMissense. This residue is located near the interaction interface of WDR31 and RAB28. RAB28 was also predicted to interact with the BBSome subunit BBS4. The complex structure of WDR31 and RAB28 has not been experimentally determined. Two missense variants in BBS4 (P199S and D251G) located in the interface of RAB28 and BBS4 (**Figure S34**) were predicted to be deleterious.

#### TMEM127 and LRP10

TMEM127 is an endo-lysosomal membrane protein that has been characterized as a negative regulator of mTOR signaling and a tumor suppressor (*158–160*). We predict a strong interaction between TMEM127 and LRP10 (low-density lipoprotein receptor-related protein 10). LRP10 has been found to regulate the transport and processing of amyloid precursor protein (*161*). Mutations in LRP10 have been implicated in the development of alpha-synucleinopathies such as familial Parkinson's disease and dementia with Lewy bodies (*162*). TMEM127 is a four-TM protein and LRP10 is a type I single-TM protein. Their interactions are mainly mediated by the third and fourth transmembrane segments of TMEM127, which interact with the transmembrane segment of LRP10 (**Figure S34**). The extracellular loop between the third and fourth transmembrane segments of TMEM127 also interacts with the membrane-proximal cysteine-rich domain of LRP10. A number of variants in TMEM127 were identified in the interaction interface in patients with hereditary pheochromocytoma-paraganglioma. The functional impact of the TMEM127-LRP10 interaction remains to be elucidated.

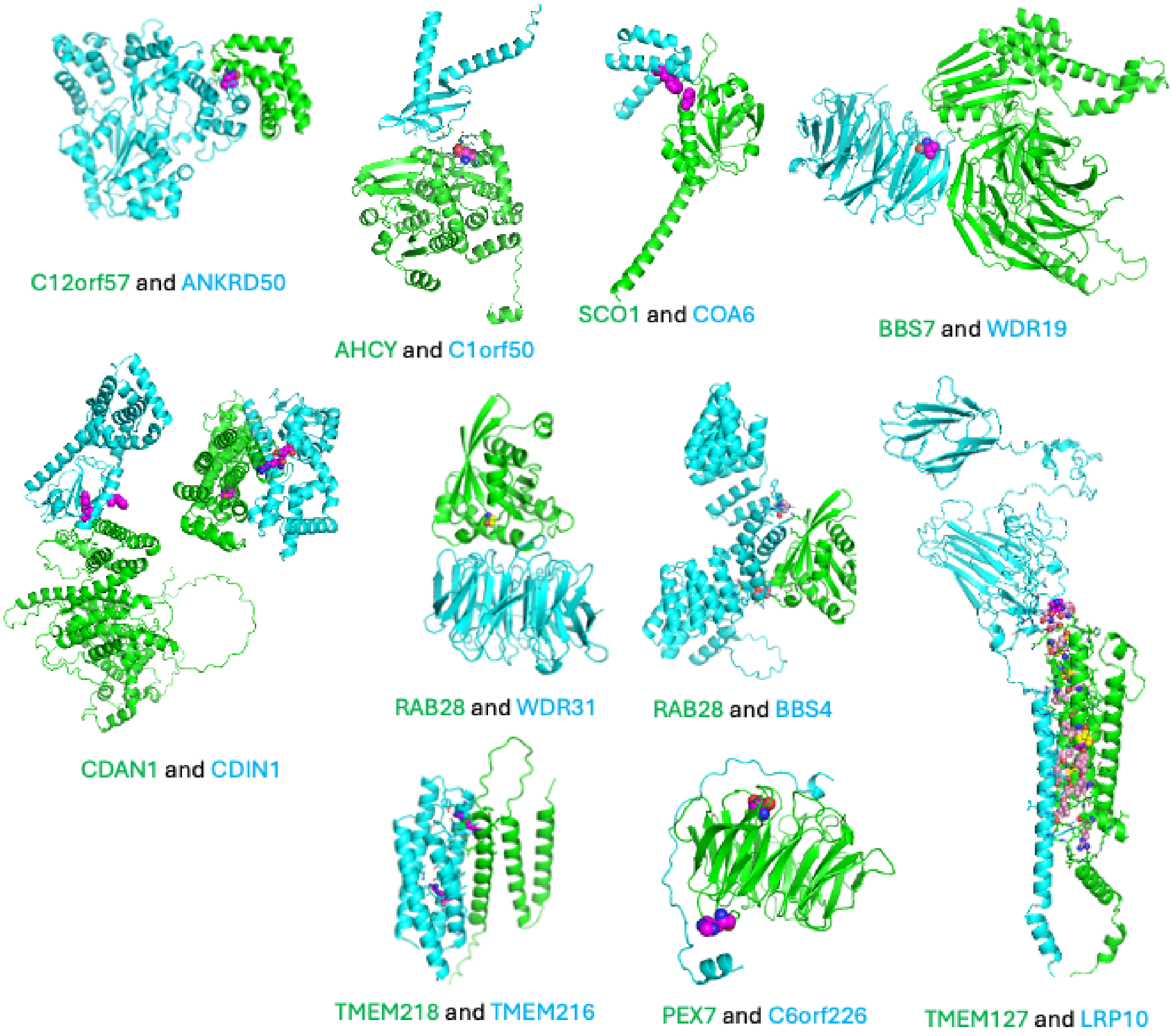


**Figure S34. Examples of PPIs with disease mutations in the interfaces.** The two chains are colored according to protein names displayed below each model (green / blue). Purple spheres indicate interface residues associated with disease-causing SAVs.

#### R2.7 Other interesting PPIs

##### YBEY and MRPS11

We predict an interaction between YbeY, an endoribonuclease which functions in rRNA maturation and ribosomal control, and 28S ribosomal protein S11 (MRPS11, **Figure S35**). In our prior bacterial interaction work (*32*) we predicted an interaction between YbeY and the 30S ribosomal protein, S11, which demonstrates how our methodology is able to detect highly conserved interactions across domains of life despite alignments only containing eukaryotic sequences.

#### SYCP2 and HORMAD1

During prophase, the axial element (AE) which is composed of meiotic cohesins and AE components, SYCP2 and SYCP3, organizes sister chromatids and provides support for meiotic recombination and homolog synapsis. HORMA domain containing proteins (HORMADs) localize along the AE and regulate meiosis. We predict an interaction between HORMA domain containing protein 1 (HORMAD1) and Synaptonemal complex protein 2 (SYCP2, **Figure S35**). Prior work shows HORMAD2 binds SYCP2 at the N-terminus (*163*) and HORMAD1 interacts with the meiotic cohesins (RAD21L and REC8) (*164*). however HORMAD1's binding to SYCP2 remained unknown.

#### LIN9 and TESMIN

The LINC/DREAM complex is an important regulator of cell cycle genes that consists of a 5-protein core. LIN9, is a component of LINC which, when depleted, leads to cell cycle arrest (*165*). Tesmin (metallothionein-like 5; Mtl5), is critical in early meiosis (*166*). We predict an interaction between an interaction between LIN9 and Mtl5 mediated by the C-terminal regions which has previously been described via truncation experiments which is essential to nuclear translocation (**Figure S35**) (*167*).

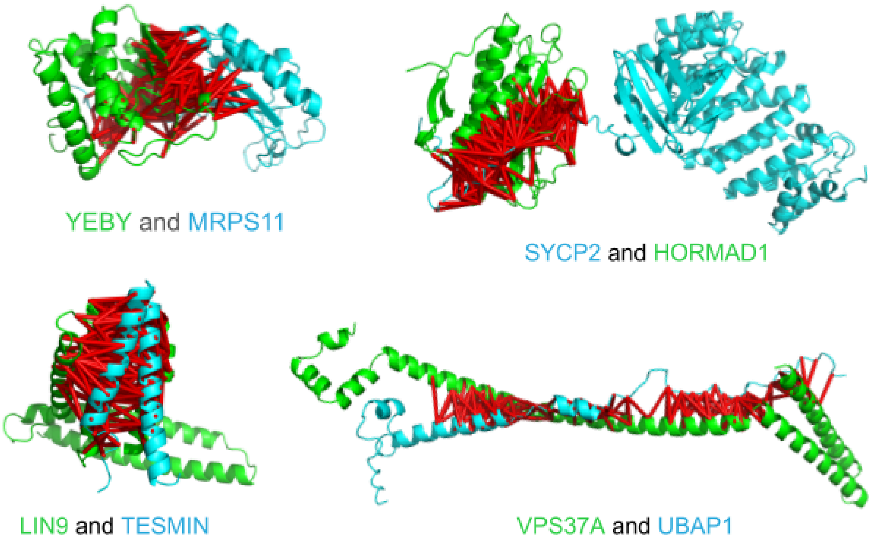


**Figure S35. Examples of PPIs with other interesting biology.** The two chains are colored according to protein names displayed below each model (green and blue) Red bars connect interacting residue pairs (distance < 12Å) predicted interaction probability > 0.9 bg AF2.

#### VPS37A and UBAP1

Endosomal sorting complexes required for transport (ESCRT), is comprised of ESCRT-0 through ESCRT-III, is highly conserved in all Eukaryotes and a number of Archaea (*168*), where it is responsible for membrane remodeling. ESCRT-1 is the bridge between ESCRT-0 and ESCRT-II where it functions to assist in creating multivesicular bodies (MVBs) by sorting ubiquitinated proteins. Ubiquitin-associated protein 1 (UBAP1) binds to ubiquitinated cargo proteins during endosomal sorting. We predict an interaction between Vacuolar protein sorting-associated protein 37A (VPS37A), a member of the ESCRT-1 complex, and UBAP1 (**Figure S35**). It has been shown that the region of VPS37A which binds to UBAP1 is part of the core core ESCRT-I-binding domain (*169*) and our model corroborates these findings.

#### R3. Multi-subunit complex modeling

The predicted binary interactions between protein pairs allowed us to identity higher-order protein complexes consisting of multiple proteins. Many of these higher-order protein complexes, although supported by previous experimental evidence, do not have experimentally determined 3D structures. The predicted structures thus provide mechanistic insights about their function, as illustrated by examples below. In these examples, components of each complex were identified based on our pairwise predictions, and the complex models were obtained in two approaches: 1) super-imposing the pairwise predictions and 2) modeling all components with the AF3 webserver at https://alphafoldserver.com/. The two approaches produced consistent results for most cases we studied, especially for the examples discussed below.

##### The NRZ complex

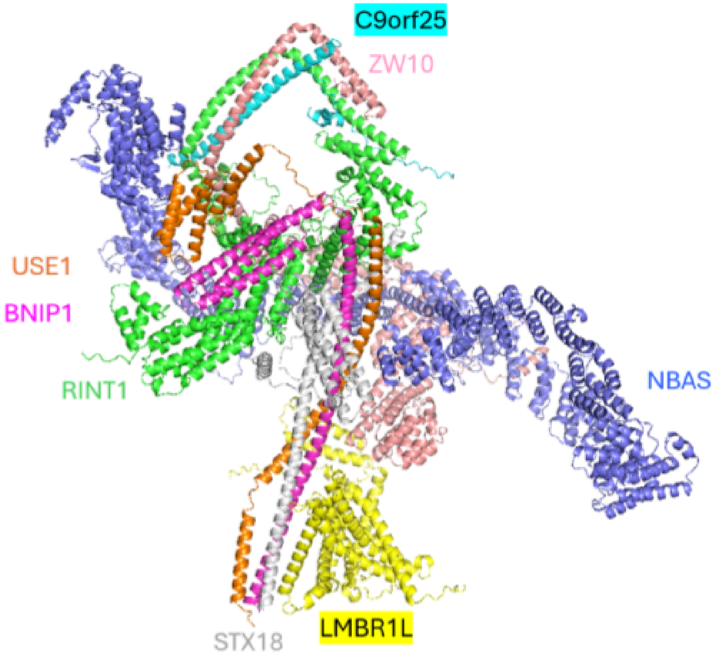


**Figure S36. A model of the NRZ complex consisting of USE1 (orange), BNIP1 (magenta), C9orf25 (cyan), ZW10 (pink), STX18 (grey), RINT1 (green), LMBR1L (yellow), and NBAS (blue).**

The NRZ tethering complex, comprising NBAS, RINT1, and ZW10, is essential for vesicle tethering to the ER and Golgi membranes, maintaining Golgi integrity, and facilitating both anterograde and retrograde transport between the ER and Golgi (*170*). Its function is crucial for proper protein sorting, cellular homeostasis, and overall vesicle trafficking within the cell. Our method predicts interactions between C19orf25 and NRZ subunits RINT1 and ZW10, suggesting that C19orf25 is a candidate subunit of the NRZ complex. C19orf25 is an uncharacterized small protein (118 residues) that has been found to bind ZW10 in tandem affinity purification screens (*171*). Multi-subunit modeling by AF3 suggests that C19orf25 forms coiled coil interactions with RINT1 and ZW10. Multiple interactions between subunits of the NRZ complex and subunits of a syntaxin complex (STX18, BNIP1 and USE1) suggest they together form a high order complex, consistent with previous studies (*170*). Interestingly, a multi-pass transmembrane protein LMBR1L interacts with both NRZ subunit RINT1 and syntaxins STX18 and BNIP1 (**Figure S36**), suggesting its roles in membrane tethering and transportation between ER and Golgi.

#### The sarcoglycan complex

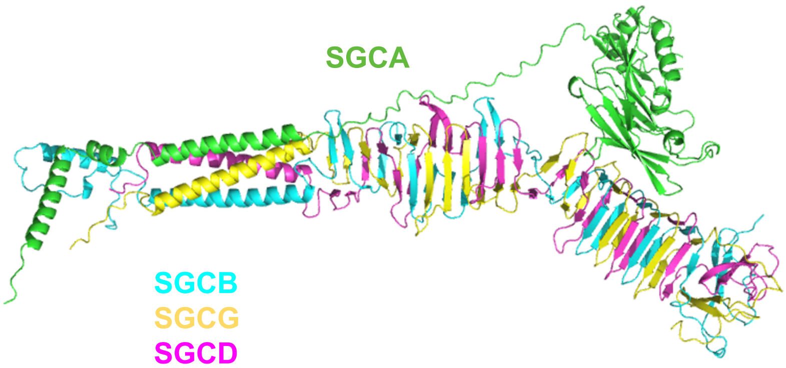


**Figure S37. A model of the Sarcoglycan complex consisting of SGCA (green), SGCB (cyan), SGCG (yellow), and SGCD (magenta).**

The sarcoglycan complex is made up of four sarcoglycan subunits (alpha, beta, gamma, and delta) that integrate into the cell membrane of skeletal muscle and cardiac muscle (*172*). In smooth muscle, epsilon- and zeta-sarcoglycans replace the alpha-and gamma-sarcoglycan in the complex (*172*). The proper assembly and function of the sarcoglycan complex are essential for maintaining the structural integrity of muscle fibers and their resistance to mechanical stress. Mutations in the genes encoding these subunits can lead to various forms of limb-girdle muscular dystrophy (LGMD) (*173*), highlighting their critical role in muscle health and function. Strong pairwise interactions were predicted between sarcoglycan complex subunits. The sarcoglycan complex with the alpha, beta, gamma, and delta subunits are modeled by AF3 (**Figure S37**). The beta, gamma, and delta subunits are homologous type II single transmembrane proteins with the extracellular C-termini consisting of repeating beta-strands. They are predicted to form a long and intertwined triple-stranded beta-helical structure resembling phage tail fiber proteins (*174*) and the spike region of the type VI secretion system spike protein VgrG (*175*). The sarcoglycan complex could originate from bacteria or bacteriophages. The alpha-sarcoglycan is a type I transmembrane protein with a long

N-terminal region in the extracellular space and a short C-terminal tail. It has two globular extracellular domains: an Ig-like domain and a ferredoxin-like SEA domain. The interactions between alpha-sarcoglycan and the triple beta-helix are mainly mediated by the Ig-like domain.

#### The SNAPc complex

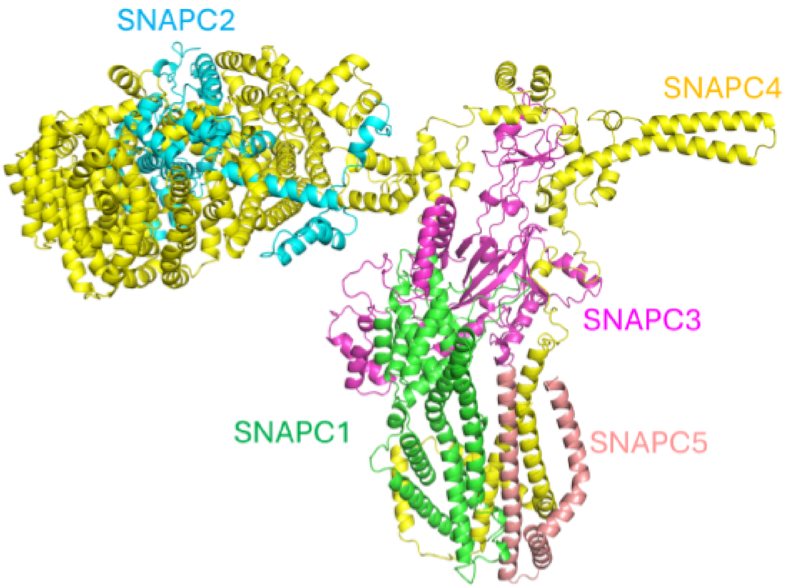


**Figure S38. A model of the SNAPc complex consisting of SNAPC1 (green), SNAPC2 (blue), SNAPC3 (magenta), SNAPC4 (yellow), and SNAPC5 (salmon).**

The snRNA-activating protein complex (also known as SNAPc or PTF) is essential for the transcription of small nuclear RNA (snRNA) genes, which are crucial components of the spliceosome machinery involved in pre-mRNA splicing (*176*). The complex specifically facilitates the transcription of snRNA genes by RNA polymerase II (Pol II) and RNA polymerase III (Pol III). The SNAPc complex is composed of five subunits (SNAPC1-SNAPC5), which together bind to the proximal sequence element (PSE) in the promoter regions of snRNA genes. Structures of parts of the SNAPc complex involving SNAPC1, SNAPC3, SNAPC5 and the N-terminal part of SNAPC4 have been characterized experimentally (*177*). However, experimental structures are not available for SNAPC2. We predict strong interactions between SNAPC2 and the C-terminal part of SNAPC4. The 3D model (**Figure S38**) shows structures of the alpha-helical domain formed by the N-terminal part of SNAPC1, the N-terminal part of SNAPC4 and SNAPC5, for which a significant portion is missing in the cryo-EM structure.

#### The TRAPPIII complex

The TRAPPIII complex is an essential multi-subunit protein complex involved in intracellular vesicle trafficking (*178*). It functions primarily as a guanine nucleotide exchange factor (GEF) for Rab1 GTPase, which is crucial for autophagy and trafficking between the endoplasmic reticulum (ER) and the Golgi apparatus. The TRAPPIII complex is composed of both core and TRAPPIII-specific subunits. The core subunits (TRAPPC1, TRAPPC2, TRAPPC2L, TRAPPC3,

TRAPPC4, TRAPPC5, TRAPPC6A, and TRAPPC6B) are shared across different TRAPP complexes. The subunits unique to the TRAPPIII complex are TRAPPC8, TRAPPC11, TRAPPC12, and TRAPPC13. While the CryoEM structure of TRAPPIII complex is available for the core subunits and parts of TRAPPC8 and TRAPPC11, the detailed structure of subcomplex formed by TRAPPC11, TRAPPC12 and TRAPPC13 are not known (*179*). We detect strong interactions between these subunits and build a high resolution predicted subcomplex model (**Figure S39**).

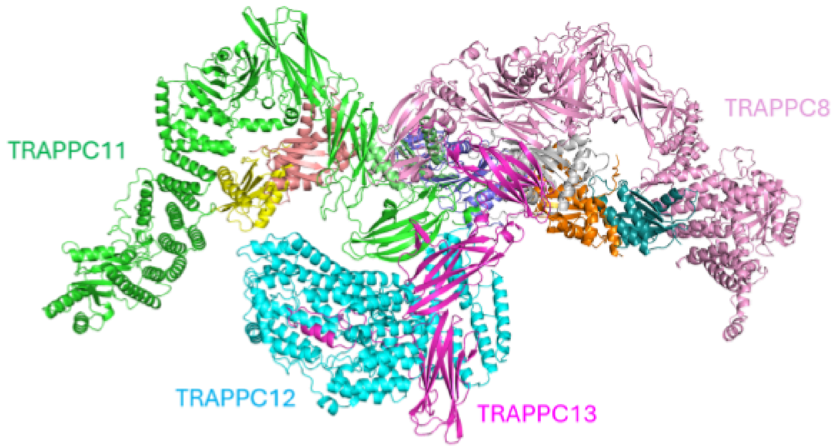


**Figure S39. A model of the TRAPPIII complex consisting of TRAPPC11 (green), TRAPPC12 (blue), TRAPPC13 (magenta), and TRAPCC8 (pink).** The core TRAPP subunits shown in yellow, salmon, grey, orange, aquamarine, and dark blue are not labeled for visual clarity.

#### The BLOC-1 complex

The BLOC-1 (biogenesis of lysosome-related organelles complex 1) complex contributes to the sorting, trafficking, and targeting of proteins and lipids to lysosome-related organelles (LROs) such as melanosomes (involved in pigment production and storage in melanocytes) and platelet-dense granules (important for blood clotting and lytic granules) (*180*, *181*). Disruptions in BLOC-1 components can lead to defects in LROs, resulting in disorders such as Hermansky-Pudlak syndrome (HPS), which is characterized by albinism, bleeding disorders, and other symptoms due to impaired LRO function. The BLOC-1 complex consists of eight subunits: BLOC1S1, BLOC1S2, BLOC1S3, BLOC1S4, BLOC1S5, BLOC1S6, SNAPIN, and DTNBP1. The atomic structure of BLOC-1 complex is not available. Our method predicts numerous interactions between BLOC-1 subunits, all of which adopt mainly coiled coil structures. Some pairwise interactions involve overlapping long alpha-helices, while other interactions are restricted between the ends of alpha-helices. The structural model of the BLOC-1 complex revealed an elongated and bent overall shape consistent with electron microscopy studies (*182*). The intertwined coiled coil complex structure consists of two four-helical bundles connected by their ends. One four-helical coiled-coil bundle consists of BLOC1S1, BLOC1S3, BLOC1S4 and BLOC1S6, and the other helical bundle involves BLOC1S2, BLOC1S5, SNAPIN and DTNBP1. The alpha-helices of the complex all have the same N- to C-direction (**Figure S40**).

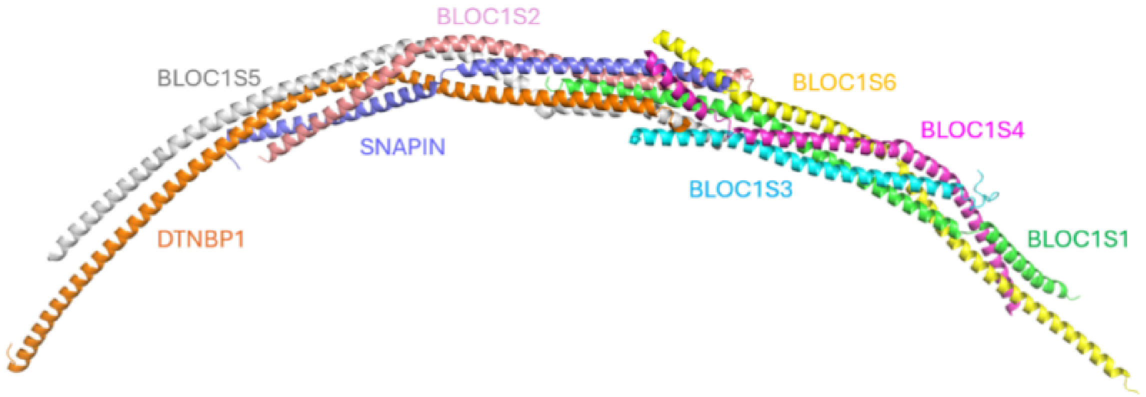


**Figure S40. A model of BLOC-1 complex consisting of BLOC1S5 (grey), DTNBP1 (orange), SNAPIN (periwinkle), BLOC1S2 (pink), BLOC1S6 (yellow), BLOC1S3 (cyan), BLOC1S4 (magenta), and BLOC1S1 (green).**

#### The BLOC-2 complex

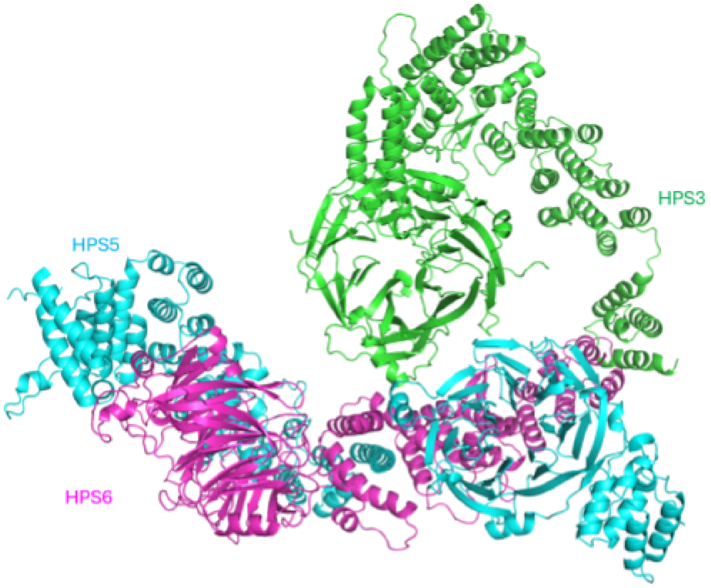


**Figure S41. A model of the BLOC-2 complex made of HPS3 (green), HPS5 (blue), and HPS6 (magenta).**

The BLOC-2 (Biogenesis of Lysosome-related Organelles Complex-2) complex is another key protein complex involved in the biogenesis and function of lysosome-related organelles (LROs) (*180*). The BLOC-2 complex is composed of three subunits: HPS3 (Hermansky-Pudlak Syndrome 3 protein), HPS5, and HPS6 (*183*). The primary function of the BLOC-2 complex is to facilitate the later stages of LRO formation and maturation. BLOC-2 works in conjunction with other protein complexes such as BLOC-1 and BLOC-3 to coordinate the various stages of LRO biogenesis. While BLOC-1 is involved in the early stages, BLOC-2 functions in the later stages, and BLOC-3 assists in the final trafficking steps. We predict pairwise interactions between BLOC-2 complex subunits: HPS3-HPS5 and HPS3-HPS6, allowing us to build a model of the

BLOC-2 complex by AF3 (**Figure S41**). The three subunits adopt the same domain architecture consisting of a N-terminal beta-propeller domain followed by C-terminal alpha-helical repeats. We also detected interactions between BLOC-2 complex subunit HPS5 and BLOC-1 complex subunits SNAPIN and BLOC1S4.

#### The SAMM50 complex

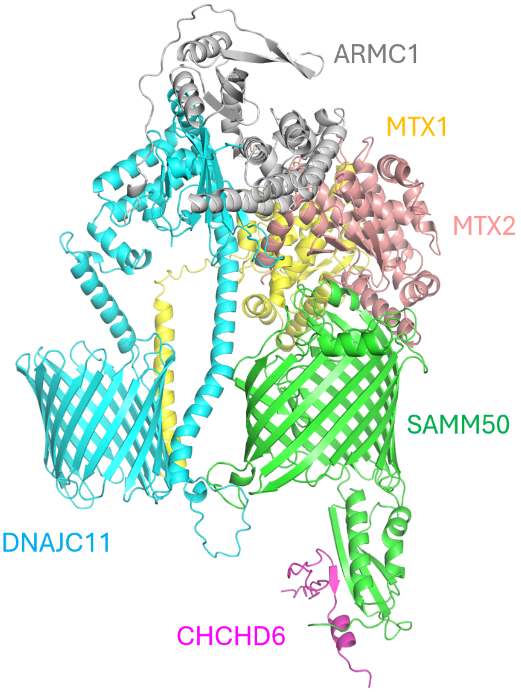


**Figure S42. A model of the SAMM50 complex consisting of SAMM50 (green), CHCHD6 (magenta), DNAJC11 (cyan), ARMC1 (grey), MTX1 (yellow), and MTX2 (salmon).**

The SAMM50 complex, also known as the SAM (Sorting and Assembly Machinery) complex, is a crucial component of the mitochondrial outer membrane (*184*). It plays a significant role in the assembly and maintenance of the mitochondrial outer membrane proteins, particularly the beta-barrel proteins. The SAM complex is essential for mitochondrial biogenesis and proper functioning of the mitochondrial outer membrane. We predict several interactors for SAMM50. DNAJC11, another transmembrane beta-barrel protein (*185*), was predicted to interact with the transmembrane barrel of SAMM50 via a transmembrane helix. SAMM50 is also predicted to interact with MTX2 from the outside of the outer mitochondrial membrane and with CHCHD6 from the intermembrane space. The interaction model of SAMM50 and MTX2 is mediated by the loop regions between the membrane beta strands of SAMM50. The N-terminal alpha-beta-domain of SAMM50 is involved in the interaction between SAMM50 and CHCHD6, which is a subunit of the MICOS complex (*185*). The interaction between SAMM50 and CHCHD6 provides a bridge between the outer membrane and the inner membrane of mitochondria. Additionally, we predict an interaction between DNAJC11 and ARMC1 and an interaction between DNAJC11 and MTX1 (**Figure S42**).

#### The BORC complex

The BORC (BLOC-One Related Complex) is a protein complex that plays a crucial role in the positioning and movement of lysosomes within cells (*186*). The proper positioning of lysosomes is essential for their various cellular functions, including nutrient sensing, signaling, and autophagy. BORC recruits kinesin-1 motor proteins to lysosomes, which move lysosomes along microtubules to the cell periphery. This positioning is crucial for cellular processes like nutrient sensing and signal transduction. By positioning lysosomes at the cell periphery, BORC plays a role in cell migration and invasion (*187*). Lysosomes can release enzymes that degrade extracellular matrix components, facilitating these processes. BORC is involved in autophagy, the process by which cells degrade and recycle their components (*188*). The BORC complex is related to the BLOC-1 complex and they share three subunits: BLOC1S1, BLOC1S2, and SNAPIN. BORC contains five additional subunits: BORCS5, BORCS6, BORCS7, BORCS8, and KXD1. Numerous pairwise interactions were predicted between members of the BORC complex and our model (**Figure S43**) reveals a structure similar to BLOC-1 complex of two coiled-coil four-helical bundles. One four-helical bundle consists of BLOC1S1, BORCS6, BORCS8, and KXD1 and the other consists of BLOC1S2, BORCS5, BORCS7, and SNAPIN.

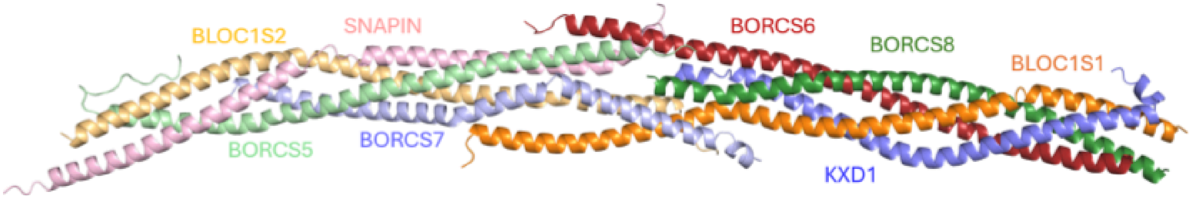


**Figure S43. A model of the BORC complex consisting of BLOC1S2 (yellow), BORCS5 (light green), SNAPIN (pink), BORCS7 (light blue), BORCS6 (dark red), BORCS8 (dark green), BLOC1S1 (dark orange), and KXD1 (dark blue).**

#### The DPM complex

Proper glycosylation is vital for cell-cell interactions, signal transduction, and immune response. The DPM complex, or Dolichol-Phosphate Mannose Synthase complex, is a critical enzyme involved in the process of glycosylation, the process by which sugar molecules are attached to proteins and lipids (*189*). Glycosylation is essential for the proper folding, stability, and function of many proteins. The DPM complex catalyzes the synthesis of Dol-P-Man (a mannose donor in various glycosylation reactions) from GDP-mannose and dolichol phosphate. The DPM complex is composed of multiple subunits: DPM1, DPM2, and DPM3. DPM1 is the catalytic subunit responsible for the synthesis of Dol-P-Man while DPM2 and DPM3 are regulatory subunits that stabilize DPM1 and enhance its activity. DPM2 and DPM3 are transmembrane proteins with two transmembrane segments that anchor the complex to the endoplasmic reticulum (ER) membrane and are necessary for the stability of DPM1. DPM2 is also a subunit of the GPI-GnT complex that interacts with PIGP and PIGY. These proteins are homologous, suggesting a common evolutionary origin of the DPM complex and the GPI-GnT complex. The experimental structure of the DPM complex is not available; however, we provide a high resolution structure model based on our predicted interaction partners (**Figure S44**).

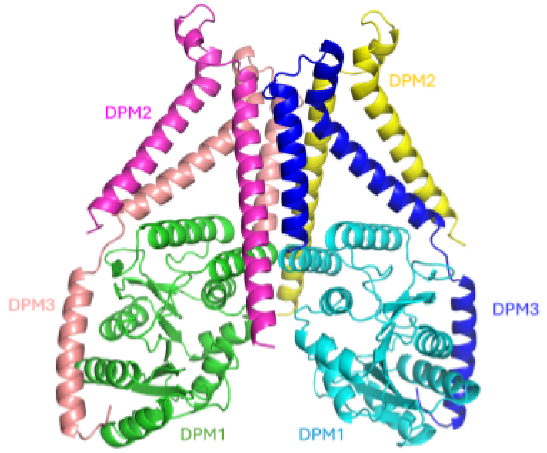


**Figure S44. A model of the DPM complex consisting of components: DPM2 (magenta), DPM3 (pink), DPM1 (green) and the homodimeric copies: DPM1' (cyan), DPM3' (blue), and DPM2' (yellow).**

#### The URI1 complex

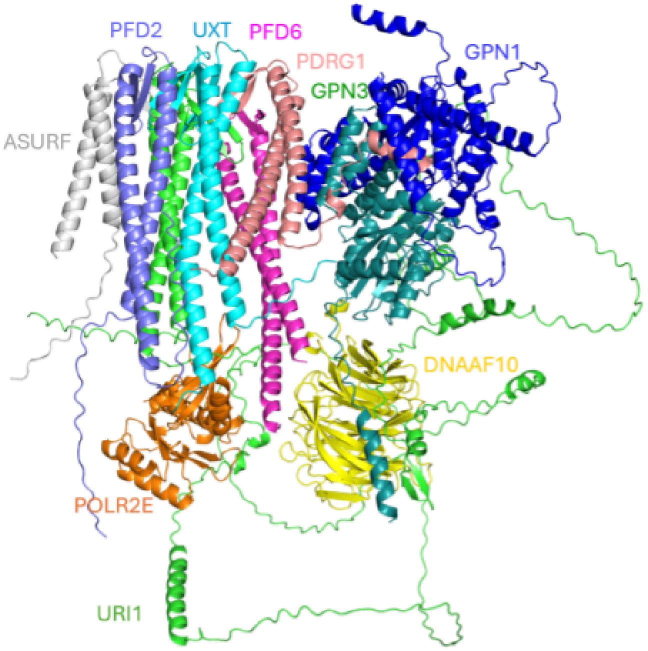


**Figure S45. A model of theURI1 complex consisting of ASURF (grey), PFD2 (periwinkle), UXT (cyan), PFD6 (magenta), PDRG1 (pink), GPN3 (aquamarine), GPN1 (blue), POLR2E (orange), DNAAF10 (yellow), and URI1 (green).**

The PAQosome (Particle for Arrangement of Quaternary structure) is a large multi-protein complex that acts as a cochaperone for the biogenesis and proper assembly of various multiprotein complexes within the cell (*190*) The PAQosome is composed of multiple proteins, primarily from the R2TP complex and Prefoldin-like proteins forming a URI1 prefoldin co-chaperone complex (*191*). The URI1 prefoldin co-chaperone complex is a specialized molecular chaperone complex that plays a crucial role in the proper folding and assembly of nascent polypeptides and protein complexes. URI1 stands for "Unconventional prefoldin RPB5 Interactor 1," reflecting its unique features and functions compared to classical prefoldin complexes. The URI1 complex is composed of multiple subunits, some of which are shared with the classical prefoldin complex, as well as additional unique components. The key subunits of the URI1 complex include: URI1, UXT, PDRG1, ADSURF, PFDN2, and PFDN6. Of these components, PFDN2 and PFDN6 are shared with the classical prefoldin complex.

We predict multiple interactions between members of the URI1 complex. The URI1 subunit which contains a long C-terminal disordered region is distinct from the other primarily globular URI1 subunits. Multiple interactions with other proteins were predicted to interact with the disordered C-terminal region of URI1, including DNAA10F, POLR2E, GPS1, and GPS3, reflecting the many functional associations. We built a multisubunit model of the URI1 complex and its interaction partners (**Figure S45**). All six subunits of the URI1 complex adopt the prefoldin fold with two alpha-helices forming a helical hairpin and one or two beta-hairpins inserted in between the helices. The beta-hairpins form two intertwined beta-barrels in the complex and the alpha-helices interact with each other to form a ring-like core structure.

149. K. G. Guruharsha, J.-F. Rual, B. Zhai, J. Mintseris, P. Vaidya, N. Vaidya, C. Beekman, C. Wong, D. Y. Rhee, O. Cenaj, E. McKillip, S. Shah, M. Stapleton, K. H. Wan, C. Yu, B. Parsa, J. W. Carlson, X. Chen, B. Kapadia, K. VijayRaghavan, S. P. Gygi, S. E. Celniker, R. A. Obar, S. Artavanis-Tsakonas, A protein complex network of Drosophila

melanogaster. *Cell* **147**, 690–703 (2011).

159. B. Bausch, F. Schiavi, Y. Ni, J. Welander, A. Patocs, J. Ngeow, U. Wellner, A. Malinoc, E. Taschin, G. Barbon, V. Lanza, P. Söderkvist, A. Stenman, C. Larsson, F. Svahn, J.-L. Chen, J. Marquard, M. Fraenkel, M. A. Walter, M. Peczkowska, A. Prejbisz, B. Jarzab, K. Hasse-Lazar, S. Petersenn, L. C. Moeller, A. Meyer, N. Reisch, A. Trupka, C. Brase, M. Galiano, S. F. Preuss, P. Kwok, N. Lendvai, G. Berisha, Ö. Makay, C. C. Boedeker, G. Weryha, K. Racz, A. Januszewicz, M. K. Walz, O. Gimm, G. Opocher, C. Eng, H. P. H. Neumann, European-American-Asian Pheochromocytoma-Paraganglioma Registry Study Group, Clinical Characterization of the Pheochromocytoma and Paraganglioma Susceptibility Genes SDHA, TMEM127, MAX, and SDHAF2 for Gene-Informed Prevention. *JAMA Oncol* **3**, 1204–1212 (2017).

160. Y. Deng, Y. Qin, S. Srikantan, A. Luo, Z.-M. Cheng, S. K. Flores, K. S. Vogel, E. Wang, P.

L. M. Dahia, The TMEM127 human tumor suppressor is a component of the mTORC1 lysosomal nutrient-sensing complex. *Hum. Mol. Genet.* **27**, 1794–1808 (2018).

161. J. Brodeur, C. Thériault, M. Lessard-Beaudoin, A. Marcil, S. Dahan, C. Lavoie, LDLR-related protein 10 (LRP10) regulates amyloid precursor protein (APP) trafficking and processing: evidence for a role in Alzheimer's disease. *Mol. Neurodegener.* **7**, 31 (2012).
162. M. Quadri, W. Mandemakers, M. M. Grochowska, R. Masius, H. Geut, E. Fabrizio, G. J. Breedveld, D. Kuipers, M. Minneboo, L. J. M. Vergouw, A. Carreras Mascaro, E. Yonova-Doing, E. Simons, T. Zhao, A. B. Di Fonzo, H.-C. Chang, P. Parchi, M. Melis, L. Correia Guedes, C. Criscuolo, A. Thomas, R. W. W. Brouwer, D. Heijsman, A. M. T. Ingrassia, G. Calandra Buonaura, J. P. Rood, S. Capellari, A. J. Rozemuller, M. Sarchioto, H. Fen Chien, N. Vanacore, S. Olgiati, Y.-H. Wu-Chou, T.-H. Yeh, A. J. W. Boon, S. E. Hoogers, M. Ghazvini, A. S. IJpma, W. F. J. van IJcken, M. Onofrj, P. Barone, D. J. Nicholl, A. Puschmann, M. De Mari, A. J. Kievit, E. Barbosa, G. De Michele, D. Majoor-Krakauer, J. C. van Swieten, F. J. de Jong, J. J. Ferreira, G. Cossu, C.-S. Lu, G. Meco, P. Cortelli, W. D. J. van de Berg, V. Bonifati, International Parkinsonism Genetics Network, LRP10 genetic variants in familial Parkinson's disease and dementia with Lewy bodies: a genome-wide linkage and sequencing study. *Lancet Neurol.* **17**, 597–608 (2018).
163. A. M. West, S. C. Rosenberg, S. N. Ur, M. K. Lehmer, Q. Ye, G. Hagemann, I. Caballero, I. Usón, A. J. MacQueen, F. Herzog, K. D. Corbett, A conserved filamentous assembly underlies the structure of the meiotic chromosome axis. *Elife* **8** (2019).
164. Y. Fujiwara, Y. Horisawa-Takada, E. Inoue, N. Tani, H. Shibuya, S. Fujimura, R. Kariyazono, T. Sakata, K. Ohta, K. Araki, Y. Okada, K.-I. Ishiguro, Meiotic cohesins mediate initial loading of HORMAD1 to the chromosomes and coordinate SC formation during meiotic prophase. *PLoS Genet.* **16**, e1009048 (2020).
165. L. Osterloh, B. von Eyss, F. Schmit, L. Rein, D. Hübner, B. Samans, S. Hauser, S. Gaubatz, The human synMuv-like protein LIN-9 is required for transcription of G2/M genes and for entry into mitosis. *EMBO J.* **26**, 144–157 (2007).
166. A. Oji, A. Isotani, Y. Fujihara, J. M. Castaneda, S. Oura, M. Ikawa, Tesmin, Metallothionein-Like 5, is Required for Spermatogenesis in Mice†. *Biol. Reprod.* **102**, 975–983 (2020).
167. X. Zhang, M. Li, X. Jiang, H. Ma, S. Fan, Y. Li, C. Yu, J. Xu, R. Khan, H. Jiang, Q. Shi, Nuclear translocation of MTL5 from cytoplasm requires its direct interaction with LIN9 and is essential for male meiosis and fertility. *PLoS Genet.* **17**, e1009753 (2021).
168. K. S. Makarova, V. Tobiasson, Y. I. Wolf, Z. Lu, Y. Liu, S. Zhang, M. Krupovic, M. Li, E. V. Koonin, Diversity, origin, and evolution of the ESCRT systems. *MBio* **15**, e0033524 (2024).
169. L. Wunderley, K. Brownhill, F. Stefani, L. Tabernero, P. Woodman, The molecular basis for selective assembly of the UBAP1-containing endosome-specific ESCRT-I complex. *J. Cell Sci.* **127**, 663–672 (2014).
170. M. Tagaya, K. Arasaki, H. Inoue, H. Kimura, Moonlighting functions of the NRZ (mammalian Dsl1) complex. *Front Cell Dev Biol* **2**, 25 (2014).
171. G. J. P. L. Kops, Y. Kim, B. A. A. Weaver, Y. Mao, I. McLeod, J. R. Yates 3rd, M. Tagaya, D. W. Cleveland, ZW10 links mitotic checkpoint signaling to the structural kinetochore. *J. Cell Biol.* **169**, 49–60 (2005).
172. E. Ozawa, Y. Mizuno, Y. Hagiwara, T. Sasaoka, M. Yoshida, Molecular and cell biology of the sarcoglycan complex. *Muscle Nerve* **32**, 563–576 (2005).
173. L. E. Lim, K. P. Campbell, The sarcoglycan complex in limb-girdle muscular dystrophy. *Curr. Opin. Neurol.* **11**, 443–452 (1998).
